# A statistical framework for cross-tissue transcriptome-wide association analysis

**DOI:** 10.1101/286013

**Authors:** Yiming Hu, Mo Li, Qiongshi Lu, Haoyi Weng, Jiawei Wang, Seyedeh M. Zekavat, Zhaolong Yu, Boyang Li, Jianlei Gu, Sydney Muchnik, Yu Shi, Brian W. Kunkle, Shubhabrata Mukherjee, Pradeep Natarajan, Adam Naj, Amanda Kuzma, Yi Zhao, Paul K. Crane, Alzheimer’s Disease Genetics Consortium, Hui Lu, Hongyu Zhao

## Abstract

Transcriptome-wide association analysis is a powerful approach to studying the genetic architecture of complex traits. A key component of this approach is to build a model to impute gene expression levels from genotypes using samples with matched genotypes and gene expression data in a given tissue. However, it is challenging to develop robust and accurate imputation models with a limited sample size for any single tissue. Here, we first introduce a multi-task learning method to jointly impute gene expression in 44 human tissues. Compared with single-tissue methods, our approach achieved an average 39% improvement in imputation accuracy and generated effective imputation models for an average 120% more genes. We then describe a summary statistic-based testing framework that combines multiple single-tissue associations into a powerful metric to quantify the overall gene-trait association. We applied our method, called UTMOST, to multiple genome wide association results and demonstrate its advantages over single-tissue strategies.

## Introduction

Genome-wide association studies (GWAS) have successfully identified numerous single-nucleotide polymorphisms (SNPs) associated with complex human traits and diseases. Despite these successes, significant problems remain in statistical power and biological interpretation of GWAS results^1,2^. In particular, the complex architecture of linkage disequilibrium (LD) and context-dependent regulatory machinery in the genome hinder our ability to accurately identify disease genes from GWAS, thereby raising challenges in downstream functional validation and therapeutics development. Recently, large-scale consortia, such as the Genotype-Tissue Expression (GTEx) project^3,4^, have generated matched genotype and expression data for various human tissues. These rich data sets have provided great insights into the mechanisms of cross-tissue transcriptional regulation and accelerated discoveries for expression quantitative trait loci (eQTL)^4–7^. In addition, integrating eQTL information in genetic association analysis has become an effective way to bridge SNPs, genes, and complex traits. Many methods have been developed to co-localize eQTL with loci identified in GWAS to identify candidate risk genes for complex traits^8–13^. Two recent studies addressed this issue through an innovative approach that is sometimes referred to as transcriptome-wide association analysis. First, based on an externally-trained imputation model, gene expression is imputed using genotype information in GWAS samples. Next, gene-level association is assessed between imputed gene expression and the trait of interest^14,15^. These methods have gained popularity in the past two years due to their capability to effectively utilize signals from multiple eQTL with moderate effects and to reduce the impact of reverse causality in expression-trait association analysis. The applications of these methods have led to novel insights into the genetic basis of many diseases and traits^16–18^.

Despite these successes, existing methods have several limitations. First, due to the tissue-dependent nature of transcription regulation, existing methods train separate imputation models for different tissues. This practice ignores the similarity in transcription regulation across tissues, thereby limiting the effective sample sizes for tissues that are difficult to acquire. Second, a hypothesis-free search across genes and tissues increases the burden of multiple testing and thus reduces statistical power. Pinpointing a subset of tissues based on prior knowledge may resolve this issue to some extent. However, for many complex traits, biologically relevant tissues are unknown. Further, reports have shown that eQTL with large effects tend to regulate gene expression in multiple tissues^4^. Genetic correlation analysis has also suggested substantial sharing of local expression regulation across tissues^19^. This would inevitably result in statistically significant associations in tissues irrelevant to the trait of interest, a phenomenon that has been extensively discussed recently^20^. Jointly analyzing data from multiple genetically-correlated tissues has the potential to resolve these issues. It has been demonstrated that multi-trait analysis could improve accuracy of genetic risk prediction^21–23^. Multi-tissue modeling has also been shown to improve the statistical power in eQTL discovery^24–27^ and gene network studies^28^. In this work, we demonstrate that a cross-tissue strategy could also improve transcriptome-wide association analysis.

We introduce UTMOST (Unified Test for MOlecular SignaTures), a principled method to perform cross-tissue expression imputation and gene-level association analysis. We demonstrate its performance through internal and external imputation validation, simulation studies, analyses of 50 complex traits, a case-study on low-density lipoprotein cholesterol (LDL-C), and a multi-stage association study for late-onset Alzheimer’s disease (LOAD). We show that UTMOST substantially improves the accuracy of expression imputation in all available tissues. In the downstream association analysis, UTMOST provides a powerful metric that summarizes gene-level associations across tissues and can be extended to integrate various molecular phenotypes.

## Results

### Model overview

The UTMOST framework consists of three main stages (Figure 1). First, for each gene in the genome, we train a cross-tissue expression imputation model using the genotype information and matched expression data from 44 tissues in GTEx. Next, we test associations between the trait of interest and imputed expression in each tissue. Lastly, a cross-tissue test is performed for each gene to summarize single-tissue association statistics into a powerful metric that quantifies the overall gene-trait association. Here, we briefly introduce the UTMOST framework. All the statistical details are discussed in the **Online Methods**.

**Figure 1.**
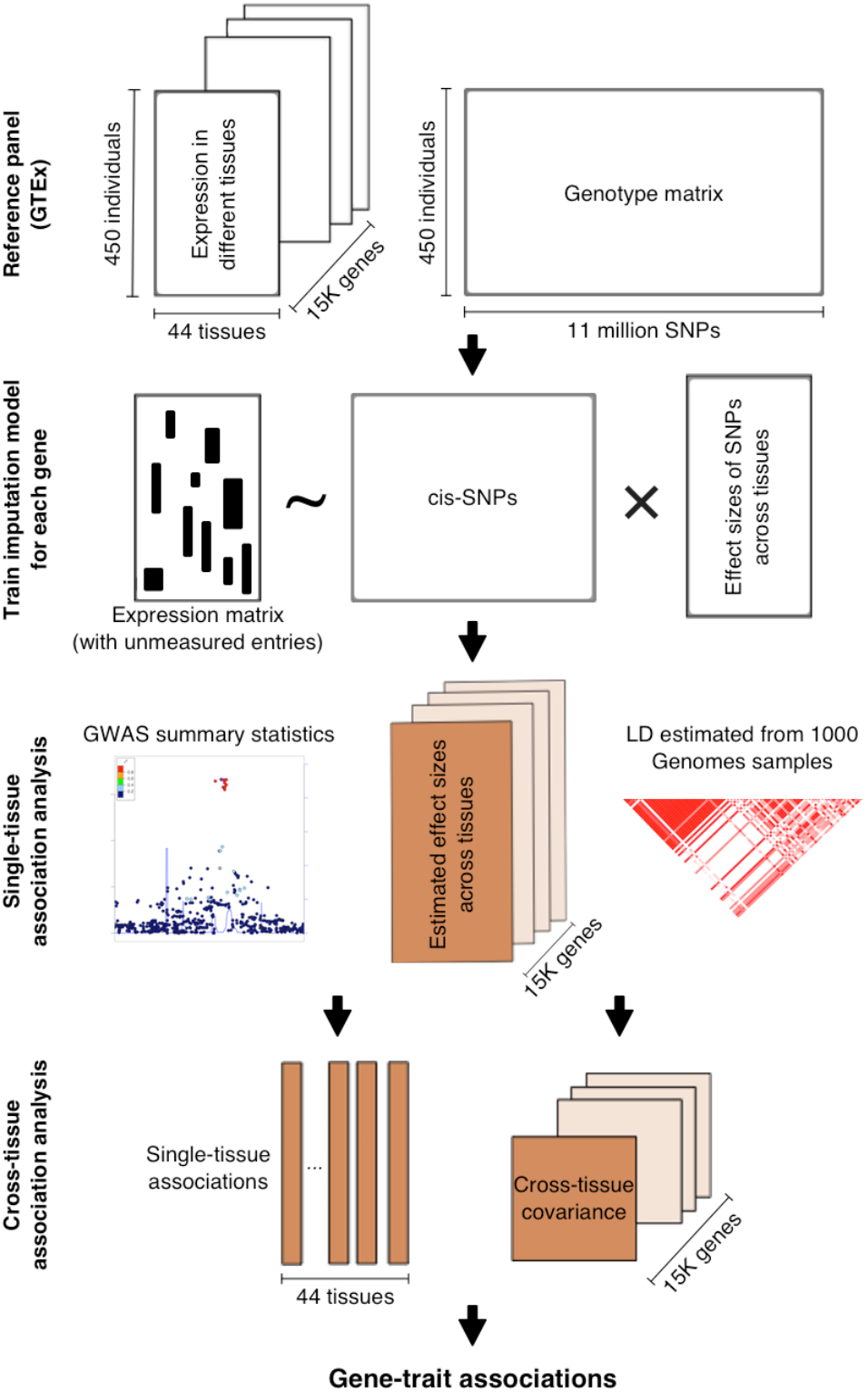
UTMOST workflow. Gray and brown boxes denote input data and computed outcomes, respectively.

We formulate cross-tissue expression imputation as a penalized multivariate regression problem:

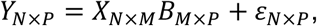

where *N, M*, and *P* denote the sample size in the training data, the number of SNPs in the imputation model, and the total number of tissues, respectively. As only a subset of tissues was collected from each individual, expression data in matrix *Y* were incomplete and sample sizes for different tissues were unbalanced. We estimate *B* by minimizing the squared loss function with a lasso penalty on the columns (within-tissue effects) and a group-lasso penalty on the rows (cross-tissue effects) (**Online Methods**).

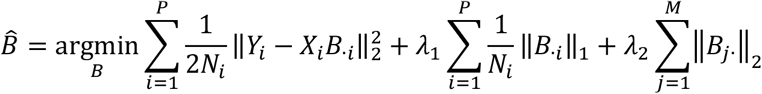

where *Y_i_,X_i_* and *N_i_* denote the observed expressions, genotypes, and sample size of the *i*th tissue, respectively. Parameters *λ*_1_ and *λ*_2_ are tuned through cross-validation. Our cross-tissue imputation model does not assume eQTL to have the same effect direction across tissues. Instead, UTMOST uses a group LASSO ^29^ penalty term the framework to encourage the presence of cross-tissue eQTL and improve the estimation of their effects.

In the second stage, we test the associations between the trait of interest and imputed gene expression in each tissue. We denote imputed gene expression in the *i*th tissue as 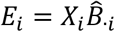 and test associations via a univariate regression model:

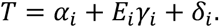

The z-scores for gene-trait associations in the *i*th tissue can be denoted as

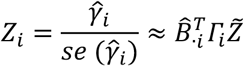

where 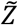 denotes the SNP-trait z-scores and *Γ_i_* is a diagonal matrix whose *j*th diagonal element denotes the ratio between the standard deviation of the *j*th SNP and that of imputed expression in the *i*th tissue (**Online Methods**). When there is no SNP-trait association, 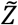 follows a multivariate normal distribution *N*(0,*D*), where *D* is the LD matrix for SNPs. The covariance matrix of *Z* = (*Z*_1_,*Z*_2_,…,*Z_P_*)*^T^* can be calculated as

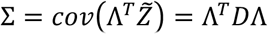

where 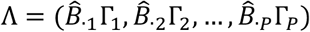

Finally, we combine single-tissue gene-trait association results using a generalized Berk-Jones (GBJ) test, which takes the covariance among single-tissue test statistics into account^30^. We note that this framework allows gene-trait associations to have different directions across tissues. Details on the GBJ statistic and p-value calculation are discussed in the **Online Methods**.

### Cross-tissue expression imputation accuracy

We first evaluated the accuracy of cross-tissue expression imputation through five-fold cross-validation. We used an elastic net model (i.e. the model used in PrediXcan^14^) trained in each tissue separately as the benchmark for prediction without leveraging cross-tissue information. We used squared Pearson correlation (i.e. *R^2^*) between the observed and predicted gene expression levels to quantify imputation accuracy. Cross-tissue imputation achieved higher imputation accuracy in all 44 tissues (Figure 2a). On average, imputation accuracy was improved by 38.6% across tissues (Figure 2b). The improvement was particularly high in tissues with low sample sizes in GTEx (N < 150; an average of 47.4% improvement). Analysis based on Spearman correlation also showed consistent results (**Supplementary Figure 1**). Next, we calculated the proportion of genes with increased imputation accuracy. In all 44 tissues, substantially more genes showed improved imputation performance (**Supplementary Table 1**). Using a false discovery rate (FDR) cutoff of 0.05 as the significance threshold, our cross-tissue method achieved 120% more significantly predicted genes across tissues. Among tissues with low sample sizes, the improvement percentage rose even further to 175% (Figure 2c). Furthermore, we compared our method with the Bayesian Sparse Linear Mixed-effects Model (BsLmM^31^), the imputation method used in TWAS^15^. Similarly, UTMOST achieved higher imputation accuracy in all 44 tissues (**Supplementary Figure 2**). On average, imputation accuracy improved 20.3% across tissues.

**Figure 2.**
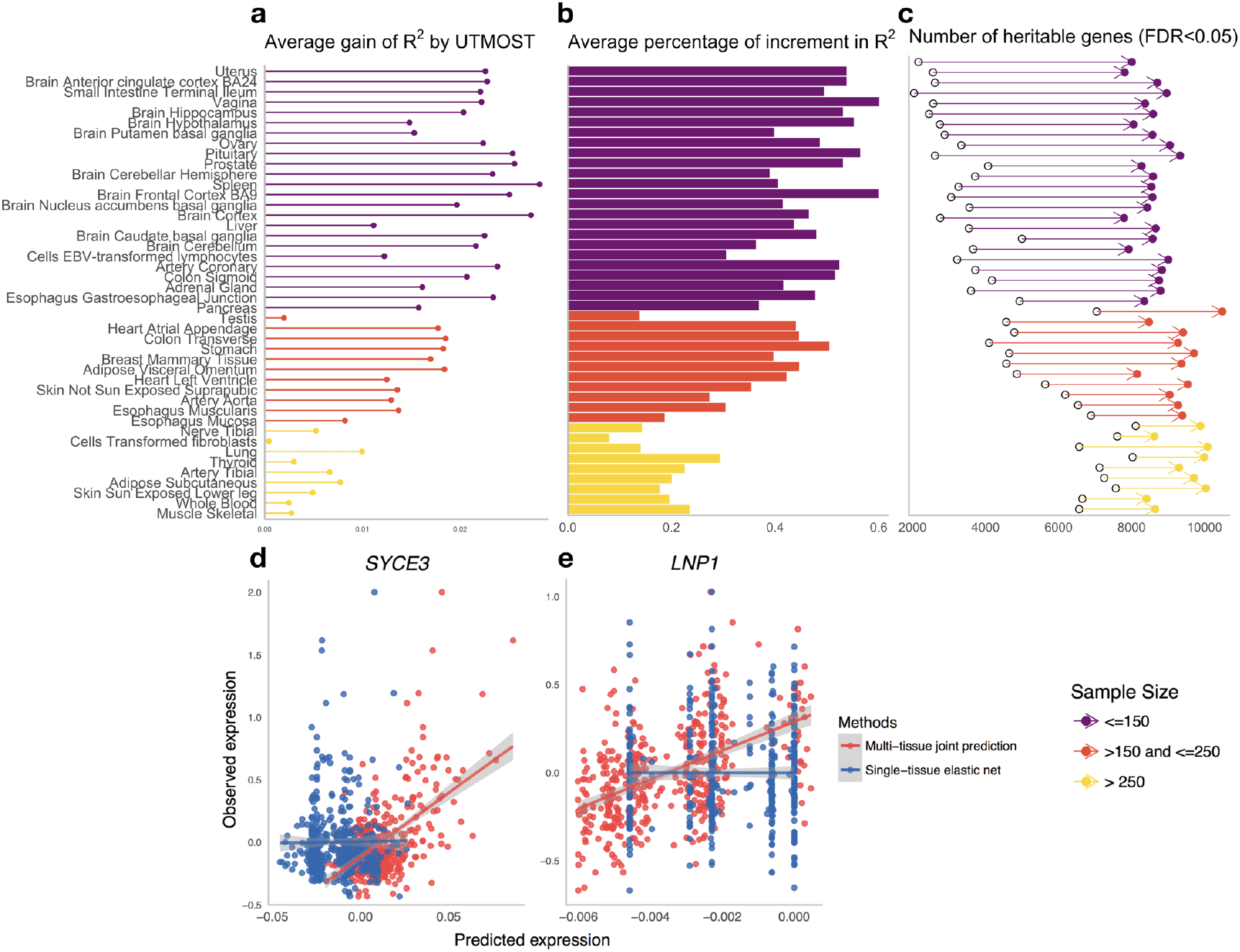
Improvement in gene expression imputation accuracy. Compared to single-tissue elastic net, UTMOST showed substantially higher **(a)** average increment in *R^2^* across genes and **(b)** relative improvement (i.e. percentage of increment in *R^2^*) in imputation accuracy. **(c)** UTMOST identified more imputed genes, especially in tissues that have smaller sample sizes in GTEx. Sample sizes of 44 GTEx tissues are listed in **Supplementary Table 1**, predictability tested by F-test with d.f. 1 and n - 2. Panels **(d-e)** show the imputation improvement in two specific examples in whole blood tissue, shaded region represents the 95% confidence band.

Next, we performed external validation using two independent datasets. We first used our imputation model for whole blood in GTEx to predict gene expression levels in GEUVADIS lymphoblastoid cell lines (LCLs)^32^ (**Online Methods**). The imputation accuracy quantified as R^2^ showed substantial departure from the expected distribution under the null (i.e. expression and SNPs are independent), which demonstrates the generalizability of cross-tissue imputation (**Supplementary Figures 3-4**). Compared to single-tissue elastic net, cross-tissue imputation achieved significantly higher prediction accuracy in different quantiles (*P* = 3.43 × 10^-7^; Kolmogorov-Smirnov test), which is consistent with our findings from cross-validation. Two examples of well-predicted genes are illustrated in Figure 2d-e, showing improved concordance between observed (gene expressions adjusted for potential confounding effects; **Online Methods**) and predicted expression values via cross-tissue imputation. Analysis on CommonMind consortium data^33^ showed similar results (Online Methods, **Supplementary Figure 5-6**).

### Cross-tissue association test

Another key advancement in the UTMOST framework is a novel gene-level association test that combines statistical evidence across multiple tissues. We performed simulation studies using samples from the Genetic Epidemiology Research Study on Adult Health and Aging (GERA; N = 12,637) to assess the association test’s type-I error rate and statistical power in a variety of settings (**Online Methods**). We did not observe inflation in the type-I error rate in two different simulation studies (**Supplementary Table 2-3**). We observed a substantial improvement in statistical power of the multi-tissue joint test when gene expressions in multiple tissues were causally related to the trait. The improvement was also consistent under different simulated genetic architectures (Figure 3). When the trait was affected by expression in only one tissue, statistical power of the joint test was comparable to that of a single-tissue test in the causal tissue. Compared to the naïve test that combines results across tissues while applying an additional Bonferroni correction, our joint test was consistently more powerful (improvement ranged from 15.3% to 24.1%).

**Figure 3.**
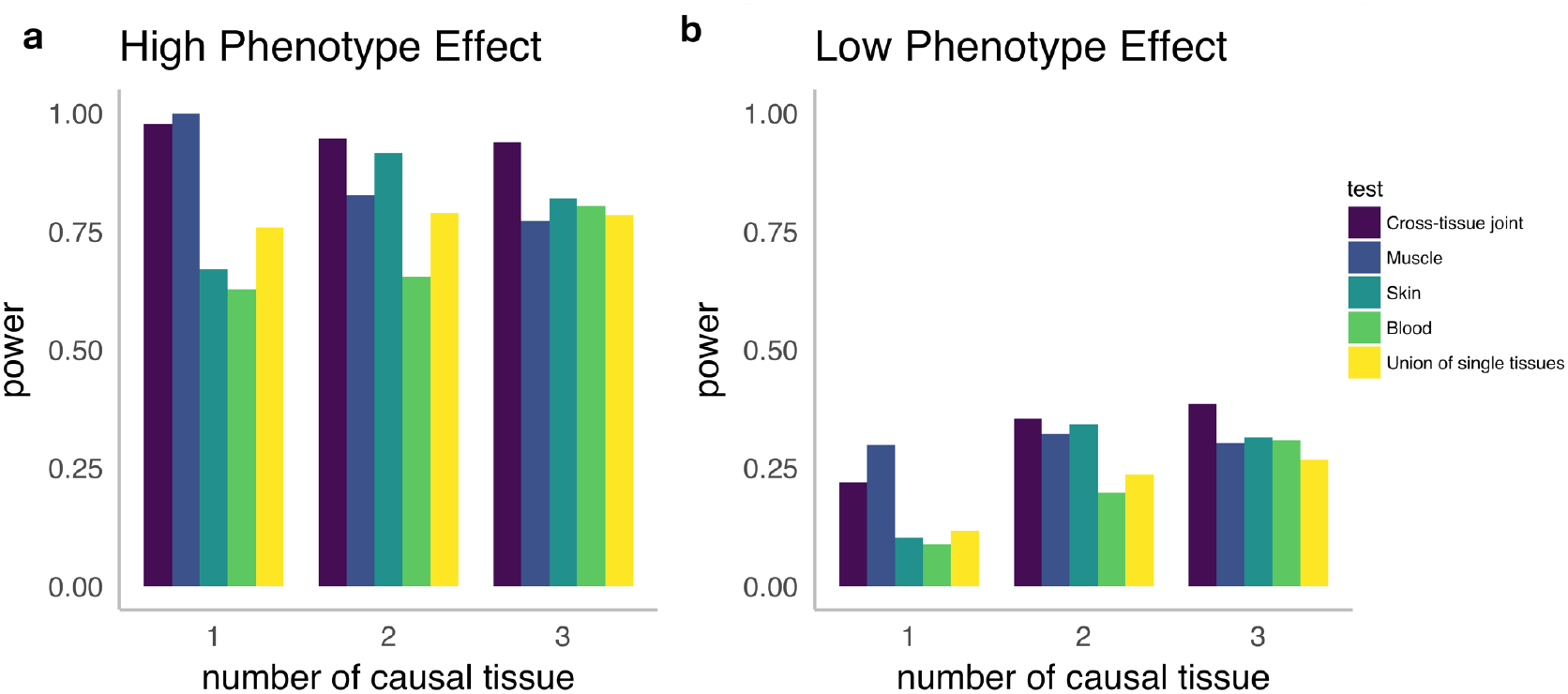
Cross-tissue analysis improves statistical power. We compared the statistical power of UTMOST, a single-tissue association test, and a simple union of findings from single-tissue analysis with various disease architectures. Left/right panels represent the cases that genes explain 1%/0.1% of trait variance in total (denoted as high/low phenotypic effects). Muscle is the only causal tissue in setting 1. Both muscle and skin are causal tissues in setting 2. All three tissues are causal in setting 3.

### UTMOST identifies more associations in relevant tissues

To evaluate the performance of single-tissue association test based on cross-tissue expression imputation, we applied UTMOST to the summary statistics from 50 GWAS (*N_total_*≈4.5 million without adjusting for sample overlap across studies; **Supplementary Table 4**) and compared the results with those of PrediXcan^14^ and TWAS^15^. To identify tissue types that are biologically relevant to these complex traits, we applied LD score regression^34^ to these datasets and partitioned heritability by tissue-specific functional genome predicted by GenoSkyline-Plus annotations^35^. Tissue-trait relevance was ranked based on enrichment p-values (**Methods**). Compared to PrediXcan and TWAS, UTMOST identified substantially more associations in the most relevant tissue for each analyzed trait, showing 69.2% improvement compared to PrediXcan *(P* = 8.79 × 10^-5^; paired Wilcoxon rank test) and 188% improvement compared to TWAS (*P* = 7.39 × 10^-8^, Figure 4). Such improvement was consistently observed across traits (**Supplementary Table 5**). In contrast, for other tissues, UTMOST identified similar number of genes and showed no significant difference compared with PrediXcan (*P* = 0.52). Comparing tissues that were most and least enriched for trait heritability, UTMOST identified significantly more associations in tissues strongly enriched for trait heritability than in tissues with the least enrichment (*P* = 0.016) while the contrast was not significant based on PrediXcan (*P* = 0.192) or TWAS (*P* = 0.085). Finally, we applied the cross-tissue joint test to these traits and compared the number of significant genes with the combined results from 44 UTMOST single-tissue tests. UTMOST joint test identified more associations than single-tissue tests in 43 out of 50 traits (*P* = 1.74 × 10^-8^; Wilcox rank test; **Supplementary Figure 7**), showing improved statistical power in cross-tissue analysis.

**Figure 4.**
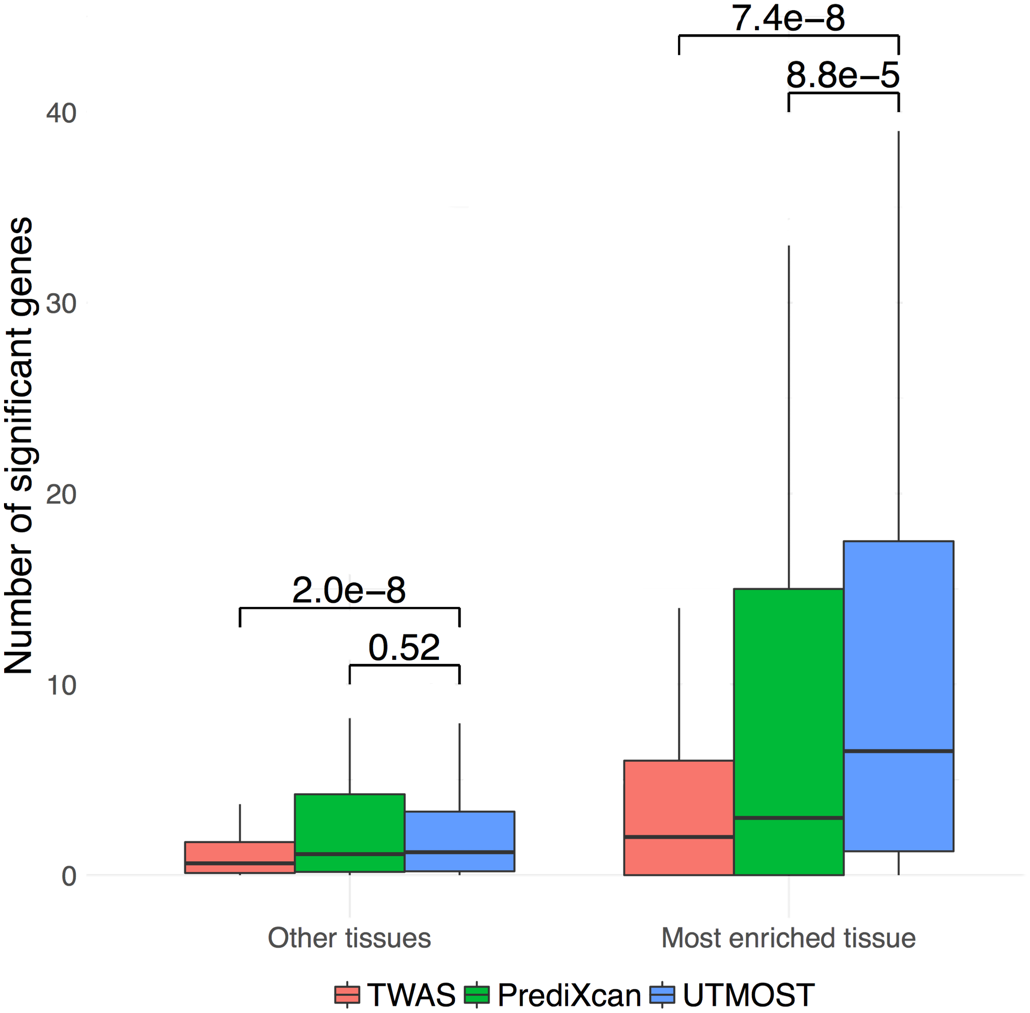
UTMOST identified more associations in biologically relevant tissues for 50 complex traits. Boxes on the left show the number of genes identified in all other tissues. Boxes on the right show the number of genes identified in the most relevant tissue for each trait. In each box, the two horizontal borders represent the upper and lower quartiles, solid line in the middle represent median. The highest and lowest points indicate the maxima and minima. P-values were calculated via one-sided paired Wilcoxon rank tests (n = 50).

### Integrating external QTL resource

We applied UTMOST to the meta-analysis summary data of LDL-C from the Global Lipids Genetics Consortium (N = 173,082)^36^. Results based on four different analytical strategies, i.e. single-tissue test using liver tissue in GTEx (N = 97), single-tissue test using liver eQTL from STARNET^37^ (N = 522), cross-tissue joint test combining 44 GTEx tissues, and cross-tissue joint test combining 44 GTEx tissues and the liver eQTL from STARNET, were compared. We identified 57, 58, 185, and 203 significant genes in the four sets of analyses, respectively (Figure 5a).

Among the identified genes in cross-tissue joint test of 44 GTEx tissues and STARNET-liver, *SORT1* had the most significant association (*P* = 3.4 × 10^-15^). *SORT1* is known to causally mediate LDL-C levels, even though the GWAS association signal at this locus is clustered around *CELSR2*^38,39^. Of note, not only was liver not implicated as the relevant tissue for *SORT1* in the association analysis, association signal at *SORT1* was completely absent in the single tissue test based on GTEx-liver due to its low imputation quality (FDR = 0.064). Limited sample size of liver tissue in GTEx (N = 97) restrained the imputation performance of SORT1, and consequently reduced the statistical power in association test. On the other hand, UTMOST successfully recovered the association signal at *SORT1* (*P* = 3.4 × 10^-15^). Additionally, UTMOST cross-tissue association test is flexible in incorporating external QTL resources along with GTEx data (**Online Methods**). Through integrating single-tissue associations in all 44 GTEx tissues and a large external liver dataset (STARNET; N = 522), we successfully recovered the association of *SORT1* (Figure 5b). Furthermore, we performed pair-wise conditional analyses between *SORT1* and other significant genes at the *SORT1* locus, and found that *SORT1* remained statistically significant in all analyses, showing that its association signal is not shadowed by other genes (**Supplementary Table 6**). Further, when correlations between gene expression were moderate, *SORT1* was more significant than all other tested genes in conditional analysis. Even when correlation was substantial (e.g. *CELSR2* and *PSRC1* both had correlation = 0.9 with *SORT1* in STARNET), *SORT1* remained statistically significant. We compared association based on STARNET only and found that *SORT1* is not the top signal in the locus in single-tissue analysis and cross-tissue approach does not increase the false-positive rate (**Supplementary Note**). These results suggest that integrative analysis of transcriptomic data from multiple tissues and multiple QTL resources can effectively increase statistical power in gene-level association mapping. UTMOST is a flexible framework and is not limited to GTEx tissues only. Integrating relevant external QTL studies via UTMOST may further improve downstream association analysis.

**Figure 5.**
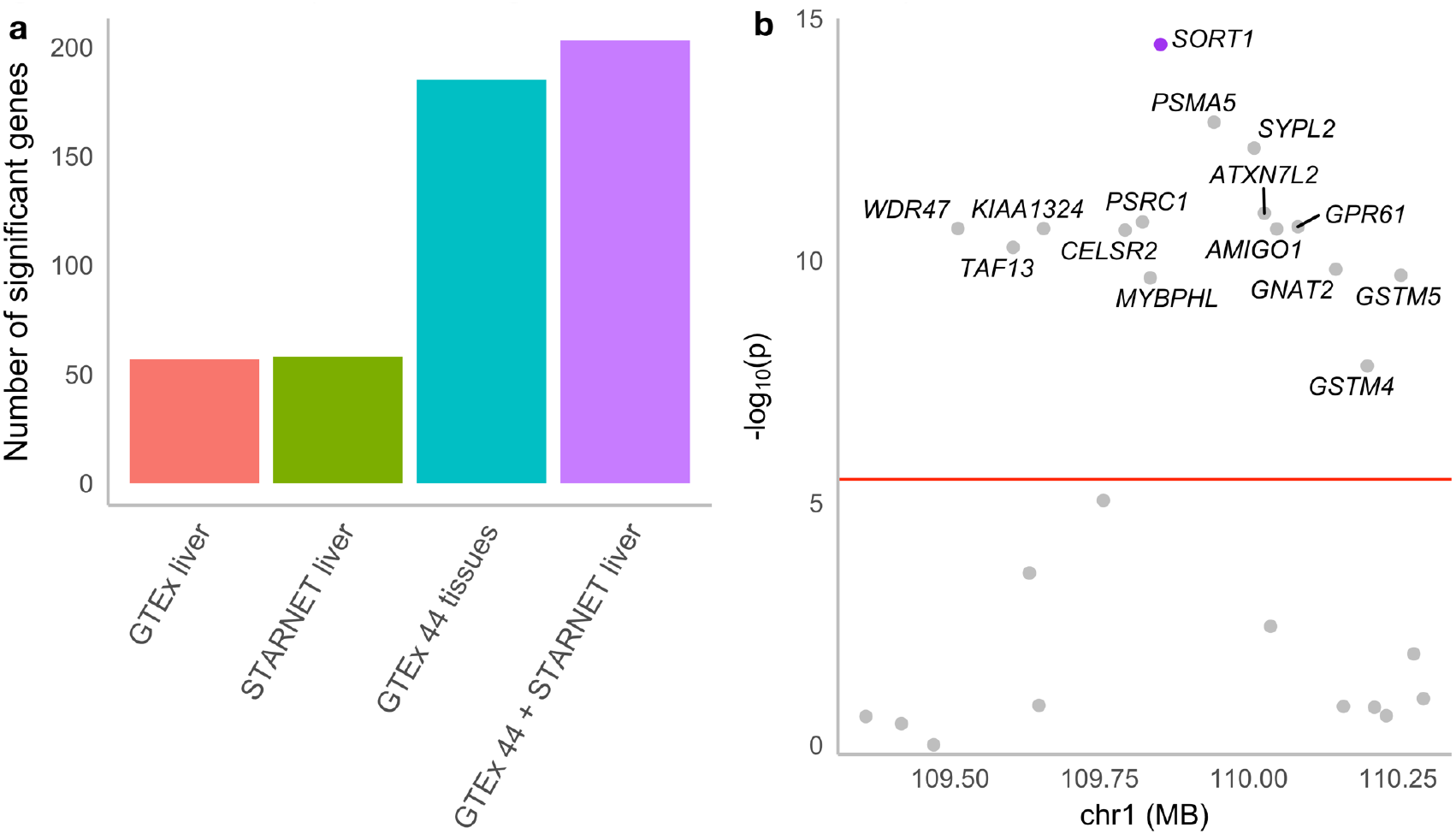
Multi-tissue analysis identifies more associations for LDL cholesterol. **(a)** Number of significant genes identified in four sets of analyses. (z-score test for single-tissue and generalized Berk-Jones for cross-tissue test, Bonferroni-corrected thresholds were used, i.e. 4.49 × 10^-6^, 8.39 × 10^-6^, 3.31 × 10^-6^ and 3.31 × 10^-6^) **(b)** Associations at the *SORT1* locus, values on the x-axis were based on the transcription start site of each gene. The horizontal line indicates the Bonferroni-corrected genome-wide significance threshold (n = 173,082, generalized Berk-Jones test).

### UTMOST identifies novel risk genes for Alzheimer’s disease

Finally, to demonstrate UTMOST’s effectiveness in real association studies, we performed a multi-stage gene-level association study for LOAD. In the discovery stage, we applied UTMOST to the stage-I GWAS summary statistics from the International Genomics of Alzheimer’s Project^40^ (IGAP; N =54,162). Multiple recent studies have suggested that functional DNA regions in liver and myeloid cells are strongly enriched for LOAD heritability^35,41,42^. It has also been suggested that alternative splicing may be a mechanism for many risk loci of LOAD^43^. Therefore, in addition to 44 tissues from GTEx, we also incorporated liver eQTL from STARNET and both eQTL and splicing (s)QTL data in three immune cell types (i.e. CD14+ monocytes, CD16+ neutrophils, and naive CD4+ T cells) from the BLUEPRINT^44^ consortium in our analysis (**Online Methods**). Single-tissue association tests were performed and then combined using the GBJ test. In total, our cross-tissue analysis identified 68 genome-wide significant genes in the discovery stage (**Supplementary Table 7, Supplementary Figure 8**).

Next, we replicated our findings in two independent datasets: using GWAS summary statistics based on samples in the Alzheimer’s Disease Genetics Consortium (ADGC) that were not used in the IGAP stage-I analysis (N = 7,050), and summary statistics from the genome-wide association study by proxy^45^ (GWAX; N = 114,564). Despite the moderate sample size in the ADGC dataset and the ‘proxy’ LOAD phenotype based on family history in GWAX analysis, replication rate was high (**Supplementary Table 7**). Seventeen and 15 out of 68 genes were successfully replicated under the Bonferroni-corrected significance threshold in ADGC and GWAX, respectively. The numbers of replicated genes rose to 41 and 30 under a relaxed p-value cutoff of 0.05. Twenty-two out of 68 genes had p-values below 0.05 in both replication datasets. We then combined p-values from all three analyses via Fisher’s method. A total of 69 genes, including 12 genes that were not significant in the discovery stage, reached genome-wide significance in the meta-analysis (Figure 6, **Supplementary Table 7-8**). These 69 genes were significantly enriched for seven gene ontology terms (**Supplementary Table 9**), with “very-low-density lipoprotein particle” being the most significant (adjusted *P* = 5.8 × 10^-3^).

**Figure 6.**
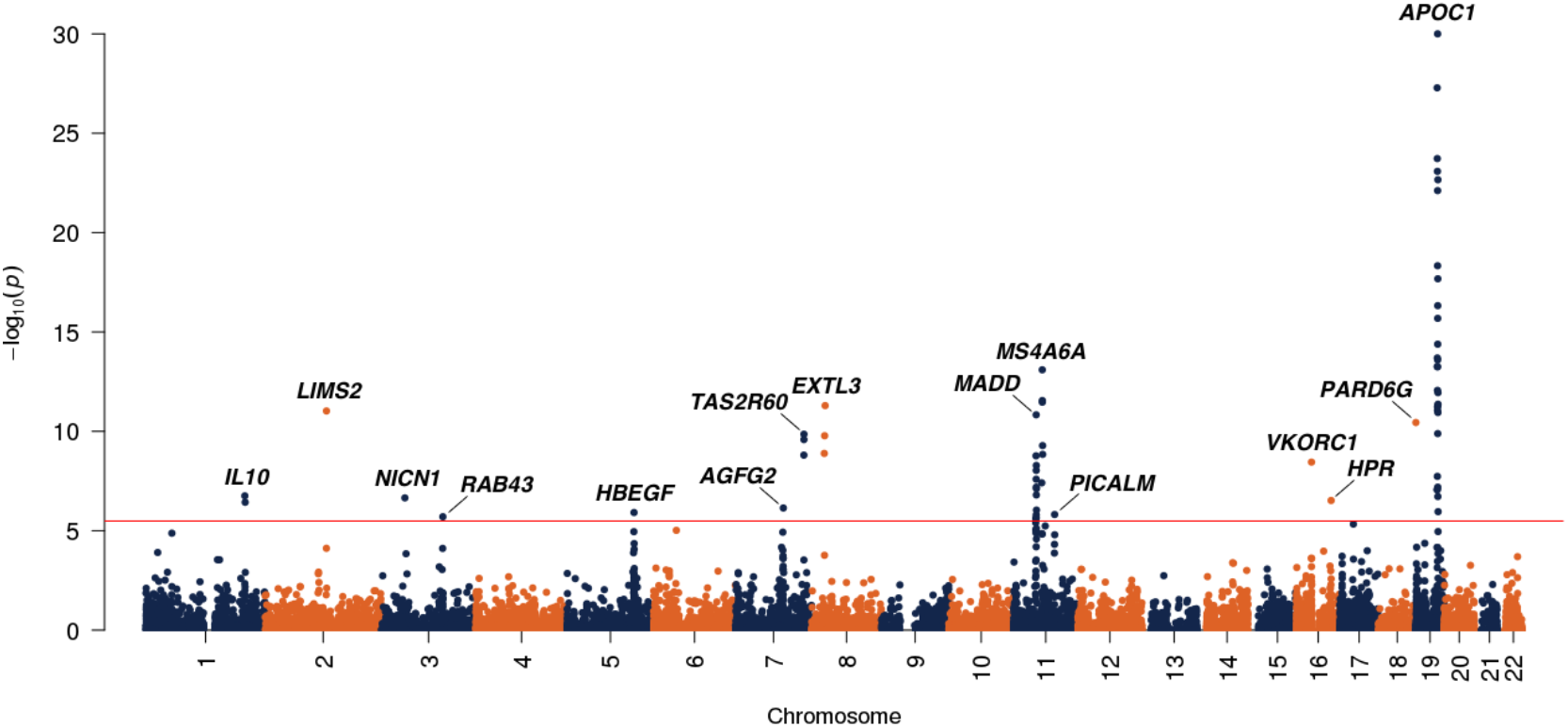
Manhattan plot for LOAD meta-analysis. P-values are truncated at 1 × 10^-30^ for visualization purpose. The horizontal line marks the genome-wide significance threshold. The most significant gene at each locus is labeled. (n = 168,726, generalized Berk-Jones test)

Most significant genes are from previously identified LOAD risk loci^40,46–51^. These include *CR1* locus on chromosome 1, *BIN1* locus on chromosome 2, *HBEGF* locus on chromosome 5, *ZCWPW1* and *EPHA1* loci on chromosome 7, *CLU* locus on chromosome 8, *CELF1*, MS4A6A, and *PICALM* loci on chromosome 11, and the *APOE* region on chromosome 19. Among these loci, *AGFG2* rather than ZCWPW1, the previously-suggested index gene at this locus^40^, was significant in the meta-analysis (*P* = 7.19 × 10^-7^). Similarly, *BIN1* was not statistically significant in our analysis. But LIMS2, a gene 500 kb upstream of BIN1, was significantly associated (*P* = 9.43 × 10^-12^). SNPs in the 3’UTR of LIMS2 have been previously suggested to associate with cognitive decline^52^. GWAS index genes for the rest of the loci were all statistically significant in our analysis.

Further, new associations at known risk loci provide novel insights into LOAD etiology. We identified a novel gene IL10 for LOAD risk (*P* = 1.77 × 10^-7^). IL10 is 700 kb upstream of CR1, a strong and consistently replicated locus in LOAD GWAS^40,51,53^. CR1 is also significant in our analysis (*P* = 3.71 × 10^-7^). Although some SNPs near the promoter region of IL10 were moderately associated with LOAD in all three datasets (**Supplementary Figure 9**), the IL10-LOAD association was mostly driven by SNPs near CR1 (**Supplementary Table 10**). An interesting observation is that even when a key SNP is missing - the most significant SNP in IGAP and ADGC (i.e. rs2093761: A>G) was not present in GWAX, other predictors (e.g. rs6690215:C>T in GWAX) still helped recover the association signal at the gene level, leading to a genome-wide significant association at IL10. To investigate if IL10 is simply a companion association signal due to co-regulation with CR1, we performed a cross-tissue conditional analysis using UTMOST with both significant genes CR1 and IL10 included in the model (**Online Methods**). Only IL10 remained significant (*P* = 1.4 × 10^-7^ for IL10 and *P* = 0.11 for CR1, **Supplementary Table 11**) in the conditional analysis. In addition to strong statistical evidence, the biological function of IL10 also supports its association with LOAD. IL10 is associated with multiple immune diseases^54–57^. It is known to encode one of the main anti-inflammatory cytokines associated with the occurrence of Alzheimer’s disease and has therapeutic potential to improve neurodegeneration^58,59^. Its protein product is also known to physically interact with the Tau protein^60^.

*CLU* is another well-replicated risk gene for LOAD. Two independent association peaks at this locus, one at *CLU* and the other at PTK2B, have previously been identified in GWAS (**Supplementary Figure 10**)^40,51^. In our analysis, in addition to *CLU* (*P* = 1.66 × 10^-10^), we identified two more significant genes at this locus, i.e. **ADRA1A** (*P* = 1.29 × 10^-9^) and *EXTL3* (*P* = 5.08 × 10^-12^). PTK2B showed marginal association (*P* = 1.72 × 10^-4^) with LOAD but did not reach genome-wide significance. Interestingly, *EXTL3* expression is predicted by a SNP in the LOAD association peak at *CLU* while **ADRA1A** is regulated by SNPs at both *CLU* and PTK2B (**Supplementary Table 12**). **ADRA1A** has been implicated in gene-gene interaction analysis for LOAD^61^. Its protein product physically interacts with amyloid precursor protein (APP)^60^ and an α1-adrenoceptor antagonist has been shown to prevent memory deficits in APP23 transgenic mice^62^. *EXTL3* encodes a putative membrane receptor for regenerating islet-derived 1α (Reg-1α), whose overexpression and involvement in the early stages of Alzheimer’s disease has been reported^63^. Further, the effect of Reg-1α on neurite outgrowth is mediated through EXTL3. Our results provide additional evidence that **IL10**, **ADRA1A**, and *EXTL3* may be involved in LOAD etiology.

Finally, we identified five novel loci for LOAD, each represented by one significant gene: *NICN1* (*P* = 2.23 × 10^-7^), *RAB43* (*P* = 1.98 × 10^-6^), *VKORC1* (*P* = 3.53 × 10^-9^), *HPR* (*P* = 3.02 × 10^-7^), and *PARD6G* (*P* = 3.60 × 10^-11^). The Rab GTPases are central regulators of intracellular membrane trafficking^64^. Although *RAB43* has not been previously identified in LOAD GWAS, USP6NL, the gene that encodes a GTPase-activating protein for *RAB43*, has been identified to associate with LOAD in two recent studies^45,50^. *USP6NL* also showed suggestive association with LOAD in the discovery stage of our analysis (*P* = 0.004). However, the associations at *RAB43* and *USP6NL* were not strongly supported by ADGC or GWAX datasets. Further, the *RAB43*-LOAD association was driven by SNPs near RPN1, a gene 400 kb downstream of *RAB43* (**Supplementary Figure 11, Supplementary Table 13**). This locus is associated with a variety of blood cell traits including monocyte count^65,66^. *VKORC1* is a critical gene in vitamin K metabolism and is the target of warfarin^67^, a commonly prescribed anticoagulant. It is known that the *APOE ε4* allele affects the efficacy of warfarin^68^. HPR has been identified to strongly associate with multiple lipid traits^69^ and interact with *APOE*^60^. *NICN1* is known to associate with inflammatory bowel disease^70^ and cognitive function^71^. These results provide potential target genes for functional validations in the future. The cross-tissue imputation models of these genes were listed in **Supplementary Tables 14-20**.

## Discussion

Despite the many improvements of UTMOST over existing methods, researchers need to be cautious when interpreting findings from UTMOST analyses. First, gene-level associations identified in UTMOST do not imply causality. It has been recently discussed that correlations among the imputed expression of multiple genes at the same locus may lead to apparent associations at non-causal genes^20^, which is comparable to linkage disequilibrium (LD)’s impact on SNP-level associations in GWAs. Consequently, TWAS-type approaches have limitations in both inferring functional genes and relevant tissues. When eQTL of different genes at the same locus are shared or in LD, irrelevant genes may be identified through significant associations. Similarly, for a given gene, if eQTL for the same gene in different tissues are shared or in LD, irrelevant tissues may show significant association signals. UTMOST cross-tissue conditional analysis can resolve the issue of gene prioritization to some extent, but fine-mapping of gene-level association remains challenging, especially in regions with extensive LD. We performed simulations to show that true associations in the causal tissue were consistently stronger than those in the non-causal tissue in most scenarios, which indicated that single-tissue association analyses have the potential to infer causal tissue (**Supplementary Note; Supplementary Figure 12**). However, as the proportion of shared eQTL increases, p-values for associations in the non-causal tissue became increasingly significant. Even when two tissues do not share eQTL, associations in the non-causal tissue still frequently passed the significance threshold, most likely due to LD between eQTL. These results are consistent with our experience and discussions in the literature^20,72^. We also note that these issues may become even more complex when sample sizes and imputation power vary across tissues. Further, we emphasize one of the principles in hypothesis testing - one should not conclude the null hypothesis when an association is not statistically significant. UTMOST is a general framework that involves many analytical steps, and technical issues might mask true gene-trait associations. For example, *SPI1* from the *CELF1* locus has been causally linked to LOAD risk^42^. We identified multiple significant associations at this locus but SPI1 was not a significant gene in our analysis. Possible reasons for this include insufficient imputation quality based on the current model, non-availability of causal tissue in the training data, key eQTL missing from the GWAS summary statistics, causal mechanism (e.g. alternative splicing) not well-represented in our analysis, or insufficient sample sizes. In practice, these issues need to be carefully investigated before ruling out any candidate gene.

Overall, UTMOST is a novel, powerful, and flexible framework to perform gene-level association analysis. It integrates biologically-informed weights with GWAS summary statistics via modern statistical techniques. Interpreted with caution, its findings may provide insights into disease and trait etiology, motivate downstream functional validation efforts, and eventually benefit the development of novel therapeutics. It is also exciting that statistical and computational methodology in this field evolves at a fast pace. Several methods on mediation analysis and functional gene fine-mapping in the context of transcriptome-wide association study have been proposed recently^73,74^. It has been shown that data-adaptive sNp weights could effectively improve statistical power at the cost of clear interpretation of associations^75^. Extension of these methods into multi-tissue analysis is an interesting possible future direction. As high-throughput data continue to be generated for more individuals, cell types, and molecular phenotypes, UTMOST promises to show even better performance and provide greater insights for complex disease genetics in the future.

## URLs

UTMOST software: https://github.com/Joker-Jerome/UTMOST

BLUEPRINT: ftp://.ebi.ac.uk/pub/databases/blueprint/blueprint_Epivar/qtlas/

STARNET: https://github.com/Wainberg/Vulnerabilities_of_TWAS

AlzData: http://alzdata.org/index.html

GLGC: http://lipidgenetics.org

IGAP: http://web.pasteur-lille.fr/en/recherche/u744/igap/igap_download.php

TWAS summary statistics: ftp://ftp.biostat.wisc.edu/pub/lu_group/Projects/UTMOST

GEUV: https://www.ebi.ac.uk/arrayexpress/experiments/E-GEUV-1/

GWAX: http://gwas-browser.nygenome.org/downloads/

GTEx: https://www.gtexportal.org

ADGC2 summary statistics: https://www.niagads.org/datasets/ngΩΩΩ76

## Supporting information

LargeSupplementaryTables

## Acknowledgements

This study was supported in part by the National Institutes of Health grants R01 GM59507 and NIH 3P30AG021342-16S2 (H.Z., Y.H., M.L.), the VA Cooperative Studies Program of the Department of Veterans Affairs, Office of Research and Development, and the Yale World Scholars Program sponsored by the China Scholarship Council (J.W., Z.L., B.L.). Q.L. was supported by the Clinical and Translational Science Award (CTSA) program, through the NIH National Center for Advancing Translational Sciences (NCaTS), grant UL1TR000427. The content is solely the responsibility of the authors and does not necessarily represent the official views of the NiH. P.G. and Sh.M. were supported by grant R01 aG042437 and U01 AG006781. J.G. and H.L. were supported by Neil shen’s SJTU Medical Research Fund. We thank Dr. Christopher Brown for his assistance in matching GTEx tissues to Roadmap cell types. This study makes use of summary statistics from many GWAS consortia. We thank the investigators in these GWAS consortia for generously sharing their data. We thank the International Genomics of Alzheimer’s Project (IGAP) for providing summary results data for these analyses. The investigators within IGAP contributed to the design and implementation of IGAP and/or provided data but did not participate in analysis or writing of this report. IGAP was made possible by the generous participation of the subjects and their families. The i-Select chips were funded by the French National Foundation on Alzheimer’s disease and related disorders. EADI was supported by the LABEX (laboratory of excellence program investment for the future) DISTALZ grant, Inserm, Institut Pasteur de Lille, Université de Lille 2, and the Lille University Hospital. GERAD was supported by the Medical Research Council (Grant n° 503480), Alzheimer’s Research UK (Grant n° 503176), the Wellcome Trust (Grant n° 082604/2/07/Z), and German Federal Ministry of Education and Research (BMBF): Competence Network Dementia (CND) grant n° 01GI0102, 01GI0711, 01GI0420. CHARGE was partly supported by the NIH/NIA grant R01 AG033193 and the NIA AG081220 and AGES contract N01-AG-12100, the NHLBI grant R01 HL105756, the Icelandic Heart Association, and the Erasmus Medical Center and Erasmus University. ADGC was supported by the NIH/NIA grants: U01 AG032984, U24 AG021886, U01 AG016976, and the Alzheimer’s Association grant ADGC-10-196728. We thank contributors who collected samples used in this study, as well as patients and their families, whose help and participation made this work possible; Data for this study were prepared, archived, and distributed by the National Institute on Aging Alzheimer’s Disease Data Storage Site (NIAGADS) at the University of Pennsylvania (U24-AG041689-01). We are also grateful for all the consortia and investigators that provided publicly accessible GWAS summary statistics.

## Author contribution

Y.H., M.L., Q.L., H.L., and H.Z. conceived the study and developed the statistical model.Y.H., M.L., Q.L., H.W., J.W., S.M.Z., B.L., Y.S., Sy.M. and J.G. performed the statistical analyses. S.M.Z. and P.N. assisted in LDL analysis. Y.H., M.L., Z.Y., and Q.L. implemented the software. B.K. prepared ADGC summary statistics. A.N., A.K. and Y.Z. assisted in data preparation Sh.M. and P.C. assisted in Alzheimer’s disease data application, curation, and interpretation. Y.H., M.L., Q.L., H.L., and H.Z. wrote the manuscript. H. Z. advised on statistical and genetic issues. All authors contributed in manuscript editing and approved the manuscript.

## Online Methods

### Penalized regression model for cross-tissue expression imputation

Given a gene, we use genotype information to predict its covariate-adjusted expression levels in *P* tissues. We use SNPs between 1 Mb upstream of the transcription start site and 1 Mb downstream of the transcription end site of the given gene as predictor variables in the model. This is denoted as an *N × M* matrix *X* where *N* is the total number of individuals and *M* denotes the number of SNPs. Throughout the paper, we assume each column of *X* to be centered but not standardized. Of note, expression data may not be available for all individuals since only a subset of tissues were collected from each individual. For the ith tissue, we use *N_i_* to denote its sample size. We further use an *N_i_*-dimensional vector *Y_i_* to denote the observed expression data in the ith tissue, and use an *N_i_× M* matrix *X_i_* to denote the genotype information for the subset of individuals. Then, cross-tissue gene expression imputation can be formulated as the following regression problem.

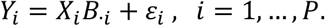

Here, the *M × P* matrix *B* summarizes SNPs’ effects on the given gene with its ith column B·_i_ denoting the effect sizes of SNPs in the *i*th tissue and the *j*th row *B._j_* denoting the effect sizes of the *j*th SNP in all *P* tissues. To effectively select biologically relevant and statistically predictive SNPs, accurately estimate their effects across tissues, and address technical issues including shared samples and incomplete data, we propose the following penalized least-squares estimator for genetic effects matrix *B*:

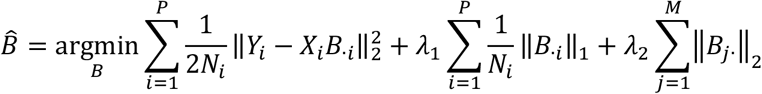

Here, ‖.‖_1_and ‖.‖_2_ denote the *l*_1_ and *l*_2_norms, respectively (i.e.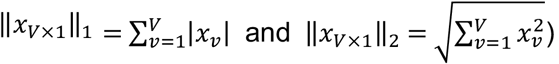. The first term in the loss function is the standard least-squares error. We use the *l*_1_ penalty to select predictive variables and impose shrinkage in effect size estimation. The penalty on each tissue is set adaptively based on the sample sizes, which reflects the idea that models for tissues with a larger sample size are more robust to overfitting and therefore are penalized less. To integrate information across multiple tissues, we introduced the third term - a group-lasso penalty on the effect sizes of one SNP ^29^. By imposing this joint penalty across tissues, UTMOST encourages eQTLs shared across tissues but still keeps tissue-specific eQTLs with strong effects. Although the penalty on tissue-specific eQTL may cause the model to exclude some true predictors, recent evidence ^76^ suggested that tissue-specific eQTL have substantially weaker effect sizes and will most likely not have major influences on association analysis (**Supplementary Note**). Tuning parameters λ1 and λ2 control the within-tissue and cross-tissue sparsity, respectively. They are selected through cross-validation. Details of optimization were attached in **Supplementary Note**.

### Model training and evaluation

We trained our cross-tissue gene expression imputation model using genotype and normalized gene expression data from 44 tissues in the GTEx project (version V6p, dbGaP accession code: phs000424.v6.p1)^3^. Sample sizes for different tissues ranged from 70 (uterus) to 361 (skeletal muscle). SNPs with ambiguous alleles or minor allele frequency (MAF) < 0.01 were removed. Normalized gene expressions were further adjusted to remove potential confounding effects from sex, sequencing platform, top three principal components of genotype data, and top probabilistic estimation of expression residuals (PEER) factors^77^. As previously recommended^17^, we included 15

PEER factors for tissues with *N* ≤ 150, 30 factors for tissues with 150 ≤ *N*< 250, and 35 factors for tissues with *N* ≥ 250. All covariates were downloaded from the GTEx portal website (**URLs**). We applied a 5-fold cross-validation for model tuning and evaluation. Specifically, we randomly divided individuals into five groups of equal size. Each time, we used three groups as the training set, one as the intermediate set for selecting tuning parameters, and the last one as the testing set for performance evaluation. Squared correlation between predicted and observed expression (i.e. R^2^) was used to quantify imputation accuracy. For each model, we selected gene-tissue pairs with FDR < 0.05 for downstream testing. External validation of imputation accuracy was performed using whole-blood expression data from 421 samples in the 1000 Genomes Project (GEUVADIS consortium)^32^ and the CommonMind consortium^33^, which collected expression in across multiple regions from > 1,000 postmortem brain samples (mainly corresponding to Brain_Frontal_Cortex_BA9 in GTEx) from donors with schizophrenia, bipolar disorder, and individuals with no neuropsychiatric disorders. For CommonMind data, we focused our analysis on 147 controls with no neuropsychiatric disorders. Average improvements in R^2^ in both external validation datasets are shown in **Supplementary** Figure 4. Although not statistically significant due to the limited sample size, the accuracy of the cross-tissue method was consistently higher than that of the single-tissue approach in different quantiles. Furthermore, comparing the tissue-tissue similarity based on the observed and imputed gene expressions indicated that cross-tissue imputation removed stochastic noises in the expression data without losing tissue-specific correlational patterns (*Supplementary Note; Supplementary Figure 5-6*).

### Gene-level association test

We combined GWAS summary statistics with SNP effects estimated in the cross-tissue imputation model (i.e. 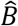) to quantify gene-trait associations in each tissue. For a given gene, we modeled its imputed expression in the *i*th tissue (i.e. 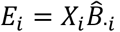 and the phenotype *T* using a linear model

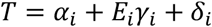

Then, the association statistic for effect size in the *i*th tissue (i.e. *γ_i_*) on the trait of interest is

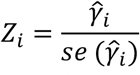

where 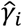 denotes the point estimate for effect size and *se* 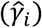 denotes its standard error. From the linear model, we have

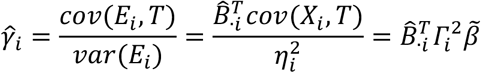

where *Г_i_* is an *M × M* diagonal matrix with the *j*th term equal to 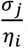, where *σ_j_* is the standard deviation of the *j*th SNP, and *η_i_* is the standard deviation of imputed gene expression in the *i*th tissue. These parameters could be estimated using a reference panel. 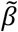 denotes the SNP-level effect size estimates acquired from GWAS summary statistics. Regarding the standard error of 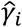, we have

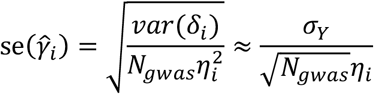

Here, *σ_Y_* denotes the standard deviation of phenotype *T* and *N_gaws_* is the sample size in GWAS. The approximation 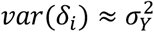 is based on the empirical observation that each gene only explains a very small proportion of phenotypic variability78. The same argument can be extended to association statistics at the SNP level. For the *j*th SNP in the model, we have

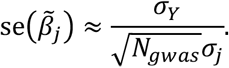

Therefore, SNP-level z-scores can be denoted as

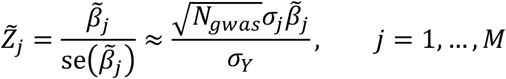

In matrix form, this is

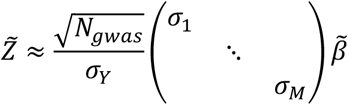

Combining the derivations above, we can denote the gene-level z-score as

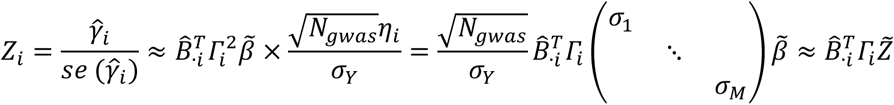

Under the null hypothesis (i.e. no SNP-trait association), 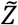 follows a multivariate normal distribution 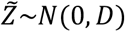 where *D* is the LD matrix for SNPs and could be estimated using an external reference panel. Denoting the cross-tissue gene-trait z-scores as *Z = (Z_1_,Z_2_,…, Z_P_)^T^*, the covariance matrix of *Z* could be calculated as

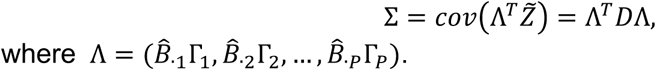

In order to combine gene-trait associations across multiple tissues, we applied the generalized Berk-Jones (GBJ) test with single-tissue association statistics *Z* and their covariance matrix Σ as inputs. This approach provides powerful inference results while explicitly taking the correlation among single-tissue test statistics into account even under a sparse alternative (i.e. biologically meaningful associations are only present in a small number tissues)^30^. The GBJ test statistic can be calculated as

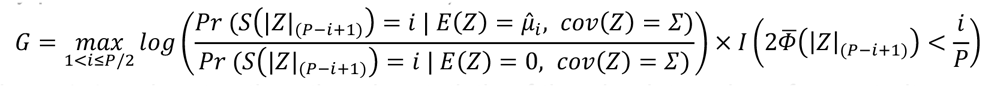

Where |z|*_(i)_* denotes the *i*th order statistic of the absolute value of gene-trait z-scores in an increasing order; 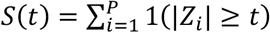> denotes the number of gene-trait z-scores with absolute value greater than a threshold t; 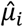 denotes the corresponding value of E(Z) that maximizes the probability of event *s(|Z|_(P-i+1)_) = i*; and 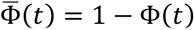 is the survival function of the standard normal distribution. The GBJ test statistic can be interpreted as the maximum of a series of one-sided likelihood ratio test statistics on the mean of *S(t)*, where the denominator denotes the maximum likelihood when no gene-trait association exists in any tissue (all z-scores have zero mean) and the numerator denotes the unconstrained maximum likelihood. Of note, calculating the exact distribution of *S(t)* is difficult when z-scores are correlated. As previously suggested, we calculate G by approximating the distribution of *S(t)* with an extended beta-binomial (EBB) distribution. As a maximum-based global statistic, the p-value of GBJ test could be written as

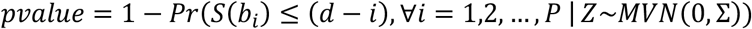

where 0 ≤ *b*_1_ ≤ *b*_2_ ≤…≤ *b_P_* are ‘boundary points’ derived from inversion of the test statistic, which depends on *G, P* and Σ. The last quantity in the equation can be calculated recursively with the EBB approximation^30^.

P-value cut-offs for gene-level association tests were determined by Bonferroni correction. For each method, we used 0.05 divided by the total number of genes tested across 44 tissues (i.e. 5.76 × 10^-7^ for TWAS, 2.44 × 10^-7^ for PrediXcan, and 1. 28 × 10^-7^ for UTMOST, respectively) as the significance threshold. As more genes can be accurately imputed (*R*^2^ significantly larger than zero with FDR < 0.05) in our cross-tissue imputation, the significance cutoff was the most stringent in UTMOST.

### Cross-tissue conditional analysis

Genes that are physically close to the true risk gene may be identified in marginal association analyses due to co-regulation of multiple genes by the same eQTL and LD between eQTL of different genes. In order to prioritize gene-level associations at the same locus, we expand UTMOST to perform cross-tissue conditional analysis. There are two major steps in this framework:

First, at any pre-defined locus, we can derive the formula of conditional analysis based on marginal associations. Denote T as the trait of interest. The goal is to perform a multiple regression analysis using K imputed gene expressions in the ith tissue (i.e. *E_i1_,…, E_i__k_*) as predictor variables:

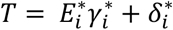

Here, we use 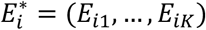 to denote an *N × K* matrix for *K* imputed gene expressions in the *i*th tissue. Regression coefficients 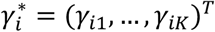 are the parameters of interest. To simplify algebra, we also assume that trait *T* and all SNPs in the genotype matrix *X* are centered so there is no intercept term in the model, but the conclusions apply to the general setting. Similar to univariate analysis, gene expressions *E_i1_,…, E_ik_* are imputed from genetic data via linear prediction models:

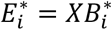

where 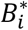 are imputation weights assigned to SNPs. The k^t^^h^ column of 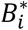 denotes the imputation model for gene expression *E_ik_*. Then, the OLS estimator 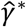 and its variance-covariance matrix can be denoted as follows:

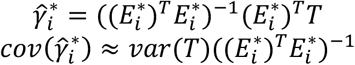

The approximation is based on the assumption that imputed gene expressions *E_i1_,…, E_ik_* collectively explain little variance in *T*, which is reasonable in complex gene expression genetics if *K* is not large. We further denote:

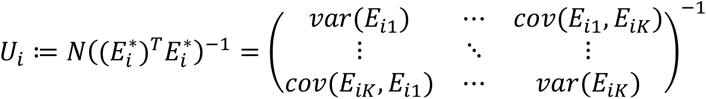

All elements in matrix *U_i_* can be approximated using a reference panel 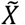. Therefore, the z-score for *γ_ik_(1≤k ≤K)* is

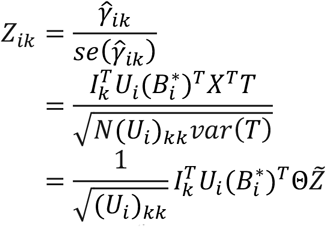

where *I_k_* is the *K* ×1 vector with the *k*^th^ element being 1 and all other elements equal to 0, Θ is a *M × M* diagonal matrix with the *j*^th^ diagonal element being 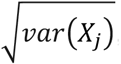 and similar to the notation in univariate analysis, 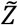 is the vector of SNP-level z-scores from the GWAS of trait *T*. Importantly, we note that given imputation models for *K* gene expressions (i.e. 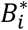), GWAS summary statistics for trait *T* (i.e. 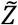), and an external genetic dataset to estimate *U_i_* and Θ, conditional analysis can be performed without individual-level genotype and phenotype data.

In the second step, we combine the conditional analysis association statistics across different tissues using the GBJ test. Note this is different from the final stage of UTMOST, which combines the marginal gene-trait-tissue associations. Through these two steps, LD between eQTL and co-regulation across tissues has been taken into account in the test. Specifically, under the null hypothesis (i.e. no SNP-trait association), 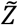 follows a multivariate normal distribution 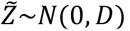 where *D* is the LD matrix for SNPs and could be estimated using an external reference panel. Denoting the cross-tissue gene-trait z-scores for gene *k* as *Z_k_ = (Z_1k_,Z_2k_,…, Z_Pk_)^T^*, the covariance matrix of *Z_k_* could be calculated as

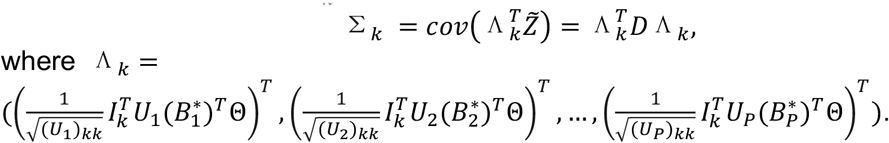

### Simulation settings

Genotype data from 12,637 individuals in the GERA dataset (dbGaP accession: phs000674), including 7,432 type-2 diabetes cases (phenotypic information not used) and 5,205 healthy controls, were used in the simulation studies. We removed SNPs with missing rate above 0.01 and individuals with genetic relatedness coefficients above 0.05. The genotype data were imputed to the 1000 Genomes Project Phase 1v3 European samples using the Michigan Imputation Server^79^. After imputation, we further removed SNPs with MAF < 0.05. After quality control, 5,932,546 SNPs remained in the dataset.

We performed two different simulation studies to evaluate the type-I error rate of our cross-tissue association test. First, we directly simulated quantitative traits from a standard normal distribution independent from the genotype data, and then performed single-tissue association tests for 44 tissues in GTEx and GBJ cross-tissue association test for all genes using the simulated data. In the second setting, we simulated genetically-regulated expression components and then simulated the GWAS trait based on gene expression values. For each gene, we simulated its expression in three tissues, namely skeletal muscle (N = 361), skin from sun-exposed lower leg (N = 302), and whole blood (N = 338). Within the i th tissue, the cis-component of gene expression was generated as 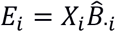 We used real effect sizes 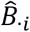estimated in our joint imputation model so that the genetic architecture of gene expression was preserved in the simulations. Next, the quantitative trait value was simulated as *Y = w_1_E_1_ + w_2_E_2_+w_3_E_3_ + ε*, where w_i_ is the effect of gene expression on the trait in the *i*th tissue. To evaluate type-I error, we set *w_1_ = w_2_ = w_3_= 0*, i.e. none of the three tissues are relevant to the trait.

To simulate data under the alternative hypothesis, we generated diverse disease architectures by considering different number of causal tissues (i.e. 1, 2, or 3) and two heritability settings (i.e. 0.01 and 0.001). Specifically, we fixed the total variance explained by *E_1_, E_2_*, and E3 and varied *w_i_* to simulate different levels of tissue specificity of the trait. We generated traits using the following three settings:

*Setting 1*. *W_1_ = 1, w _2_ =w_3_ = 0*. Only the first tissue contributes to the disease, the other two tissues are not relevant.

*Setting 2*. 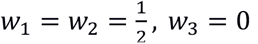. Both the first and the second tissue contribute equally to disease, the third tissue is irrelevant to the disease.

*Setting 3*. 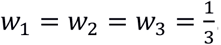 All three tissues contribute equally to the disease.

Single-tissue and cross-tissue gene-trait associations were then estimated using the UTMOST framework. We repeated the whole procedure on 200 randomly selected genes. For each gene, we further replicated 5 times. Statistical power is calculated as the proportion of test p-values reaching the significance threshold, i.e. 0.05/15000 for both single-tissue and cross-tissue tests and 0.05/45000 for single tissue tests while accounting for the number of tissues.

### GWAS data analysis

We applied UTMOST to GWAS summary statistics for 50 complex diseases and traits. Details of these 50 studies are summarized in **Supplementary Table 4**. GWAS summary statistics for LDL cholesterol was downloaded from the Global Lipids Genetics Consortium website (URLs). Summary statistics from the IGAP stage-I analysis was downloaded from the IGAP website (URLs). GWAX result for LOAD was downloaded from New York Genome Center website (URLs). ADGC phase 2 summary statistics were generated by first analyzing individual datasets using logistic regression adjusting for age, sex and the first three principal components in the program SNPTest v2^80^. Meta-analysis of the individual dataset results was then performed using the inverse-variance weighted approach in METAL^81^.

To identify trait-related tissue, we first used GenoSkyline-Plus, an unsupervised learning framework trained on various epigenetic marks from the Roadmap Epigenomics Project ^82^, to quantify tissue-specific functionality in the human genome ^83.^ We then estimated the enrichment for trait heritability in each tissue’s predicted functional genome using LD score regression ^34^. More specifically, annotation-stratified LD scores were estimated using the 1000 Genomes samples of European ancestry and a 1-centiMorgan window. GenoSkyline-Plus annotations for 27 tissues that can be matched between Roadmap and GTEx were included in the LD score regression model together with 53 baseline annotations, as previously suggested ^34^. For each tissue-specific annotation, partitioned heritability was estimated and enrichment was calculated as the ratio of the proportion of explained heritability and the proportion of SNPs in each annotated category. Tissue-trait relevance was then ranked based on enrichment p-values. We use term “most enriched tissues” to denote the tissues that were most significantly enriched for heritability of each trait. Authors of ^84^ also applied LDSC with tissue specific annotations based on GTEx data to infer trait-related tissues. Since UTMOST was based on GTEx data, we used an independent data from the Roadmap project to infer trait-relevant tissues for the purpose of fair comparison.

In the UTMOST analytical framework, multiple parameters need to be estimated using an external reference panel (e.g. LD). We used samples with European ancestry from the 1000 Genomes Project for this estimation^85^. When performing cross-tissue association tests, we combined single-tissue statistics from tissues that passed FDR < 0.05 criteria to reduce noise in the analysis. Genome-wide significance was defined as 3.3 × 10^-6^ (i.e. Bonferroni correction based on 15,120 genes that passed the quality control steps). For heritability enrichment analysis, we applied LDSC to 27 GenoSkyline-Plus tissue-specific annotations that have matched tissue types in GTEx (**Supplementary Table 21**). The 53 LDSC baseline annotations were also included in the model as previously recommended^34^. The most and least relevant tissues were selected based on the enrichment test p-values. Gene ontology enrichment analysis was performed using DAVID^86^. Protein-protein interaction information was acquired from AlzData website (URLs)^60^. Locus plots for SNP-level GWAS associations were generated using LocusZoom^87^. Manhattan plots were generated using the qqman package in R^88^.

### Additional QTL data

Imputation model for liver tissue in the STARNET study (N = 522) was downloaded from (URLs). Predictor effects were trained using an elastic-net model with variants within 500kb range of the transcription-starting site. Details on the quality control procedure has been previously reported^20^. We have also collected additional eQTL and sQTL data for three immune cell types (CD14+ monocytes, CD16+ neutrophils, and naive CD4+ T cells; 169-194 samples per tissue) from the BLUEPRINT consortium (URLs). eQTLs with FDR < 0.01 and sQTLs with FDR < 0.05 were used in the gene-level association analysis for LOAD.

We also downloaded monocyte eQTL summary statistics from the Immune Variation Project^89^ as a comparison with BLUEPRINT results in LOAD. We first compared the monocyte eQTL identified in BLUEPRINT with what was identified in this dataset (denote as ImmVar). Only a very low fraction (3.5%) of the eQTLs could be replicated in ImmVar. We further performed single-tissue analysis on LOAD with weights constructed from ImmVar and compared the identified associations with those identified using BLUEPRINT data (**Supplementary Tables 22-23**). Significant genes did not match between the two analyses which is most likely due to the small overlap of eQTLs between two datasets. However, UTMOST uses the Generalized Berk-Jones statistic to combine associations across datasets and therefore has the flexibility to incorporate single-tissue associations based on external eQTL studies. As we demonstrated in the case study of LDL-C at the *SORT1* locus, incorporating STARNET liver eQTL significantly increased the statistical power despite the fact that liver was an available tissue in GTEx. As sample sizes and tissue types in QTL studies continue to grow, UTMOST will be able to incorporate additional data sources and provide better results.

### Statistical tests

We tested the difference in R^2^ across genes with one-sided Kolmogorov-Smirnov test, which calculates the largest distance between the empirical cumulative distribution functions and uses it to test if two distributions are identical (**Supplementary Figures 3-4**). And we used a paired Wilcoxon rank test to compare the number of genes identified in different tissues between different methods, which is a non-parametric test used to compare two matched samples to access whether their population mean differ (Figure 4, **Supplementary Figure 7**).

## Data Availability

All data used in the manuscript are publicly available (see URLs). GTEx and GERA data can be accessed by application to dbGaP. CommonMind data are available through formal application to NIMH. ADGC phase 2 summary statistics used for validation are available through NIAGADS portal (see URLs) with accession number NG00076.

