## SupplementaryMaterials for "A statistical framework for cross-tissue transcriptome-wide association analysis"

### Supplementary Notes

#### Optimization of cross-tissue imputation model

To minimize the objective function, we developed a novel coordinate descent approach that extended the optimization procedure in [79] to deal with incomplete data in the outcome matrix. In each iteration, we update a row of $B$. When $B_{(-j)\cdot}$ (i.e. all the rows except for the $j$th) have been updated, the procedure is equivalent to minimizing

$$L(B_{j\cdot}) \coloneqq L(B_{j\cdot}; B_{(-j)\cdot}, Y_1, \dots, Y_P, X_1, \dots, X_P) = \sum_{i=1}^{P} \frac{1}{2N_i} \left\| \tilde{Y}_i - (X_i)_{\cdot j} B_{ji} \right\|_2^2 + \lambda_1 \sum_{i=1}^{P} \frac{1}{N_i} |B_{ji}| + \lambda_2 \sqrt{\sum_{i=1}^{P} B_{ji}^2}$$

where $\tilde{Y}_i = Y_i - (X_i)_{\cdot(-j)} B_{(-j)i}$ and $B_{j\cdot} = (B_{j1}, \dots, B_{jP})$. $(X_i)_{\cdot j}$ and $(X_i)_{\cdot(-j)}$ denote the $j$th column and the matrix excluding the $j$th column of $X_i$ ($N_i \times M$), respectively. For a specific $i \in (1, \dots, P)$, when $B_{ji} > 0$, its partial derivative is

$$\begin{aligned} \frac{\partial L}{\partial B_{ji}} &= \frac{(X_i)_{\cdot j}^T (X_i)_{\cdot j}}{N_i} B_{ji} - \frac{(X_i)_{\cdot j}^T \tilde{Y}_i}{N_i} + \frac{\lambda_1}{N_i} + \frac{\lambda_2}{\|B_{j\cdot}\|_2} B_{ji} \\ &= \left( \frac{(X_i)_{\cdot j}^T (X_i)_{\cdot j}}{N_i} + \frac{\lambda_2}{\|B_{j\cdot}\|_2} \right) B_{ji} - \left( \frac{(X_i)_{\cdot j}^T \tilde{Y}_i}{N_i} - \frac{\lambda_1}{N_i} \right) \end{aligned}$$

Therefore, when $B_{ji} > 0$, condition $\frac{\partial L}{\partial B_{ji}} > 0$ is equivalent to $B_{ji} > \tilde{B}_{ji}^+$, where $\tilde{B}_{ji}^+$ is defined as

$$\tilde{B}_{ji}^+ \coloneqq \frac{\frac{(X_i)_{\cdot j}^T \tilde{Y}_i}{N_i} - \frac{\lambda_1}{N_i}}{\frac{(X_i)_{\cdot j}^T (X_i)_{\cdot j}}{N_i} + \frac{\lambda_2}{\|B_{j\cdot}\|_2}} = \frac{\frac{(X_i)_{\cdot j}^T \tilde{Y}_i}{N_i}}{\frac{(X_i)_{\cdot j}^T (X_i)_{\cdot j}}{N_i} + \frac{\lambda_2}{\|B_{j\cdot}\|_2}} \left( 1 - \frac{\lambda_1}{(X_i)_{\cdot j}^T \tilde{Y}_i} \right)$$

And the minimum of $L(B_{ji}; B_{j(-i)})$ with $B_{ji} > 0$ and $B_{j(-i)}$ fixed is

$$\hat{B}_{ji,min}^+ \coloneqq \begin{cases} \tilde{B}_{ji}^+, & if\ \frac{(X_i)_{\cdot j}^T \tilde{Y}_i}{N_i} \left( 1 - \frac{\lambda_1}{(X_i)_{\cdot j}^T \tilde{Y}_i} \right) > 0 \\ 0, & if\ \frac{(X_i)_{\cdot j}^T \tilde{Y}_i}{N_i} \left( 1 - \frac{\lambda_1}{(X_i)_{\cdot j}^T \tilde{Y}_i} \right) \le 0 \end{cases}$$

Similarly, when $B_{ji} \le 0$, we have

$$\frac{\partial L}{\partial B_{ji}} = \left( \frac{(X_i)_{\cdot j}^T (X_i)_{\cdot j}}{N_i} + \frac{\lambda_2}{\|B_{j\cdot}\|_2} \right) B_{ji} - \left( \frac{(X_i)_{\cdot j}^T \tilde{Y}_i}{N_i} + \frac{\lambda_1}{N_i} \right)$$

Condition $\frac{\partial L}{\partial B_{ji}} > 0$ is equivalent to $B_{ji} > \tilde{B}_{ji}^-$, where $\tilde{B}_{ji}^-$ is defined as

$$\tilde{B}_{ji}^- \coloneqq \frac{\frac{(X_i)_{\cdot j}^T \tilde{Y}_i}{N_i} + \frac{\lambda_1}{N_i}}{\frac{(X_i)_{\cdot j}^T (X_i)_{\cdot j}}{N_i} + \frac{\lambda_2}{\|B_{j\cdot}\|_2}} = \frac{\frac{(X_i)_{\cdot j}^T \tilde{Y}_i}{N_i}}{\frac{(X_i)_{\cdot j}^T (X_i)_{\cdot j}}{N_i} + \frac{\lambda_2}{\|B_{j\cdot}\|_2}} \left( 1 + \frac{\lambda_1}{(X_i)_{\cdot j}^T \tilde{Y}_i} \right)$$

The minimum of $L(B_{ji}; B_{j(-i)})$ with $B_{ji} \le 0$ and $B_{j(-i)}$ fixed is

$$\hat{B}_{ji,min}^- \coloneqq \begin{cases} \tilde{B}_{ji}^-, if\ \frac{(X_i)_{\cdot j}^T \tilde{Y}_i}{N_i} \left( 1 + \frac{\lambda_1}{(X_i)_{\cdot j}^T \tilde{Y}_i} \right) < 0 \\ 0, if\ \frac{(X_i)_{\cdot j}^T \tilde{Y}_i}{N_i} \left( 1 + \frac{\lambda_1}{(X_i)_{\cdot j}^T \tilde{Y}_i} \right) \ge 0 \end{cases}$$

Denote the minimum of $L\left(B_{ji}; B_{j(-i)}\right)$ with $B_{j(-i)}$ fixed as $\hat{B}_{ji,min}$. If $(X_i)_{\cdot j}^T\tilde{Y}_i > 0$, and $\frac{\partial L}{\partial B_{ji}} < 0$ for $B_{ji} < 0$, then $B_{ji,min} \geq 0$ and

$$\hat{B}_{ji,min} = \hat{B}_{ji,min}^{+} = \frac{\frac{(X_i)_{\cdot j}^T\tilde{Y}_i}{N_i}}{\frac{(X_i)_{\cdot j}^T(X_i)_{\cdot j}}{N_i} + \frac{\lambda_2}{\|B_{j\cdot}\|_2}}\left(1 - \frac{\lambda_1}{\left|(X_i)_{\cdot j}^T\tilde{Y}_i\right|}\right)_+$$

Similarly, if $(X_i)_{\cdot j}^T\tilde{Y}_i \leq 0$ and $\frac{\partial L}{\partial B_{ji}} \geq 0$ for $B_{ji} \geq 0$, then $\hat{B}_{ji,min} \leq 0$ and

$$\hat{B}_{ji,min} = \hat{B}_{ji,min}^{-} = \frac{\frac{(X_i)_{\cdot j}^T\tilde{Y}_i}{N_i}}{\frac{(X_i)_{\cdot j}^T(X_i)_{\cdot j}}{N_i} + \frac{\lambda_2}{\|B_{j\cdot}\|_2}}\left(1 - \frac{\lambda_1}{\left|(X_i)_{\cdot j}^T\tilde{Y}_i\right|}\right)_+$$

which is the same formula as in the previous case. Denoting the minimum of $L\left(B_{j\cdot}\right)$ as $\hat{B}_{j\cdot,min} := \left(\hat{B}_{j1,min}, \dots, \hat{B}_{jP,min}\right)$, we have shown that if such a minimum exists, it should satisfy:

$$\hat{B}_{ji,min} = \frac{\frac{(X_i)_{\cdot j}^T\tilde{Y}_i}{N_i}}{\frac{(X_i)_{\cdot j}^T(X_i)_{\cdot j}}{N_i} + \frac{\lambda_2}{\|\hat{B}_{j\cdot,min}\|_2}}\left(1 - \frac{\lambda_1}{\left|(X_i)_{\cdot j}^T\tilde{Y}_i\right|}\right)_+ = \frac{\frac{X_{ij\cdot}^T X_{ij\cdot}}{N_i}}{\frac{(X_i)_{\cdot j}^T(X_i)_{\cdot j}}{N_i} + \frac{\lambda_2}{\|\hat{B}_{j\cdot,min}\|_2}}\hat{B}_{ji}^{lasso}$$

where $\hat{B}_{ji}^{lasso} = \frac{(X_i)_{\cdot j}^T\tilde{Y}_i}{(X_i)_{\cdot j}^T(X_i)_{\cdot j}}\left(1 - \frac{\lambda_1}{\left|(X_i)_{\cdot j}^T\tilde{Y}_i\right|}\right)_+$ and $i = 1, \dots, P$. And minimizing $L\left(B_{j\cdot}\right)$ is the equivalent to solving the above non-linear equation system. However, this system does not have closed-form solution and solving it numerically is very computationally expensive, especially considering we need to train over 15,000 models. To address this challenge, we proposed a efficient procedure to approximate $\hat{B}_{ji,min}$.

Note that $\frac{(X_i)_{\cdot j}^T(X_i)_{\cdot j}}{N_i}$ is the diagonal term of the LD matrix estimated by $N_i$ individuals, we could thus approximate it with the LD matrix estimated with entire genotype matrix, i.e. $\frac{X_{\cdot j}^T X_{\cdot j}}{N}$, and we got a new non-linear equation systems, which is close to the original one but has a closed-form solution.

$$\hat{B}_{ji,min} \approx \frac{\frac{X_{\cdot j}^T X_{\cdot j}}{N}}{\frac{X_{\cdot j}^T X_{\cdot j}}{N} + \frac{\lambda_2}{\|\hat{B}_{j\cdot,min}\|_2}}\hat{B}_{ji}^{lasso}, \quad i = 1, \dots, P$$

By squaring both sides of the approximation and summing over $i$, we obtain the following approximation:

$$\|\hat{B}_{j\cdot,min}\|_2 \approx \frac{\frac{X_{\cdot j}^T X_{\cdot j}}{N}}{\frac{X_{\cdot j}^T X_{\cdot j}}{N} + \frac{\lambda_2}{\|\hat{B}_{j\cdot,min}\|_2}}\|\hat{B}_{j\cdot}^{lasso}\|_2$$

where $\hat{B}_{j\cdot}^{lasso} = (\hat{B}_{j1}^{lasso}, \dots, \hat{B}_{jP}^{lasso})$. That is

$$\|\hat{B}_{j\cdot,min}\|_2 \approx \|\hat{B}_{j\cdot}^{lasso}\|_2 - \frac{N\lambda_2}{X_{\cdot j}^T X_{\cdot j}}$$

And given $B_{(-j)\cdot}$, $B_{j\cdot}$ is updated by

$$\hat{B}_{j\cdot} = \left(1 - \frac{\lambda_2}{\left\|\hat{B}_{j\cdot}^{lasso}\right\|_2 \frac{X_{\cdot j}^T X_{\cdot j}}{N}}\right)\hat{B}_{j\cdot}^{lasso}$$

We keep training the model until convergence in $B$ or the mean squared error starts increasing in the validation set. Numerical experiments demonstrated that the proposed procedure provided reasonable approximation in each iteration and resulted in robust prediction models (**Figure 2** and **Supplementary Table 1**).

##### Tissue-tissue similarity based on original and imputed gene expression

We computed the tissue-tissue similarity matrix with the observed expression levels and predicted expression levels in the following way:

1) For each gene, calculate the pairwise correlation between two tissues based on individuals who have expression levels measured in both tissues. Similarly for predicted expression, predicting gene expression on individuals with measured expression in both tissues and compute the correlation between them.
2) For each tissue-tissue pair, use the average of the absolute values of correlations across genes as the correlation between two tissues.

As shown in **Supplementary Figure 5**, we observed that the predicted gene expression levels in general have higher correlations, which is as expected, since fitting the prediction models reduces noise in gene expression. But the similarity patterns of tissues were preserved. Specifically, 1) we stacked elements of two similarity matrices as two vectors and the correlation between them is 0.52 (6.1E-65), which demonstrates that the similarity patterns are highly preserved; 2) We compared the similarities within 10 brain tissues and similarities of brain tissues with the rest tissues for both observed and imputed gene expression levels. As expected, for observed gene expression levels, similarities within brain tissues are larger than their similarities with other tissues and the same pattern remains for imputed gene expression levels, indicating that although the overall correlations between tissues increase, the similarity patterns and the tissue specific expression patterns are preserved (**Supplementary Figure 6**).

##### Single-tissue association and cross-tissue eQTL

To investigate how cross-tissue eQTLs affect single-tissue association, we performed simulations to study how the associations in causal tissues and non-causal tissues change under different levels of eQTL sharing. We first simulated gene expression levels and a continuous trait using the same genotype data described in **Section Methods** (**Simulation settings)**. We randomly selected a gene, *GLMN* (ENSG00000174842), and used SNPs within its 50MB flanking region (~2,000 SNPs in total) to simulate gene expression levels. We simulated gene expression for two tissues with different numbers of shared eQTL. Specifically,

1. We randomly selected eQTL. Denote the number of eQTLs in tissue 1 as $n_1$ and that in tissue 2 as $n_2$, and denote the number of shared eQTLs between them as $n_s$. We used $\frac{n_s}{(n_1+n_2)/2}$ to measure the extent of co-regulation between two tissues.
2. Effect sizes of cross-tissue eQTLs are generated from a normal distribution $N(0,\ \sigma_s^2)$ and effect sizes of tissue-specific eQTL are generated from $N(0,\ \sigma_t^2)$, where $\sigma_t^2 = 0.02^2$ and $\sigma_s^2 = 0.05^2$. Note that we set the parameters so that the cross-tissue eQTLs have larger effect sizes and these numbers were based on the estimates from GTEx.

3. The trait values were simulated by assuming tissue 1 as the causal tissue. Heritability of the gene was set to be 0.05.
4. Finally, we calculated the associations between trait and expression in two tissues under different values of $\frac{n_s}{(n_1+n_2)/2}$. For each setting, we ran 100 replicates.

#### Other relevant penalized regression methods for gene expression imputation

In related papers [1-3], other penalized regression models including LASSO, elastic net, and Bayesian sparse linear mixed effect model (BSLMM) have been applied in the TWAS framework to predict gene expression levels. LASSO assumes effect sizes are sparse and shrinks smaller effects to zero and could be too aggressive when the actual effect sizes are small and less sparse. On the other hand, elastic net adds an additional quadratic term to the penalty, which is a robust alternative to LASSO. However, both methods penalize each coefficient individually and could not efficiently utilize the structure information of variables; as illustrated in Figure 1 in [4], the contour of LASSO penalty has a polyhedron-shape and each coordinate direction is treated differently from other directions, and this encourages sparsity in individual coefficients. The L2-penalty (squared $l_2$-norm) has a sphere shape and treats all directions equally and does not encourage sparsity. The group lasso has cone shape and encourages sparsity at the group-level. For example when a group of variable should be selected into or out of a model together based on prior knowledge, which is suitable for the cross-tissue prediction, eQTLs with large effect sizes usually regulate multiple tissues at the same time. Group LASSO penalty could select the cross-tissue eQTLs with strong effect sizes and achieve more robust prediction. In this case, group LASSO is a natural choice to incorporate external information and outperforms LASSO when group structure exists [4]. Furthermore, Austin et al. showed in [5] that elastic net outperforms lasso and ridge regression under different genetic architectures. Similarly in [3, 6], Zhou et al. demonstrated that BSLMM, a Bayesian sparse regression method, has the similar flexibility as elastic-net and could therefore perform well in both sparse and polygenic models. They also showed that BSLMM outperforms traditional Bayesian variable selection methods in many cases. We have empirically compared our approach with elastic net and BSLMM. Both cross-validation and external validations showed that our model achieves better imputation performance and is more robust in tissues with limited sample sizes.

#### LDL-C analysis using only STARNET data

We performed single-tissue association analysis based on STARNET data only on LDL-C GWAS. In *SORT1* locus, the most significant genes identified were *CELSR2* and *PSRC1*. (**Supplementary Figure 13**). However, when combining GTEx tissues with STARNET, *SORT1* showed up as the most significant signal at this locus, which demonstrates that cross-tissue approach can better prioritize causal genes (**Figure 5a**). Furthermore, STARNET single-tissue analysis (**Supplementary Figure 13**) identified 8 genes at this locus (all included in the 14 genes identified by cross-tissue analysis). STARNET has 5,963 gene imputation models and including 15 genes at this locus, while UTMOST has 15,120 genes for testing, including 27 at this locus. Therefore, even if *SORT1* is the only causal gene in this locus, the cross-tissue approach did not increase the false-positive rate.

### Supplementary Tables

#### Supplementary Table 1. Improvement in expression imputation across 44 GTEx tissues.

| | | Number of effectively imputed genes | | |
|---|---|---|---|---|
| **Tissue** | **Sample size** | **Elastic net** | **UTMOST** | **Improvement** |
| Adipose Subcutaneous | 298 | 7249 | 9707 | 33.9% |
| Adipose Visceral Omentum | 185 | 4568 | 9366 | 105.0% |
| Adrenal Gland | 126 | 4174 | 8752 | 109.7% |
| Artery Aorta | 197 | 6182 | 9042 | 46.3% |
| Artery Coronary | 118 | 3222 | 9003 | 179.4% |
| Artery Tibial | 285 | 7121 | 9299 | 30.6% |
| Brain Anterior cingulate cortex BA24 | 72 | 2559 | 7810 | 205.2% |
| Brain Caudate basal ganglia | 100 | 3544 | 8659 | 144.3% |
| Brain Cerebellar Hemisphere | 89 | 4068 | 8274 | 103.4% |
| Brain Cerebellum | 103 | 4995 | 8577 | 71.7% |
| Brain Cortex | 96 | 3558 | 8427 | 136.8% |
| Brain Frontal Cortex BA9 | 92 | 3258 | 8543 | 162.2% |
| Brain Hippocampus | 81 | 2566 | 8370 | 226.2% |
| Brain Hypothalamus | 81 | 2451 | 8589 | 250.4% |
| Brain Nucleus accumbens basal ganglia | 93 | 3057 | 8572 | 180.4% |
| Brain Putamen basal ganglia | 82 | 2749 | 8054 | 193.0% |
| Breast Mammary Tissue | 183 | 4648 | 9709 | 108.9% |
| Cells EBV-transformed lymphocytes | 114 | 3660 | 7921 | 116.4% |
| Cells Transformed fibroblasts | 272 | 7609 | 8625 | 13.4% |
| Colon Sigmoid | 124 | 3720 | 8831 | 137.4% |
| Colon Transverse | 169 | 4788 | 9408 | 96.5% |
| Esophagus Gastroesophageal Junction | 127 | 3601 | 8792 | 144.2% |
| Esophagus Mucosa | 241 | 6889 | 9387 | 36.3% |
| Esophagus Muscularis | 218 | 6533 | 9273 | 41.9% |
| Heart Atrial Appendage | 159 | 4565 | 8474 | 85.6% |
| Heart Left Ventricle | 190 | 4858 | 8150 | 67.8% |
| Liver | 97 | 2759 | 7792 | 182.4% |
| Lung | 278 | 6564 | 10086 | 53.7% |
| Muscle Skeletal | 361 | 6563 | 8645 | 31.7% |
| Nerve Tibial | 256 | 8113 | 9883 | 21.8% |
| Ovary | 85 | 2880 | 8572 | 197.6% |
| Pancreas | 149 | 4931 | 8348 | 69.3% |
| Pituitary | 87 | 3335 | 9049 | 171.3% |
| Prostate | 87 | 2614 | 9329 | 256.9% |
| Skin - Not Sun Exposed Suprapubic | 196 | 5633 | 9540 | 69.4% |
| Skin - Sun Exposed Lower leg | 302 | 7567 | 10038 | 32.7% |
| Small Intestine Terminal Ileum | 77 | 2613 | 8702 | 233.0% |
| Spleen | 89 | 3715 | 8587 | 131.1% |
| Stomach | 170 | 4096 | 9272 | 126.4% |
| Testis | 157 | 7043 | 10482 | 48.8% |
| Thyroid | 278 | 8026 | 9991 | 24.5% |
| Uterus | 70 | 2159 | 8004 | 270.7% |
| Vagina | 79 | 2041 | 8959 | 339.0% |
| Whole Blood | 338 | 6650 | 8410 | 26.5% |

**Supplementary Table 2. Type-I error rate in the first simulation analysis.** Quantitative trait values were directly simulated from a standard normal distribution, independent from genotype data.

| $\alpha$ | Type-I error rate |
|---|---|
| 0.1 | 6.57E-02 |
| 0.01 | 8.22E-03 |
| 0.001 | 9.70E-04 |
| 0.0001 | 6.47E-05 |

**Supplementary Table 3. Type-I error rate in the second simulation analysis.** Quantitative trait values were simulated based on genetically-regulated expression values in three tissues.

| | **Type-I error rate** | | | |
|---|---|---|---|---|
| **α** | **Joint** | **Muscle** | **Skin** | **Blood** |
| 0.010 | 0.006 | 0.006 | 0.011 | 0.013 |
| 0.020 | 0.011 | 0.011 | 0.017 | 0.020 |
| 0.030 | 0.017 | 0.030 | 0.026 | 0.032 |
| 0.040 | 0.025 | 0.039 | 0.031 | 0.041 |
| 0.050 | 0.032 | 0.044 | 0.043 | 0.046 |

**Supplementary Table 4. Information of 50 GWAS traits.**

| Trait | Acronym | Reference |
|---|---|---|
| Age at First Birth | AFB | [7] |
| Age at Menarche | AM | [8] |
| Age at Natural Menopause | ANM | [9] |
| Age-related Macular Degeneration | AMD | [10] |
| Alzheimer’s Disease | AD | [11] |
| Amyotrophic Lateral Sclerosis | ALS | [12] |
| Anorexia Nervosa | AN | [13] |
| Anxiety Disorder | ANX | [14] |
| Asthma | AST | [15] |
| Autism Spectrum Disorder | ASD | [16] |
| Bipolar Disorder | BIP | [17] |
| Birth Weight | BW | [18] |
| Body Mass Index | BMI | [19] |
| Celiac Disease | CEL | [20] |
| Chronic Kidney Disease | CKD | [21] |
| Chronotype | CHT | [22] |
| Cognitive Performance | COG | [23] |
| Coronary Artery Disease | CAD | [24] |
| Crohn's Disease | CD | [25] |
| Depressive Symptoms | DEP | [26] |
| Eczema | ECZ | [27] |
| Education Years | EDU | [28] |
| Epilepsy | EPL | [29] |
| Fasting Glucose | GLU | [30] |
| Fasting Insulin | INS | [30] |
| Femoral Neck Bone Mineral Density | FNBMD | [31] |
| Gout | GOUT | [32] |
| HDL Cholesterol | HDL | [33] |
| Height | HGT | [34] |
| Inflammatory Bowel Disease | IBD | [25] |
| LDL Cholesterol | LDL | [33] |
| Lumbar Spine Bone Mineral Density | LSBMD | [31] |
| Major Depressive Disorder | MDD | [35] |
| Multiple Sclerosis | MS | [36] |
| Neuroticism | NEU | [26] |
| Number of Children Ever Born | NCEB | [7] |
| Primary Angle Closure Glaucoma | PACG | [37] |
| Primary Biliary Cirrhosis | PBC | [38] |
| Resting Heart Rate | RHR | [39] |
| Rheumatoid Arthritis | RA | [40] |
| Schizophrenia | SCZ | [41] |
| Serum Urate | SU | [32] |
| Smoking Behavior | SMK | [42] |
| Subjective Well-being | SWB | [26] |
| Systemic Lupus Erythematosus | SLE | [43] |
| Total Cholesterol | TC | [33] |
| Triglycerides | TG | [33] |
| Type-II Diabetes | T2D | [44] |
| Ulcerative Colitis | UC | [25] |
| Waist Hip Ratio adjusted for BMI | WHR | [45] |

**Supplementary Table 5. Significant associations in the most relevant tissue for 50 complex traits.**

| Trait | TWAS | MetaXcan | UTMOST |
|---|---|---|---|
| AD | 0 | 0 | 8 |
| AFB | 2 | 5 | 8 |
| ALS | 0 | 0 | 1 |
| AM | 0 | 4 | 9 |
| AMD | 5 | 25 | 16 |
| AN | 0 | 0 | 0 |
| ANM | 14 | 21 | 25 |
| ANX | 0 | 0 | 0 |
| ASD | 0 | 0 | 0 |
| AST | 2 | 1 | 2 |
| BIP | 2 | 1 | 3 |
| BMI | 13 | 16 | 39 |
| BW | 12 | 13 | 20 |
| CAD | 4 | 4 | 12 |
| CD | 5 | 7 | 23 |
| CEL | 0 | 1 | 2 |
| CHT | 0 | 1 | 2 |
| CKD | 0 | 0 | 2 |
| COG | 0 | 1 | 3 |
| DEP | 0 | 1 | 2 |
| ECZ | 0 | 0 | 2 |
| EDU | 25 | 24 | 55 |
| EPL | 0 | 0 | 0 |
| FNBMD | 3 | 3 | 7 |
| GLU | 0 | 0 | 6 |
| GOUT | 2 | 2 | 2 |
| HDL | 30 | 33 | 52 |
| HGT | 95 | 134 | 267 |
| IBD | 13 | 23 | 38 |
| INS | 0 | 0 | 0 |
| LDL | 5 | 10 | 22 |
| LSBMD | 9 | 3 | 7 |
| MDD | 0 | 0 | 0 |
| MS | 4 | 15 | 4 |
| NCEB | 0 | 0 | 0 |
| NEU | 12 | 4 | 14 |
| PACG | 0 | 1 | 1 |
| PBC | 3 | 15 | 7 |
| RA | 0 | 32 | 16 |
| RHR | 11 | 24 | 34 |
| SCZ | 12 | 26 | 72 |
| SLE | 1 | 22 | 23 |
| SMK | 0 | 0 | 0 |
| SU | 1 | 7 | 23 |
| SWB | 0 | 0 | 0 |
| T2D | 0 | 3 | 1 |
| TC | 6 | 10 | 26 |
| TG | 19 | 32 | 37 |
| UC | 4 | 12 | 11 |
| WHR | 6 | 9 | 18 |

**Supplementary Table 6. Pairwise conditional analysis of *SORT1* and other significant genes.** p_Gene: UTMOST cross-tissue association; p_*SORT1*: p value of *SORT1* in pairwise conditional analysis with the corresponding gene (n=173,082, two-sided z-score test); r_STARNET: Pearson correlation between imputed gene expression in the STARNET data (n = 522)

| Gene | p_Gene | p_*SORT1* | r_STARNET |
|---|---|---|---|
| *AMIGO1* | 1.07E-07 | 2.33E-13 | -0.17 |
| *ATXN7L2* | 1.01E-04 | 2.34E-13 | -0.26 |
| *CELSR2* | 2.36E-11 | 5.67E-03 | 0.90 |
| *GNAT2* | 2.91E-09 | 2.33E-13 | NA |
| *GPR61* | 2.18E-11 | 2.33E-13 | NA |
| *GSTM4* | 1.72E-01 | 2.32E-13 | 0.17 |
| *GSTM5* | 3.48E-12 | 2.33E-13 | 0.21 |
| *KIAA1324* | 2.18E-11 | 2.33E-13 | NA |
| *MYBPHL* | 3.60E-11 | 2.32E-13 | NA |
| *PSMA5* | 1.31E-13 | 2.30E-13 | 0.18 |
| *PSRC1* | 2.31E-11 | 8.89E-03 | 0.90 |
| *SYPL2* | 7.18E-03 | 2.33E-13 | -0.40 |
| *TAF13* | 2.25E-03 | 2.33E-13 | 0.02 |
| *WDR47* | 2.18E-11 | 2.33E-13 | NA |

**Supplementary Table 7. Association results for 68 genes that were significant in the discovery-stage analysis for LOAD** (IGAP: n = 54,162; ADGC2: n = 7,050; GWAX: n = 114,564; generalized Berk-Jones test; the last column: n = ,175,776 Fisher's test).

| Gene | Chr | Coordinates | p_IGAP | p_ADGC2 | p_GWAX | p_Fisher |
|---|---|---|---|---|---|---|
| *IL10* | 1 | 206940947-206945839 | 6.16E-08 | 0.111 | 0.106 | 1.77E-07 |
| *CR1* | 1 | 207669492-207813992 | 2.09E-06 | 0.108 | 0.007 | 3.71E-07 |
| *MYO7B* | 2 | 128293378-128395304 | 1.54E-06 | 0.532 | 0.802 | 7.67E-05 |
| *LIMS2* | 2 | 128395956-128439360 | 1.77E-14 | 1.000 | 1.000 | 9.43E-12 |
| *RAB43* | 3 | 128806412-128841644 | 8.79E-07 | 0.485 | 0.025 | 1.98E-06 |
| *PROB1* | 5 | 138727635-138730885 | 2.31E-06 | 1.000 | 0.520 | 1.29E-04 |
| *NRG2* | 5 | 139226364-139422884 | 1.12E-07 | 1.000 | 0.657 | 1.12E-05 |
| *HBEGF* | 5 | 139712428-139726216 | 2.14E-08 | 1.000 | 0.286 | 1.22E-06 |
| *CD14* | 5 | 140011313-140013286 | 2.31E-06 | 0.588 | 0.259 | 4.45E-05 |
| *HARS2* | 5 | 140071011-140078889 | 2.61E-06 | 1.000 | 0.287 | 8.57E-05 |
| *AGFG2* | 7 | 100136834-100165842 | 1.33E-06 | 0.070 | 0.037 | 7.19E-07 |
| *CASP2* | 7 | 142985308-143004789 | 6.42E-12 | 0.769 | 0.861 | 1.57E-09 |
| *EPHA1* | 7 | 143087382-143105985 | 5.66E-12 | 0.701 | 0.154 | 2.60E-10 |
| *TAS2R60* | 7 | 143140546-143141502 | 1.27E-12 | 0.246 | 1.000 | 1.39E-10 |
| *ADRA1A* | 8 | 26605667-26724790 | 2.07E-10 | 0.761 | 0.022 | 1.29E-09 |
| *CLU* | 8 | 27454434-27472548 | 1.81E-11 | 0.266 | 0.079 | 1.66E-10 |
| *EXTL3* | 8 | 28457986-28613116 | 7.23E-12 | 0.069 | 0.018 | 5.08E-12 |
| *MDK* | 11 | 46402306-46405375 | 2.54E-06 | 0.009 | 1.000 | 3.97E-06 |
| *ATG13* | 11 | 46638826-46696368 | 7.82E-07 | 0.024 | 0.704 | 2.46E-06 |
| *LRP4* | 11 | 46878419-46940193 | 7.19E-07 | 0.058 | 1.11E-04 | 1.71E-09 |
| *MADD* | 11 | 47290712-47351582 | 1.54E-08 | 0.005 | 3.51E-04 | 1.48E-11 |
| *SLC39A13* | 11 | 47428683-47438047 | 7.64E-08 | 0.030 | 0.013 | 9.22E-09 |
| *RAPSN* | 11 | 47459308-47470730 | 1.89E-06 | 0.065 | 0.002 | 6.42E-08 |
| *CELF1* | 11 | 47487496-47587121 | 2.85E-07 | 0.017 | 0.003 | 5.20E-09 |
| *MTCH2* | 11 | 47638867-47664175 | 1.54E-06 | 0.031 | 1.000 | 7.74E-06 |
| *FNBP4* | 11 | 47738072-47788995 | 4.80E-07 | 9.27E-04 | 0.647 | 7.61E-08 |
| *NUP160* | 11 | 47799639-47870107 | 9.65E-08 | 0.046 | 1.000 | 9.13E-07 |
| *FAM111B* | 11 | 58874658-58894883 | 2.20E-08 | 0.041 | 0.153 | 3.90E-08 |
| *MS4A2* | 11 | 59855734-59863444 | 1.14E-06 | 0.089 | 1.000 | 1.49E-05 |
| *MS4A6A* | 11 | 59939081-59952139 | 1.87E-13 | 8.62E-04 | 0.699 | 8.02E-14 |
| *MS4A4E* | 11 | 59968726-60010561 | 7.10E-09 | 0.003 | 1.000 | 7.65E-09 |
| *MS4A4A* | 11 | 60048014-60076445 | 1.30E-10 | 2.89E-04 | 0.159 | 3.41E-12 |
| *MS4A6E* | 11 | 60102304-60164069 | 6.66E-10 | 5.46E-05 | 0.135 | 2.83E-12 |
| *PTGDR2* | 11 | 60618413-60623444 | 6.27E-09 | 2.06E-04 | 1.000 | 5.21E-10 |
| *SLC15A3* | 11 | 60704556-60720002 | 3.38E-09 | 0.003 | 0.437 | 1.44E-09 |
| *LRRC25* | 19 | 18501954-18508427 | 1.46E-06 | 0.233 | 1.000 | 4.30E-05 |
| *ZNF221* | 19 | 44455375-44471861 | 1.27E-11 | 1.25E-09 | 1.27E-11 | 5.19E-28 |
| *ZNF223* | 19 | 44555520-44572144 | 4.25E-07 | 7.48E-04 | 1.000 | 8.33E-08 |
| *ZNF227* | 19 | 44711700-44741421 | 2.18E-11 | 2.72E-11 | 1.59E-06 | 1.89E-24 |
| *ZNF235* | 19 | 44732882-44809199 | 1.48E-13 | 0.004 | 0.048 | 2.03E-14 |
| *ZNF233* | 19 | 44754318-44779470 | 3.08E-08 | 9.36E-06 | 2.59E-04 | 5.45E-14 |
| *ZNF112* | 19 | 44830708-44871377 | 5.00E-08 | 0.026 | 0.047 | 1.84E-08 |
| *ZNF180* | 19 | 44979854-45004576 | 2.36E-11 | 1.31E-10 | 1.40E-06 | 8.28E-24 |
| *IGSF23* | 19 | 45116940-45140081 | 2.98E-12 | 0.007 | 0.071 | 8.69E-13 |
| *PVR* | 19 | 45147098-45166850 | 5.73E-11 | 9.88E-04 | 0.188 | 5.88E-12 |
| *CEACAM19* | 19 | 45165545-45187631 | 2.35E-11 | 5.23E-05 | 0.064 | 5.72E-14 |
| *CEACAM16* | 19 | 45202421-45213986 | 5.46E-12 | 0.026 | 0.118 | 8.88E-12 |
| *BCAM* | 19 | 45312328-45324673 | 4.02E-12 | 2.38E-06 | 3.83E-05 | 4.65E-19 |
| *PVRL2* | 19 | 45349432-45392485 | 2.36E-11 | 2.55E-11 | 2.55E-11 | 4.23E-29 |
| *APOE* | 19 | 45409011-45412650 | 1.45E-11 | 1.53E-11 | 1.96E-04 | 7.71E-23 |
| *APOC1* | 19 | 45417504-45422606 | 1.75E-59 | 1.99E-14 | 2.79E-29 | 2.65E-97 |
| *APOC4* | 19 | 45445495-45452820 | 4.90E-09 | 0.052 | 0.080 | 6.72E-09 |
| *APOC2* | 19 | 45449243-45452822 | 2.00E-11 | 3.36E-14 | 0.005 | 6.11E-24 |
| *CLPTM1* | 19 | 45457842-45496599 | 1.36E-10 | 6.36E-04 | 3.86E-04 | 2.53E-14 |
| *RELB* | 19 | 45504688-45541452 | 1.07E-08 | 0.634 | 0.317 | 4.75E-07 |
| *ZNF296* | 19 | 45574758-45579846 | 2.55E-11 | 1.82E-06 | 0.005 | 2.07E-16 |
| *PPP1R37* | 19 | 45594654-45651335 | 1.07E-11 | 0.002 | 2.13E-04 | 4.12E-15 |
| *NKPD1* | 19 | 45653008-45663408 | 7.08E-09 | 0.001 | 0.028 | 1.30E-10 |
| *TRAPPC6A* | 19 | 45666186-45681495 | 5.60E-08 | 0.244 | 0.018 | 6.42E-08 |
| *BLOC1S3* | 19 | 45682003-45685059 | 8.88E-16 | 0.002 | 0.030 | 4.75E-17 |
| *KLC3* | 19 | 45836692-45854778 | 7.67E-13 | 8.51E-05 | 2.71E-05 | 2.11E-18 |
| *ERCC1* | 19 | 45910591-45982086 | 2.18E-13 | 0.105 | 0.395 | 5.02E-12 |
| *RTN2* | 19 | 45988547-46000319 | 2.18E-11 | 2.97E-11 | 1.84E-05 | 2.20E-23 |
| *VASP* | 19 | 46009837-46030241 | 3.90E-08 | 0.004 | 1.50E-04 | 1.13E-11 |
| *EML2* | 19 | 46110252-46148887 | 4.34E-12 | 0.008 | 0.054 | 1.13E-12 |
| *QPCTL* | 19 | 46195741-46207247 | 6.84E-12 | 0.004 | 0.297 | 4.33E-12 |
| *DMWD* | 19 | 46286205-46296060 | 1.73E-06 | 1.000 | 0.778 | 1.43E-04 |
| *RSPH6A* | 19 | 46298968-46318577 | 1.67E-07 | 1.000 | 0.433 | 1.10E-05 |

**Supplementary Table 8. Association results for 12 new genes that became genome-wide significant in the meta-analysis** (IGAP: n = 54,162; ADGC2: n = 7,050; GWAX: n = 114,564; generalized Berk-Jones test; the last column: n = ,175,776 Fisher's test).

| Gene | Chr | Coordinates (hg19) | p_IGAP | p_ADGC2 | p_GWAX | p_Fisher |
|---|---|---|---|---|---|---|
| *NICN1* | 3 | 49460379-49466759 | 1.000 | 9.12E-02 | 1.03E-08 | 2.23E-07 |
| *NR1H3* | 11 | 47269851-47290396 | 1.55E-05 | 0.120 | 4.73E-05 | 2.57E-08 |
| *MYBPC3* | 11 | 47352957-47374253 | 9.87E-06 | 0.068 | 0.012 | 1.60E-06 |
| *PSMC3* | 11 | 47440320-47447993 | 4.17E-06 | 0.087 | 0.034 | 2.26E-06 |
| *FAM180B* | 11 | 47608198-47610746 | 3.44E-06 | 0.026 | 0.007 | 1.59E-07 |
| *PICALM* | 11 | 85668727-85780924 | 3.66E-06 | 0.048 | 0.045 | 1.53E-06 |
| *VKORC1* | 16 | 31102163-31107301 | 0.018 | 0.583 | 9.49E-10 | 3.53E-09 |
| *HPR* | 16 | 72088522-72111145 | 0.865 | 0.552 | 2.74E-09 | 3.02E-07 |
| *PARD6G* | 18 | 77915115-78005429 | 1.000 | 1.000 | 7.36E-14 | 3.60E-11 |
| *ZNF230* | 19 | 44507100-44518078 | 2.82E-05 | 0.002 | 0.007 | 8.77E-08 |
| *ERCC2* | 19 | 45853095-45874176 | 0.005 | 1.51E-07 | 1.000 | 1.91E-07 |
| *MYPOP* | 19 | 46393278-46405862 | 5.46E-06 | 0.001 | 1.000 | 1.10E-06 |

**Supplementary Table 9. Gene ontology enrichment results.** A total of 7 terms were significantly enriched after Benjamini-Hochberg correction (n = 15,120, Fisher's exact test).

| Category | GO Term | P-adjusted |
|---|---|---|
| Cellular component | very-low-density lipoprotein particle | 5.80E-03 |
| | spherical high-density lipoprotein particle | 2.10E-02 |
| | chylomicron | 4.40E-02 |
| Biological processes | triglyceride homeostasis | 4.80E-02 |
| | chylomicron remnant clearance | 4.50E-02 |
| | positive regulation of beta-amyloid formation | 4.20E-02 |
| | negative regulation of receptor-mediated endocytosis | 4.20E-02 |

**Supplementary Table 10. LOAD GWAS associations at SNPs that are predictors for *IL10* expression in at least one tissue in GTEx** (IGAP: n = 54,162; ADGC2: n = 7,050; GWAX: n = 114,564; generalized Berk-Jones test).

| | | | | | IGAP | | | ADGC2 | | | GWAX | | |
|---|---|---|---|---|---|---|---|---|---|---|---|---|---|
| **SNP** | **Chr** | **BP (hg19)** | **A1** | **A2** | **Beta** | **SE** | **P** | **Beta** | **SE** | **P** | **Beta** | **SE** | **P** |
| rs9242 | 1 | 206637395 | T | C | 0.003 | 0.016 | 0.858 | -0.050 | 0.040 | 0.220 | -0.002 | 0.013 | 0.904 |
| rs6666087 | 1 | 206708725 | C | T | 0.030 | 0.017 | 0.076 | -0.031 | 0.043 | 0.461 | 0.016 | 0.014 | 0.246 |
| rs2352794 | 1 | 207001040 | A | C | NA | NA | NA | -0.107 | 0.050 | 0.031 | 0.001 | 0.017 | 0.933 |
| rs10864035 | 1 | 207208335 | T | C | 0.022 | 0.016 | 0.171 | 0.062 | 0.041 | 0.131 | -0.012 | 0.014 | 0.383 |
| rs11120092 | 1 | 207211161 | G | A | 0.025 | 0.016 | 0.130 | 0.078 | 0.041 | 0.056 | -0.015 | 0.014 | 0.287 |
| rs1060286 | 1 | 207250300 | A | G | -0.027 | 0.016 | 0.091 | -0.066 | 0.042 | 0.114 | 0.010 | 0.014 | 0.475 |
| rs2842754 | 1 | 207257303 | C | T | 0.015 | 0.018 | 0.411 | -0.047 | 0.047 | 0.319 | NA | NA | NA |
| rs2491393 | 1 | 207300259 | G | A | -0.033 | 0.017 | 0.048 | -0.010 | 0.043 | 0.817 | -0.034 | 0.014 | 0.016 |
| rs6690215 | 1 | 207656050 | T | C | -0.057 | 0.016 | 5.44E-04 | -0.051 | 0.040 | 0.207 | -0.053 | 0.014 | 1.21E-04 |
| rs2093761 | 1 | 207786542 | G | A | -0.147 | 0.019 | 2.63E-14 | -0.169 | 0.051 | 9.23E-04 | NA | NA | NA |
| rs138953022 | 1 | 207859518 | C | T | -0.049 | 0.018 | 0.007 | -0.034 | 0.048 | 0.479 | NA | NA | NA |
| rs12073783 | 1 | 207875431 | G | A | -0.070 | 0.017 | 2.50E-05 | -0.022 | 0.042 | 0.605 | -0.028 | 0.014 | 0.041 |

**Supplementary Table 11. Cross-tissue conditional analysis results for genes that are significant in UTMOST cross-tissue analysis** (IGAP: n = 54,162; ADGC2: n = 7,050; GWAX: n = 114,564; generalized Berk-Jones test; the last column: n = ,175,776 Fisher's test).

| Gene | Chr | Coordinates | p_IGAP | p_ADGC2 | p_GWAX | p_Fisher |
|---|---|---|---|---|---|---|
| *IL10* | 1 | 206940947-206945839 | 5.88E-08 | 0.600 | 0.008 | 1.43E-07 |
| *CR1* | 1 | 207669492-207813992 | 0.023 | 0.678 | 0.137 | 0.111 |
| *CASP2* | 7 | 142985308-143004789 | 1.79E-09 | 0.817 | 0.687 | 4.75E-07 |
| *EPHA1* | 7 | 143087382-143105985 | 6.47E-09 | 1 | 0.290 | 8.37E-07 |
| *TAS2R60* | 7 | 143140546-143141502 | 0.004 | 0.811 | 1 | 0.158 |
| *ADRA1A* | 8 | 26605667-26724790 | 2.54E-08 | 0.509 | 0.002 | 1.92E-08 |
| *CLU* | 8 | 27454434-27472548 | 3.69E-11 | 0.349 | 0.154 | 1.55E-09 |
| *EXTL3* | 8 | 28457986-28613116 | 2.30E-10 | 0.169 | 0.079 | 2.33E-09 |
| *FAM111B* | 11 | 58874658-58894883 | 1.61E-05 | 2.68E-07 | 0.002 | 9.29E-12 |
| *MS4A6A* | 11 | 59939081-59952139 | 0.015 | 0.001 | 1 | 0.002 |
| *MS4A4E* | 11 | 59968726-60010561 | 1.52E-04 | 0.002 | 1 | 6.32E-05 |
| *MS4A4A* | 11 | 60048014-60076445 | 0.007 | 0.216 | 0.241 | 0.031 |
| *MS4A6E* | 11 | 60102304-60164069 | 0.014 | 0.095 | 0.273 | 0.029 |
| *PTGDR2* | 11 | 60618413-60623444 | 3.33E-08 | 5.15E-05 | 0.092 | 1.47E-10 |
| *SLC15A3* | 11 | 60704556-60720002 | 6.20E-07 | 1 | 0.567 | 8.88E-05 |

**Supplementary Table 12. LOAD GWAS associations at SNPs that are predictors for *CLU*, *ADRA1A*, and *EXTL3* expression in at least one tissue in GTEx** (IGAP: n = 54,162; ADGC2: n = 7,050; GWAX: n = 114,564; generalized Berk-Jones test).

| | | | | | | IGAP | | | ADGC2 | | | GWAX | | |
|---|---|---|---|---|---|---|---|---|---|---|---|---|---|---|
| **Gene** | **SNP** | **Chr** | **BP (hg19)** | **A1** | **A2** | **Beta** | **SE** | **P** | **Beta** | **SE** | **P** | **Beta** | **SE** | **P** |
| *ADRA1A* | rs11135904 | 8 | 25821871 | G | A | -0.010 | 0.016 | 0.539 | -0.014 | 0.041 | 0.728 | 0.002 | 0.014 | 0.861 |
| | rs2976321 | 8 | 25977337 | C | T | NA | NA | NA | -0.014 | 0.041 | 0.735 | -0.023 | 0.014 | 0.087 |
| | rs12544566 | 8 | 26293789 | A | C | -0.028 | 0.017 | 0.098 | -0.076 | 0.043 | 0.074 | 0.014 | 0.014 | 0.327 |
| | rs17055703 | 8 | 26551416 | C | A | -0.0001 | 0.017 | 0.994 | -0.036 | 0.044 | 0.420 | -0.011 | 0.015 | 0.473 |
| | rs12547624 | 8 | 26683805 | C | A | -0.013 | 0.021 | 0.544 | 0.037 | 0.053 | 0.487 | NA | NA | NA |
| | rs117380715 | 8 | 26696715 | A | G | -0.010 | 0.028 | 0.722 | 0.026 | 0.051 | 0.605 | -0.047 | 0.017 | 0.005 |
| | rs61761852 | 8 | 26697578 | C | T | -0.010 | 0.020 | 0.618 | 0.026 | 0.051 | 0.608 | -0.046 | 0.017 | 0.006 |
| | rs1079078 | 8 | 26698047 | C | A | -0.009 | 0.020 | 0.645 | 0.025 | 0.051 | 0.617 | -0.046 | 0.017 | 0.005 |
| | rs17056076 | 8 | 26709571 | G | A | -0.009 | 0.020 | 0.627 | 0.031 | 0.051 | 0.545 | -0.044 | 0.017 | 0.008 |
| | rs2046186 | 8 | 26711214 | G | A | -0.008 | 0.020 | 0.687 | 0.031 | 0.051 | 0.545 | -0.045 | 0.017 | 0.008 |
| | rs4639517 | 8 | 26728035 | T | C | 0.012 | 0.017 | 0.481 | 0.091 | 0.044 | 0.036 | -0.050 | 0.015 | 9.79E-04 |
| | rs61381027 | 8 | 26816143 | C | T | -0.038 | 0.020 | 0.059 | -0.074 | 0.052 | 0.157 | -0.004 | 0.017 | 0.826 |
| | rs6981466 | 8 | 26818171 | T | G | -0.040 | 0.020 | 0.045 | -0.072 | 0.052 | 0.163 | -0.003 | 0.017 | 0.872 |
| | rs1317819 | 8 | 26904800 | C | T | -0.010 | 0.019 | 0.600 | -0.029 | 0.048 | 0.552 | NA | NA | NA |
| | rs4732710 | 8 | 26980681 | T | C | -0.039 | 0.017 | 0.019 | -0.038 | 0.041 | 0.355 | -0.013 | 0.014 | 0.345 |
| | rs34274195 | 8 | 27168902 | A | G | 0.007 | 0.016 | 0.637 | 0.031 | 0.041 | 0.456 | -0.001 | 0.014 | 0.943 |
| | rs6999346 | 8 | 27174861 | T | G | 0.011 | 0.016 | 0.499 | 0.049 | 0.041 | 0.227 | -0.003 | 0.014 | 0.816 |
| | rs2322719 | 8 | 27184695 | A | G | 0.007 | 0.016 | 0.667 | 0.032 | 0.041 | 0.436 | -0.003 | 0.014 | 0.845 |
| | rs28834970 | 8 | 27195121 | C | T | 0.096 | 0.016 | 3.27E-09 | 0.028 | 0.043 | 0.526 | 0.044 | 0.014 | 0.001 |
| | rs73223431 | 8 | 27219987 | T | C | 0.091 | 0.016 | 1.38E-08 | 0.037 | 0.043 | 0.397 | 0.043 | 0.014 | 0.002 |
| | rs536332 | 8 | 27475767 | A | G | 0.083 | 0.016 | 2.07E-07 | 0.034 | 0.041 | 0.409 | 0.014 | 0.014 | 0.295 |
| | rs538181 | 8 | 27476815 | A | G | 0.084 | 0.016 | 2.68E-07 | 0.036 | 0.041 | 0.381 | 0.014 | 0.014 | 0.296 |
| | rs520186 | 8 | 27479103 | G | A | 0.078 | 0.016 | 6.13E-07 | 0.033 | 0.041 | 0.413 | NA | NA | NA |
| *CLU* | rs1016733 | 8 | 26592410 | T | C | 0.015 | 0.017 | 0.373 | -0.010 | 0.042 | 0.817 | 0.008 | 0.014 | 0.558 |
| | rs2036106 | 8 | 26593589 | A | C | -0.008 | 0.016 | 0.639 | 0.036 | 0.041 | 0.380 | -0.012 | 0.014 | 0.402 |
| | rs4732840 | 8 | 26593680 | A | G | -0.009 | 0.016 | 0.572 | 0.017 | 0.041 | 0.673 | -0.011 | 0.013 | 0.398 |
| | rs7813625 | 8 | 27159459 | T | C | -0.003 | 0.016 | 0.838 | 0.028 | 0.040 | 0.490 | 0.003 | 0.014 | 0.811 |
| | rs7005244 | 8 | 27168961 | A | G | -0.010 | 0.016 | 0.544 | 0.028 | 0.040 | 0.484 | 0.003 | 0.014 | 0.837 |
| | rs7819756 | 8 | 27330937 | T | C | -0.024 | 0.017 | 0.157 | -0.065 | 0.041 | 0.113 | NA | NA | NA |
| | rs2279590 | 8 | 27456253 | C | T | 0.143 | 0.017 | 3.75E-17 | 0.069 | 0.042 | 0.105 | 0.032 | 0.014 | 0.021 |
| | rs7982 | 8 | 27462481 | G | A | 0.140 | 0.017 | 2.48E-17 | 0.077 | 0.042 | 0.063 | 0.034 | 0.014 | 0.013 |
| | rs4236673 | 8 | 27464929 | G | A | 0.141 | 0.017 | 2.82E-17 | 0.072 | 0.042 | 0.086 | 0.036 | 0.014 | 0.008 |
| | rs1532277 | 8 | 27466181 | C | T | 0.139 | 0.017 | 3.06E-16 | 0.071 | 0.042 | 0.088 | 0.034 | 0.014 | 0.013 |
| | rs504038 | 8 | 27475320 | G | T | 0.074 | 0.016 | 6.91E-06 | 0.059 | 0.042 | 0.162 | 0.018 | 0.014 | 0.205 |
| | rs4732732 | 8 | 27476426 | A | G | 0.069 | 0.016 | 1.97E-05 | 0.004 | 0.041 | 0.930 | 0.017 | 0.014 | 0.212 |
| | rs538181 | 8 | 27476815 | A | G | 0.084 | 0.016 | 2.68E-07 | 0.036 | 0.041 | 0.381 | 0.014 | 0.014 | 0.296 |
| | rs4545046 | 8 | 27556526 | A | C | 0.017 | 0.016 | 0.272 | -0.074 | 0.042 | 0.080 | -0.003 | 0.014 | 0.857 |
| | rs11780158 | 8 | 27567885 | T | C | 0.006 | 0.017 | 0.725 | -0.047 | 0.041 | 0.254 | NA | NA | NA |
| | rs28692838 | 8 | 27581771 | T | C | -0.016 | 0.016 | 0.329 | 0.029 | 0.041 | 0.480 | -0.005 | 0.014 | 0.732 |
| | rs13259826 | 8 | 27582025 | C | A | -0.015 | 0.016 | 0.340 | 0.027 | 0.041 | 0.512 | -0.005 | 0.014 | 0.715 |
| | rs7465373 | 8 | 27586427 | G | A | -0.014 | 0.016 | 0.356 | 0.026 | 0.041 | 0.523 | -0.006 | 0.014 | 0.648 |
| | rs2004539 | 8 | 27587105 | T | C | -0.014 | 0.016 | 0.370 | 0.033 | 0.041 | 0.425 | -0.006 | 0.014 | 0.650 |
| | rs28460356 | 8 | 27595365 | A | G | -0.009 | 0.016 | 0.556 | 0.026 | 0.041 | 0.523 | -0.019 | 0.014 | 0.164 |
| | rs7825085 | 8 | 27598300 | T | C | -0.019 | 0.016 | 0.232 | 0.025 | 0.041 | 0.536 | -0.019 | 0.014 | 0.161 |
| | rs12334989 | 8 | 27600802 | A | C | -0.016 | 0.015 | 0.306 | 0.024 | 0.041 | 0.562 | -0.019 | 0.014 | 0.152 |
| | rs62498015 | 8 | 27603227 | T | C | -0.016 | 0.016 | 0.312 | 0.025 | 0.041 | 0.537 | -0.007 | 0.014 | 0.610 |
| | rs725361 | 8 | 27609538 | A | G | -0.017 | 0.015 | 0.276 | 0.026 | 0.041 | 0.529 | -0.019 | 0.014 | 0.164 |
| | rs10093956 | 8 | 27628886 | A | G | 0.017 | 0.016 | 0.269 | -0.016 | 0.041 | 0.686 | 0.018 | 0.014 | 0.184 |
| | rs7842666 | 8 | 27630152 | A | G | 0.016 | 0.015 | 0.313 | -0.021 | 0.041 | 0.602 | 0.018 | 0.014 | 0.186 |
| | rs11775158 | 8 | 28297929 | C | T | -0.010 | 0.034 | 0.772 | 0.009 | 0.055 | 0.867 | NA | NA | NA |
| *EXTL3* | rs7982 | 8 | 27462481 | G | A | 0.140 | 0.017 | 2.48E-17 | 0.077 | 0.042 | 0.063 | 0.034 | 0.014 | 0.013 |
| | rs17485069 | 8 | 27865907 | T | C | -0.029 | 0.016 | 0.065 | -0.003 | 0.041 | 0.941 | -0.008 | 0.014 | 0.536 |
| | rs4732800 | 8 | 27866875 | A | G | -0.025 | 0.016 | 0.109 | -0.004 | 0.041 | 0.924 | -0.008 | 0.014 | 0.564 |
| | rs6994291 | 8 | 28489068 | A | G | -0.009 | 0.019 | 0.617 | -0.081 | 0.042 | 0.055 | -0.009 | 0.014 | 0.520 |

**Supplementary Table 13. LOAD GWAS associations at SNPs that are predictors for *RAB43* expression in at least one tissue in GTEx** (IGAP: n = 54,162; ADGC2: n = 7,050; GWAX: n = 114,564; generalized Berk-Jones test).

| | | | | | IGAP | | | ADGC2 | | | GWAX | | |
|---|---|---|---|---|---|---|---|---|---|---|---|---|---|
| **SNP** | **Chr** | **BP (hg19)** | **A1** | **A2** | **Beta** | **SE** | **P** | **Beta** | **SE** | **P** | **Beta** | **SE** | **P** |
| rs6773936 | 3 | 128299708 | A | G | -0.075 | 0.021 | 4.52E-04 | 0.037 | 0.053 | 0.485 | -0.023 | 0.018 | 0.201 |
| rs61159406 | 3 | 128336009 | A | G | -0.055 | 0.016 | 5.10E-04 | 0.098 | 0.042 | 0.018 | -0.009 | 0.014 | 0.517 |
| rs13325747 | 3 | 128336023 | A | G | -0.056 | 0.016 | 4.21E-04 | 0.098 | 0.042 | 0.019 | -0.009 | 0.014 | 0.532 |
| rs7646974 | 3 | 128346861 | G | A | -0.068 | 0.018 | 1.05E-04 | 0.055 | 0.042 | 0.192 | -0.017 | 0.014 | 0.211 |
| rs13098522 | 3 | 128551055 | G | A | 0.057 | 0.021 | 0.007 | -0.048 | 0.053 | 0.362 | 0.024 | 0.017 | 0.166 |
| rs2713624 | 3 | 129282137 | C | T | -0.021 | 0.019 | 0.274 | 0.041 | 0.045 | 0.361 | 0.008 | 0.015 | 0.567 |
| rs7610060 | 3 | 129300291 | C | A | 0.020 | 0.016 | 0.216 | -0.036 | 0.043 | 0.406 | -0.009 | 0.015 | 0.525 |
| rs34703744 | 3 | 129429683 | G | A | 0.039 | 0.020 | 0.050 | -0.006 | 0.045 | 0.901 | -0.010 | 0.014 | 0.447 |
| rs6791923 | 3 | 129706392 | A | G | 0.009 | 0.016 | 0.559 | -0.006 | 0.041 | 0.880 | 0.031 | 0.014 | 0.025 |
| rs28576732 | 3 | 129765931 | T | G | NA | NA | NA | -0.057 | 0.051 | 0.259 | NA | NA | NA |
| rs13076869 | 3 | 129769381 | A | G | NA | NA | NA | -0.075 | 0.050 | 0.133 | NA | NA | NA |

Supplementary Tables 14-20 are not displayed here due to large sizes and are uploaded in separate files.

**Supplementary Table 14. LOAD GWAS associations at SNPs that are predictors for *IL10* and their weights in different tissues** (IGAP: n = 54,162; ADGC2: n = 7,050; GWAX: n = 114,564; generalized Berk-Jones test).

**Supplementary Table 15. LOAD GWAS associations at SNPs that are predictors for *CLU* and their weights in different tissues** (IGAP: n = 54,162; ADGC2: n = 7,050; GWAX: n = 114,564; generalized Berk-Jones test).

**Supplementary Table 16. LOAD GWAS associations at SNPs that are predictors for *ADRA1A* and their weights in different tissues** (IGAP: n = 54,162; ADGC2: n = 7,050; GWAX: n = 114,564; generalized Berk-Jones test).

**Supplementary Table 17. LOAD GWAS associations at SNPs that are predictors for *EXTL3* and their weights in different tissues** (IGAP: n = 54,162; ADGC2: n = 7,050; GWAX: n = 114,564; generalized Berk-Jones test).

**Supplementary Table 18. LOAD GWAS associations at SNPs that are predictors for *RAB43* and their weights in different tissues** (IGAP: n = 54,162; ADGC2: n = 7,050; GWAX: n = 114,564; generalized Berk-Jones test).

**Supplementary Table 19. R2 in prediction step for *IL10, CLU, ADRA1A, EXTL3* and *RAB43* in different tissues. NA indicates no prediction model available for that tissue.**

**Supplementary Table 20. Single tissue association in LOAD for *IL10, CLU, ADRA1A, EXTL3* and *RAB43.*** (IGAP: n = 54,162; ADGC2: n = 7,050; GWAX: n = 114,564; z-score test, two-sided).

**Supplementary Table 21. Matched tissues between GTEx and GenoSkyline-Plus annotation tracks.**

| GTEx Tissues | Roadmap Epigenome IDs | Roadmap Epigenome Tissues |
|---|---|---|
| Adipose_Visceral_Omentum | E063 | Adipose_Nuclei |
| Adipose_Subcutaneous | E063 | Adipose_Nuclei |
| Liver | E066 | Adult_Liver |
| Artery_Aorta | E065 | Aorta |
| Brain_Cortex | E067 | Brain_Angular_Gyrus |
| Brain_Caudate_basal_ganglia | E068 | Brain_Anterior_Caudate |
| Brain_Anterior_cingulate_cortex_BA 24 | E069 | Brain_Cingulate_Gyrus |
| Brain_Hippocampus | E071 | Brain_Hippocampus_Middle |
| Brain_Cortex | E072 | Brain_Inferior_Temporal_Lobe |
| Brain_Cortex | E073 | Brain_Mid_Frontal_Lobe |
| Brain_Frontal_Cortex_BA9 | E072 | Brain_Inferior_Temporal_Lobe |
| Brain_Frontal_Cortex_BA9 | E073 | Brain_Mid_Frontal_Lobe |
| Breast_Mammary_Tissue | E027 | Breast_Myoepithelial_Cells |
| Colon_Transverse | E075 | Colonic_Mucosa |
| Colon_Transverse | E076 | Colon_Smooth_Muscle |
| Esophagus_Gastroesophageal_Junction | E079 | Esophagus |
| Esophagus_Mucosa | E079 | Esophagus |
| Esophagus_Muscularis | E079 | Esophagus |
| Cells_EBV-transformed_lymphocytes | E116 | GM12878_Lymphoblastoid |
| Heart_Left_Ventricle | E095 | Left_Ventricle |
| Lung | E096 | Lung |
| Ovary | E097 | Ovary |
| Pancreas | E098 | Pancreas |
| Whole_Blood | E062 | Peripheral_Blood_Mononuclear_Primary_Cells |
| Heart_Atrial_Appendage | E104 | Right_Atrium |
| Colon_Sigmoid | E106 | Sigmoid_Colon |
| Muscle_Skeletal | E108 | Skeletal_Muscle_Female |
| Muscle_Skeletal | E107 | Skeletal_Muscle_Male |
| Small_Intestine_Terminal_Ileum | E109 | Small_Intestine |
| Spleen | E113 | Spleen |
| Stomach | E110 | Stomach_Mucosa |
| Stomach | E111 | Stomach_Smooth_Muscle |

**Supplementary Table 22. Significant genes identified in BLUEPRINT data.** Bonferroni-corrected significance level is 0.05/4758=1.05e-5 , n = 54,162, z-score test, two-sided.

| Gene | z-score | Effect size | p-value |
|---|---|---|---|
| *NDUFS3* | -5.093 | -0.001 | 3.51E-07 |
| *SNRPD2* | 4.804 | 0.008 | 1.55E-06 |
| *HLA-DOB* | -4.581 | -0.012 | 4.61E-06 |
| *DMWD* | 4.478 | 0.002 | 7.51E-06 |
| *PTK2B* | -4.477 | -0.150 | 7.55E-06 |
| *ZNF232* | -4.421 | -7.4E-04 | 9.80E-06 |
| *MTCH2* | 4.418 | 0.001 | 9.93E-06 |

**Supplementary Table 23. Significant genes identified in ImmVar data.** Bonferroni-corrected significance level is 0.05/2632=1.90e-5, n = 54,162, z-score test, two-sided.

| Gene | z-score | Effect size | p-value |
|---|---|---|---|
| *FIS1* | -36.387 | -628.6 | 6.77E-290 |
| *MRPS21* | -8.286 | 298.9 | 1.16E-16 |
| *MS4A4A* | 6.614 | 0.040 | 3.74E-11 |
| *TAS2R41* | -5.845 | -0.022 | 5.07E-09 |
| *TAS2R60* | -5.840 | -0.020 | 5.23E-09 |
| *KEAP1* | 4.842 | 66.7 | 1.28E-06 |
| *SNX11* | -4.483 | -2.815 | 7.35E-06 |
| *NUP160* | 4.458 | 0.007 | 8.26E-06 |
| *CHRNE* | 4.335 | 0.036 | 1.46E-05 |
| *SLC39A13* | 4.302 | 0.247 | 1.69E-05 |

Supplementary Tables 24-25 are not displayed here due to large sizes and are uploaded in separate files.

**Supplementary Table 24. Most significantly and non-significantly enriched tissue in LDSC with GenoSkyline-Plus annotations and the number of genes identified by TWAS, PrediXcan and UTMOST** (enrichment of heritability tested by LD score regression).

**Supplementary Table 25. Number of genes identified in single-tissue association test in 50 complex traits, z-score test, two-sided.**

**Supplementary Table 26. Number of significant associations in 44 tissues identified in IGAP** (IGAP: n = 54,162; z-score test, two-sided).

| tissue | num_sig |
|---|---|
| Adipose_Subcutaneous | 3 |
| Adipose_Visceral_Omentum | 5 |
| Adrenal_Gland | 8 |
| Artery_Aorta | 8 |
| Artery_Coronary | 7 |
| Artery_Tibial | 6 |
| Brain_Anterior_cingulate_cortex_BA24 | 5 |
| Brain_Caudate_basal_ganglia | 7 |
| Brain_Cerebellar_Hemisphere | 7 |
| Brain_Cerebellum | 13 |
| Brain_Cortex | 8 |
| Brain_Frontal_Cortex_BA9 | 8 |
| Brain_Hippocampus | 5 |
| Brain_Hypothalamus | 9 |
| Brain_Nucleus_accumbens_basal_ganglia | 7 |
| Brain_Putamen_basal_ganglia | 8 |
| Breast_Mammary_Tissue | 8 |
| Cells_EBV-transformed_lymphocytes | 5 |
| Cells_Transformed_fibroblasts | 3 |
| Colon_Sigmoid | 8 |
| Colon_Transverse | 5 |
| Esophagus_Gastroesophageal_Junction | 5 |
| Esophagus_Mucosa | 5 |
| Esophagus_Muscularis | 9 |
| Heart_Atrial_Appendage | 7 |
| Heart_Left_Ventricle | 6 |
| Liver_STARNET | 5 |
| Liver | 6 |
| Lung | 6 |
| mono_eqtl | 6 |
| mono_sqtl | 3 |
| Muscle_Skeletal | 6 |
| Nerve_Tibial | 5 |
| neut_eqtl | 4 |
| neut_sqtl | 0 |
| Ovary | 9 |
| Pancreas | 5 |
| Pituitary | 10 |
| Prostate | 9 |
| Skin_Not_Sun_Exposed_Suprapubic | 8 |
| Skin_Sun_Exposed_Lower_leg | 9 |
| Small_Intestine_Terminal_Ileum | 11 |
| Spleen | 8 |
| Stomach | 5 |
| tcel_eqtl | 0 |
| tcel_sqtl | 0 |
| Testis | 11 |
| Thyroid | 9 |
| Uterus | 5 |
| Vagina | 10 |
| Whole_Blood | 9 |

**Supplementary Table 27. eQTL effect size estimates in single tissue model and cross-tissue model of *LNT1*.**

| SNP | A1 | A2 | Effect |
|---|---|---|---|
| **Single-tissue elastic net** | | | |
| 3_100786402_G_T_b37 | G | T | -0.001670313 |
| 3_100797007_G_A_b37 | G | A | -0.000103896 |
| 3_100798050_G_T_b37 | G | T | -0.000516352 |
| **UTMOST** | | | |
| 3_100053083_G_A_b37 | G | A | -5.72E-05 |
| 3_100055273_G_A_b37 | G | A | -1.00E-03 |
| 3_100057203_A_G_b37 | A | G | -4.23E-05 |
| 3_100073826_T_G_b37 | T | G | -4.13E-04 |
| 3_100076980_A_G_b37 | A | G | -2.20E-05 |
| 3_100085854_G_A_b37 | G | A | -6.31E-05 |
| 3_100086477_G_A_b37 | G | A | -2.69E-04 |
| 3_100099142_C_T_b37 | C | T | -2.98E-06 |
| 3_100104114_C_T_b37 | C | T | -5.50E-06 |
| 3_100106412_C_T_b37 | C | T | -8.89E-05 |
| 3_100112168_C_T_b37 | C | T | -6.36E-05 |
| 3_100112816_C_T_b37 | C | T | -6.24E-05 |
| 3_100114263_A_G_b37 | A | G | -6.31E-05 |
| 3_100120850_G_A_b37 | G | A | -5.83E-05 |
| 3_100121121_G_A_b37 | G | A | -5.55E-05 |
| 3_100122081_C_T_b37 | C | T | -5.54E-05 |
| 3_100131897_A_G_b37 | A | G | -1.34E-05 |
| 3_100135624_T_C_b37 | T | C | -1.29E-04 |
| 3_100136644_A_G_b37 | A | G | -3.31E-05 |
| 3_100141714_A_C_b37 | A | C | -4.54E-05 |
| 3_100169270_T_C_b37 | T | C | -1.34E-04 |
| 3_100170825_G_A_b37 | G | A | -3.12E-05 |
| 3_100173525_C_T_b37 | C | T | -2.55E-05 |
| 3_100174722_G_A_b37 | G | A | -4.37E-05 |
| 3_100176360_G_A_b37 | G | A | -4.75E-05 |
| 3_100178459_G_A_b37 | G | A | -1.53E-04 |
| 3_100181868_G_A_b37 | G | A | -1.94E-05 |
| 3_100258358_C_T_b37 | C | T | 1.14E-04 |
| 3_100556348_A_G_b37 | A | G | -9.42E-06 |

**Supplementary Table 28. Significant genes for LDL cholesterol based on STARNET** (n = 173,082, z-score test, two-sided, p-value threshold = 8.39E-6).

| Gene | z-score | Effect size | p-value |
|---|---|---|---|
| *CELSR2* | 46.201 | 0.919 | 0 |
| *PSRC1* | 44.867 | 0.858 | 0 |
| *SORT1* | 41.493 | 0.666 | 0 |
| *APOC1* | 22.932 | 0.629 | 2.19E-116 |
| *PSMA5* | 19.995 | 0.465 | 5.97E-89 |
| *FEN1* | 13.041 | 0.436 | 7.07E-39 |
| *FADS1* | 12.721 | 0.130 | 4.48E-37 |
| *FADS3* | 12.546 | 0.148 | 4.13E-36 |
| *CETP* | 11.349 | 0.280 | 7.49E-30 |
| *FADS2* | 10.439 | 0.227 | 1.64E-25 |
| *ANGPTL3* | 10.266 | 0.529 | 9.94E-25 |
| *ST3GAL4* | 10.112 | 0.287 | 4.88E-24 |
| *RP11-115J16.1* | 9.894 | 0.134 | 4.38E-23 |
| *LPIN3* | 8.943 | 0.139 | 3.76E-19 |
| *APOC1P1* | -8.927 | -0.097 | 4.38E-19 |
| *HPR* | -8.860 | -0.070 | 7.95E-19 |
| *ZHX3* | 8.473 | 0.834 | 2.38E-17 |
| *MYLIP* | 8.002 | 0.420 | 1.22E-15 |
| *RP11-10A14.4* | -7.938 | -0.155 | 2.05E-15 |
| *AC079602.1* | 7.820 | 0.264 | 5.27E-15 |
| *ULK3* | -7.717 | -0.368 | 1.18E-14 |
| *ZNF229* | 7.399 | 0.226 | 1.37E-13 |
| *DDX56* | 7.250 | 0.164 | 4.15E-13 |
| *LPA* | 7.165 | 0.063 | 7.77E-13 |
| *TNKS* | 7.124 | 0.077 | 1.04E-12 |
| *SLC22A1* | -7.079 | -0.099 | 1.45E-12 |
| *PLCG1* | 6.828 | 0.148 | 8.60E-12 |
| *TMED4* | 6.753 | 0.069 | 1.44E-11 |
| *EVI5* | -6.676 | -0.090 | 2.45E-11 |
| *TAF13* | 6.561 | 0.258 | 5.33E-11 |
| *LPAL2* | 6.543 | 0.231 | 6.00E-11 |
| *RHCE* | 6.429 | 0.047 | 1.28E-10 |
| *IRF2BP2* | 6.323 | 0.314 | 2.56E-10 |
| *ZDHHC18* | 6.264 | 0.146 | 3.75E-10 |
| *EFCAB13* | 6.231 | 0.055 | 4.63E-10 |
| *AMIGO1* | -6.059 | -0.034 | 1.36E-09 |
| *CTB-171A8.1* | 5.943 | 0.171 | 2.79E-09 |

| | | | |
|---|---|---|---|
| *RP4-781K5.7* | -5.915 | -0.077 | 3.30E-09 |
| *TMEM50A* | -5.646 | -0.145 | 1.64E-08 |
| *MICA* | -5.589 | -0.047 | 2.28E-08 |
| *RGL3* | 5.544 | 0.121 | 2.95E-08 |
| *CEACAM19* | 5.532 | 0.153 | 3.16E-08 |
| *NUDCD3* | 5.496 | 0.094 | 3.88E-08 |
| *EHBP1* | -5.463 | -0.157 | 4.67E-08 |
| *KPNB1* | -5.420 | -0.089 | 5.94E-08 |
| *NT5DC1* | -5.327 | -0.143 | 9.98E-08 |
| *FRK* | 5.286 | 0.111 | 1.25E-07 |

**Supplementary Table 29. Conditional analysis results in *SORT1* loci for cardio-vascular disease (CAD).** n = 86,995, z-score test in conditional analysis, two-sided.

| Gene | p-value |
|---|---|
| *AKNAD1* | 5.44E-01 |
| *AMIGO1* | 2.70E-01 |
| *AMPD2* | 7.75E-01 |
| *ATXN7L2* | 5.38E-02 |
| *C1orf194* | 1.88E-02 |
| *CELSR2* | 5.57E-06 |
| *CLCC1* | 3.80E-01 |
| *CYB561D1* | 1.09E-01 |
| *EPS8L3* | 1.86E-01 |
| *GNAT2* | 4.93E-01 |
| *GPR61* | 1.19E-02 |
| *GPSM2* | 7.68E-01 |
| *GSTM1* | 5.52E-01 |
| *GSTM2* | 1.00E+00 |
| *GSTM3* | 3.58E-01 |
| *GSTM4* | 5.67E-02 |
| *GSTM5* | 3.20E-01 |
| *KIAA1324* | 1.56E-02 |
| *MYBPHL* | 5.57E-02 |
| *PSMA5* | 2.12E-01 |
| *PSRC1* | 1.60E-11 |
| *SARS* | 2.50E-02 |
| *SORT1* | 9.87E-11 |
| *SYPL2* | 4.44E-01 |
| *TAF13* | 1.00E+00 |
| *TMEM167B* | 6.45E-01 |
| *WDR47* | 2.27E-12 |

### Supplementary Figures

**Supplementary Figure 1. Average improvement in squared Spearman correlation.** Sample sizes of 44 GTEx tissues are listed in Supplementary Table 1**.**

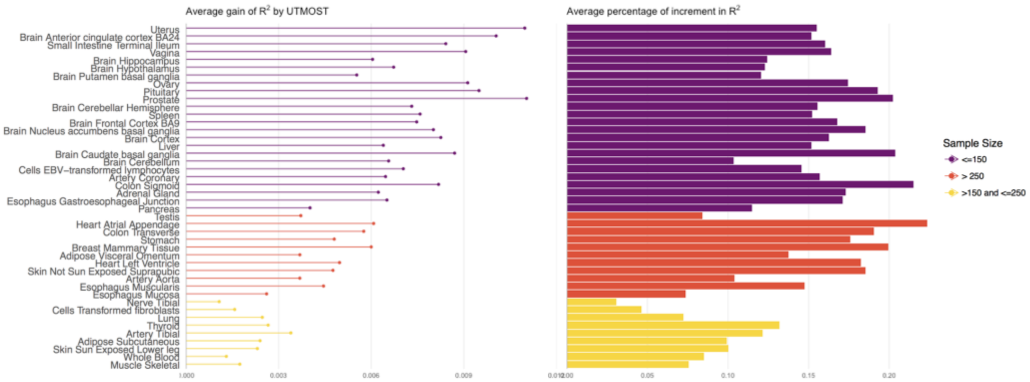

**Supplementary Figure 2**. **Improvement in gene expression imputation accuracy compared to BSLMM trained on each tissue**. UTMOST showed substantially higher imputation accuracy (A, B). The improvement is higher in tissues with smaller sample sizes. Sample sizes of 44 GTEx tissues are listed in Supplementary Table 1.

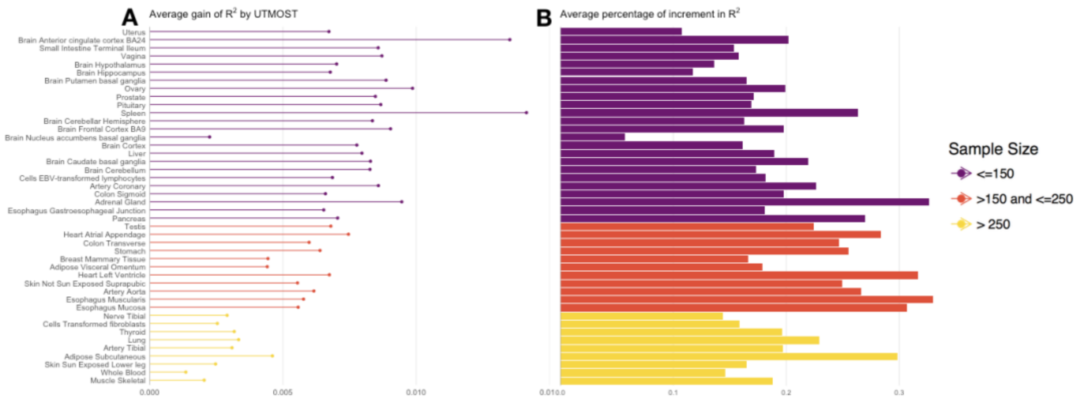

**Supplementary Figure 3. External validation of imputation accuracy using the GEUVADIS LCL data.** Multi-tissue was trained on 44 GTEx tissues while the single-tissue elastic net model was trained only on the whole blood tissue from GTEx. Both models were used to predict gene expression levels in the GEUVADIS LDL dataset. The prediction $R^2$ of 4,141 genes, of which the effect size estimates are nonzero by both models, were compared against the null case where no SNP is predictive of the expression. Both models showed significant deviation from the null case and multi-tissue joint prediction significantly outperformed the single-tissue elastic net (Kolmogorov–Smirnov test, one-sided).

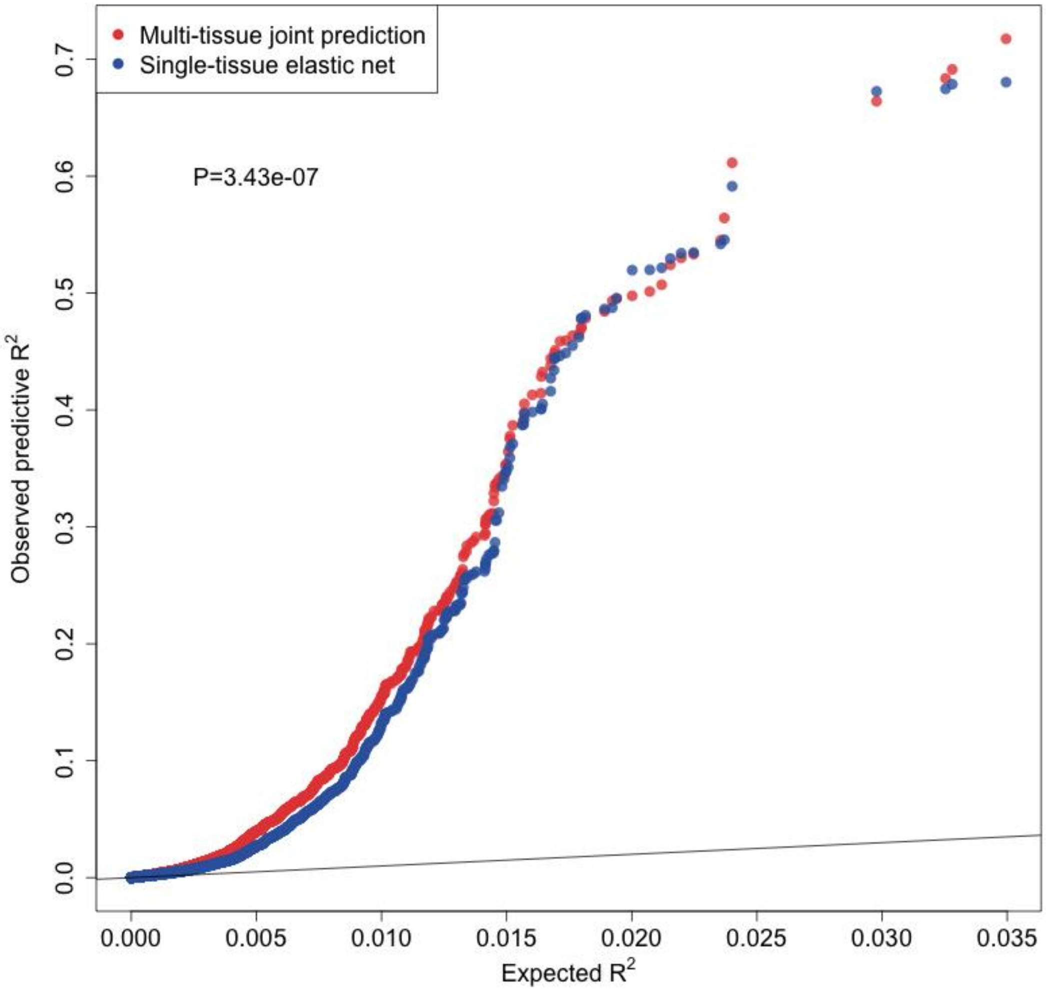

**Supplementary Figure 4. External validation of imputation accuracy using the GEUVADIS LCL and CommonMind data**. Multi-tissue model was trained on 44 GTEx tissues while the single-tissue elastic net model was trained only on Whole Blood or Brain Frontal Cortex BA9 tissue from GTEx. Both models were used to predict gene expression levels in GEUVADIS LCL (Whole Blood) and CommonMind (Brain Frontal Cortex BA9) datasets. UTMOST showed improved performance in both external validation data.

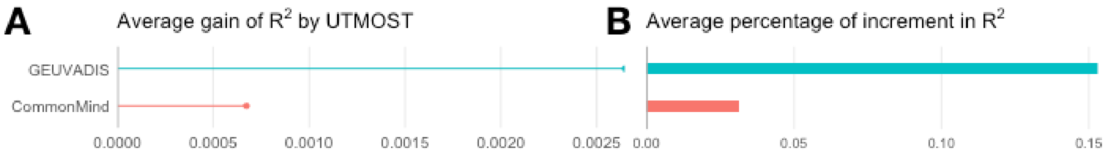

**Supplementary Figure 5. Similarity matrix between tissues calculated by** (A) observed gene expression levels and (B) imputed gene expression levels. Tissue-tissue similarity was measured by the averaged absolute correlation of gene expressions between tissues. Similarities between tissues with no gene expression measured on shared individuals were denoted as missing (grey boxes in the figure).

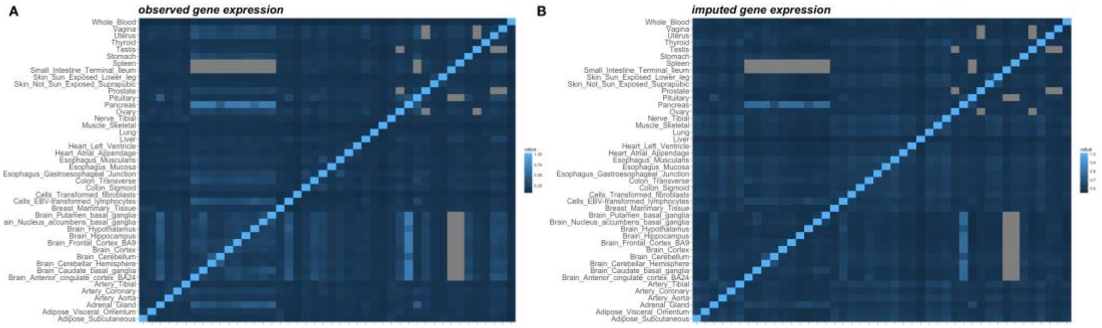

**Supplementary Figure 6. Box plots of within-cluster and between-cluster correlations.** Within-cluster: correlation between each pair of 10 brain tissues (n = 45); Between-cluster correlations between 10 brain tissues and the rest 34 tissues (n = 340). (**A**) Pearson correlation calculated with imputed gene expression levels; (**B**) Pearson correlation calculated with observed gene expression levels. In each box, the two horizontal borders represent the upper and lower quartiles, solid line in the middle represent median. The highest and lowest points indicate the maxima and minima.

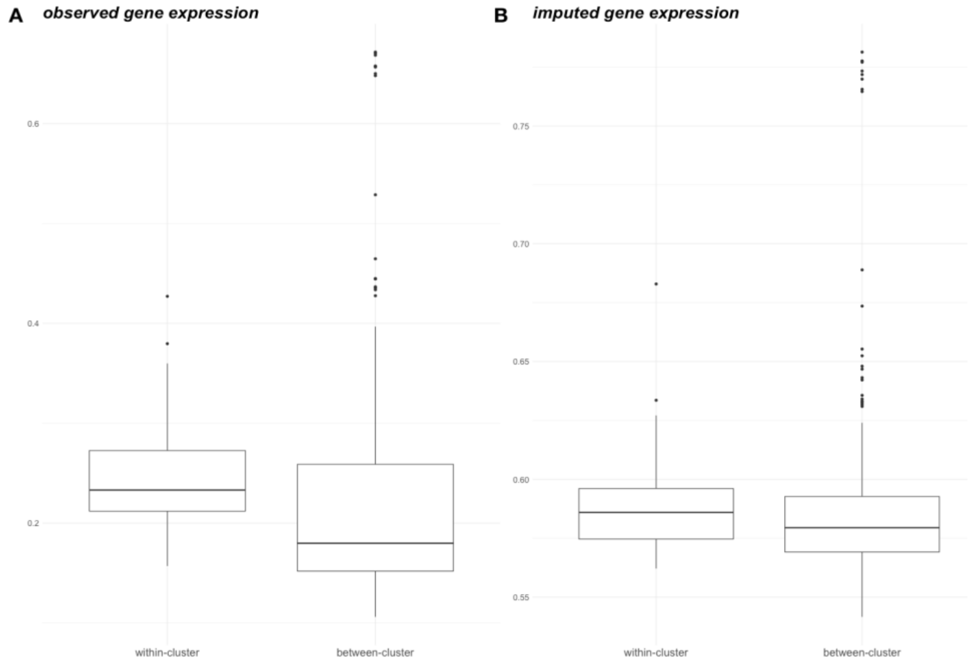

**Supplementary Figure 7. Number of significant genes identified in UTMOST single-tissue tests and cross-tissue joint test.** Each point represents one of 50 analyzed complex traits. Values on both axes show log-transformed number of significant genes. Overall, UTMOST joint test identified more genes than single-tissue tests combined. P-value shown on the figure was calculated via one-sided paired-Wilcox Rank test, n = 50.

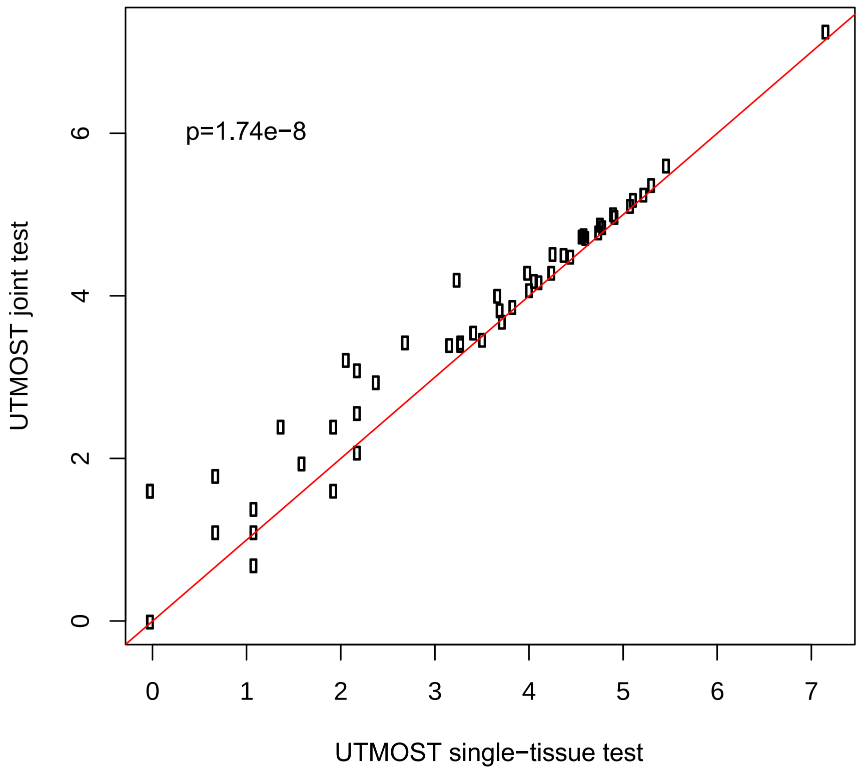

**Supplementary Figure 8. Manhattan plot for the discovery stage of LOAD association analysis.** The horizontal line indicates the genome-wide significance threshold based on Bonferroni correction (n = 54,162, z-score test, two-sided).

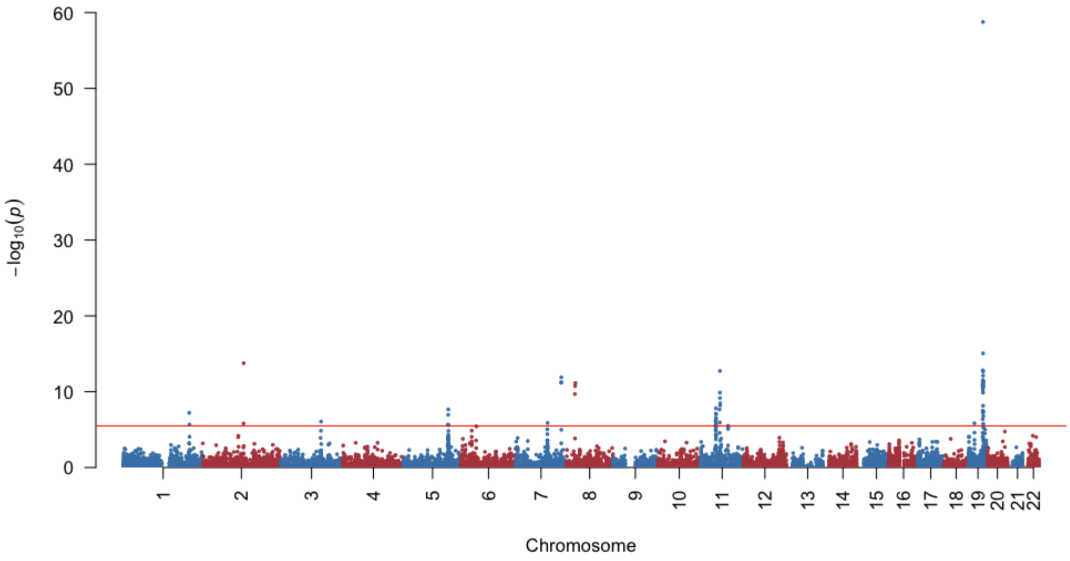

**Supplementary Figure 9. GWAS associations at the *IL10-CR1* locus in three LOAD datasets.** Panels A-C show SNP-level associations in IGAP, ADGC, and GWAX data, respectively (IGAP: n = 54,162; ADGC2: n = 7,050; GWAX: n = 114,564, two-sided z-score test).

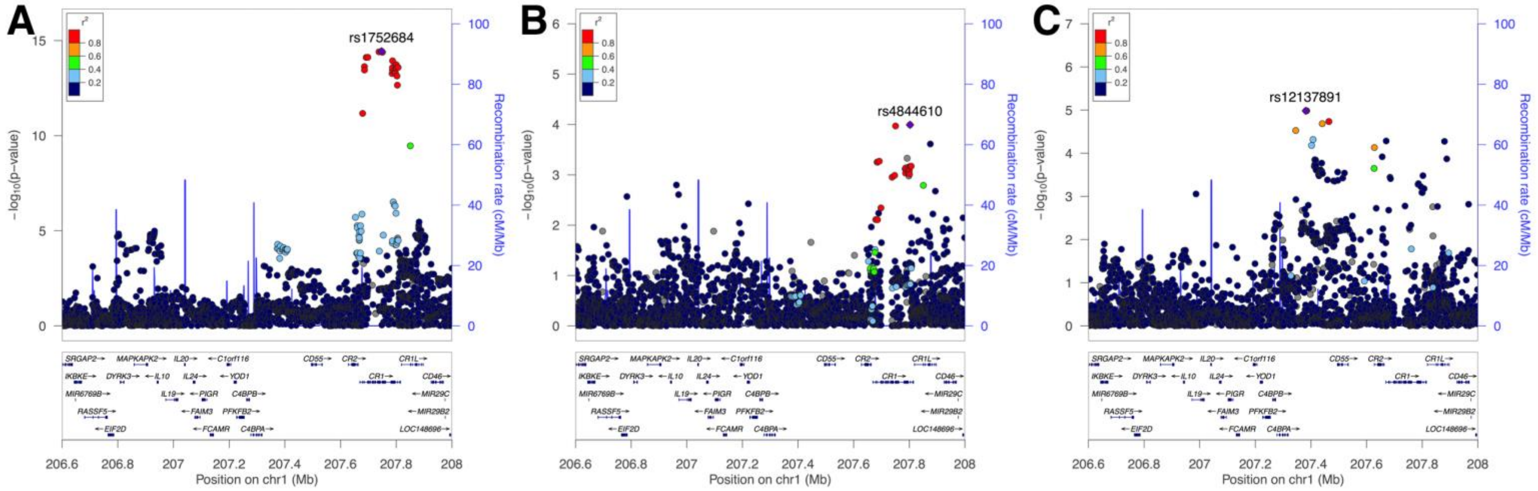

**Supplementary Figure 10. GWAS associations at the *CLU-PTK2B* locus in three LOAD datasets.** Panels A-C show SNP-level associations in IGAP, ADGC, and GWAX data, respectively (IGAP: n = 54,162; ADGC2: n = 7,050; GWAX: n = 114,564, two-sided z-score test).

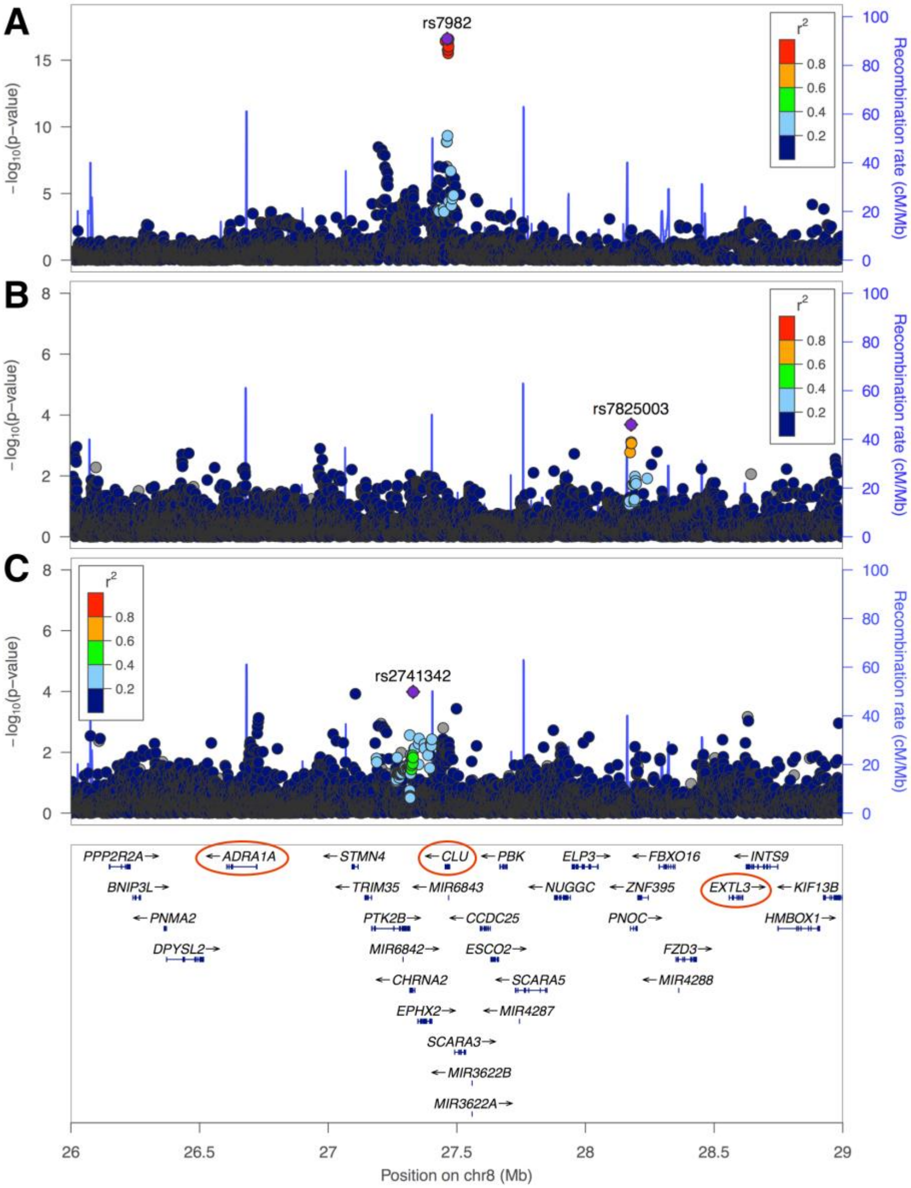

**Supplementary Figure 11. GWAS associations at the *RAB43-RPN1* locus in three LOAD datasets.** Panels A-C show SNP-level associations in IGAP, ADGC, and GWAX data, respectively (IGAP: n = 54,162; ADGC2: n = 7,050; GWAX: n = 114,564, two-sided z-score test).

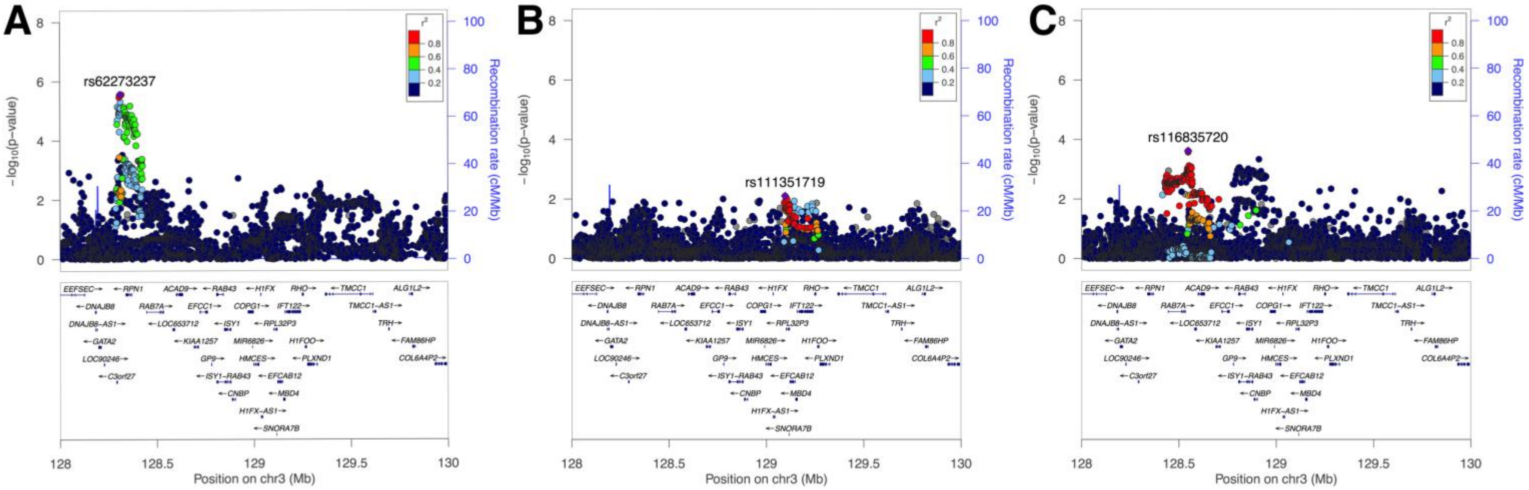

**Supplementary Figure 12. Associations between simulated traits and genetically regulated gene expression levels in the causal and non-causal tissues.** Different panels indicate results based on different proportions of shared eQTLs (ranging from 0 to 1, indicated by the title of each panel). Red horizontal lines indicate the significance threshold after adjusting for the number of tissues (i.e. $-\log_{10}(0.05/2)$). Each box represents –log(p) of 100 replicates for a setting. In each box, the two horizontal borders represent the upper and lower quartiles, solid line in the middle represent median. The highest and lowest points indicate the maxima and minima.

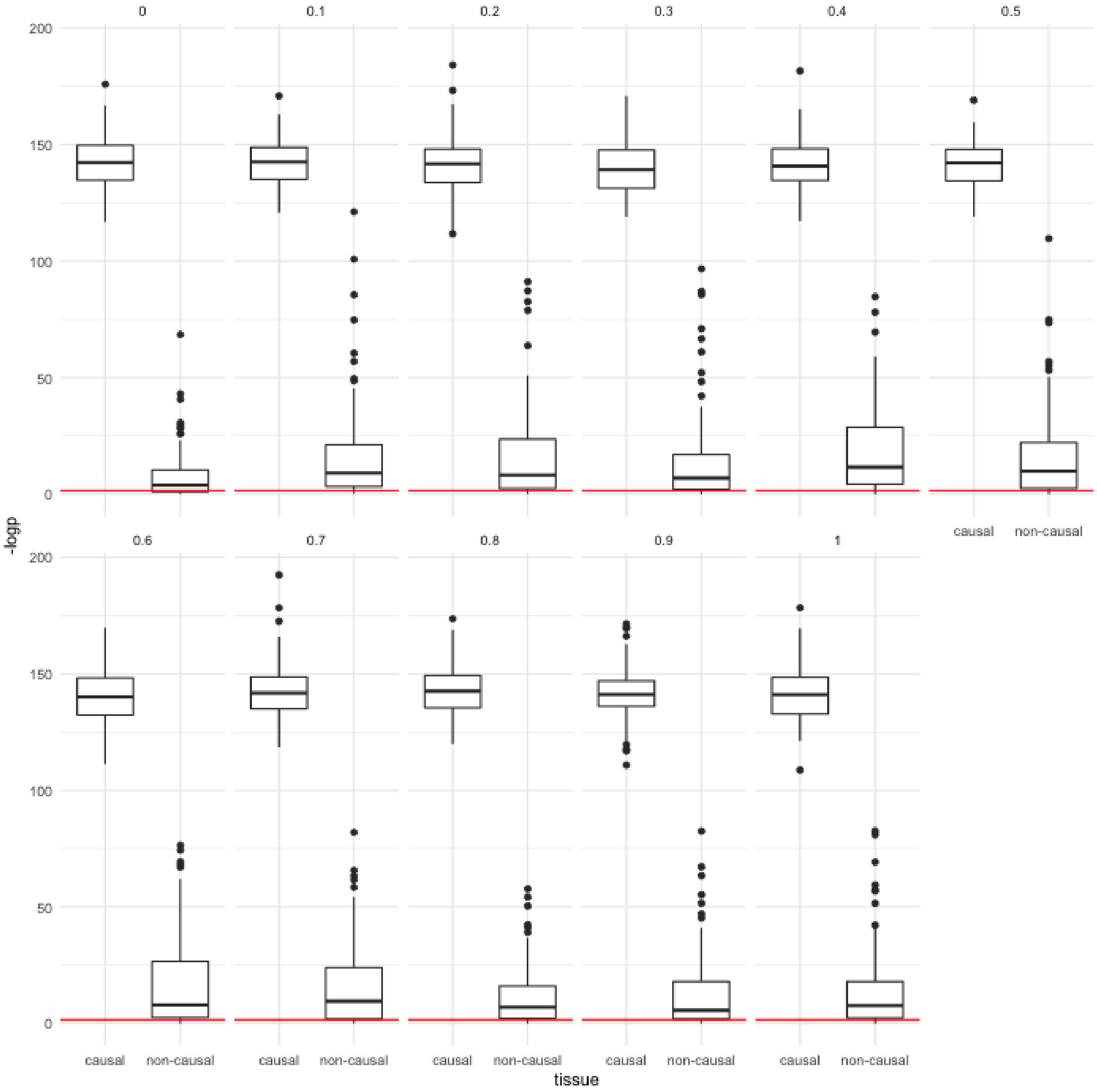

**Supplementary Figure 13**. Associations at *SORT1* locus in the single-tissue analysis based on STARNET data. The horizontal line indicates the Bonferroni-corrected genome-wide significance threshold (n=173,082, two-sided z-score test).

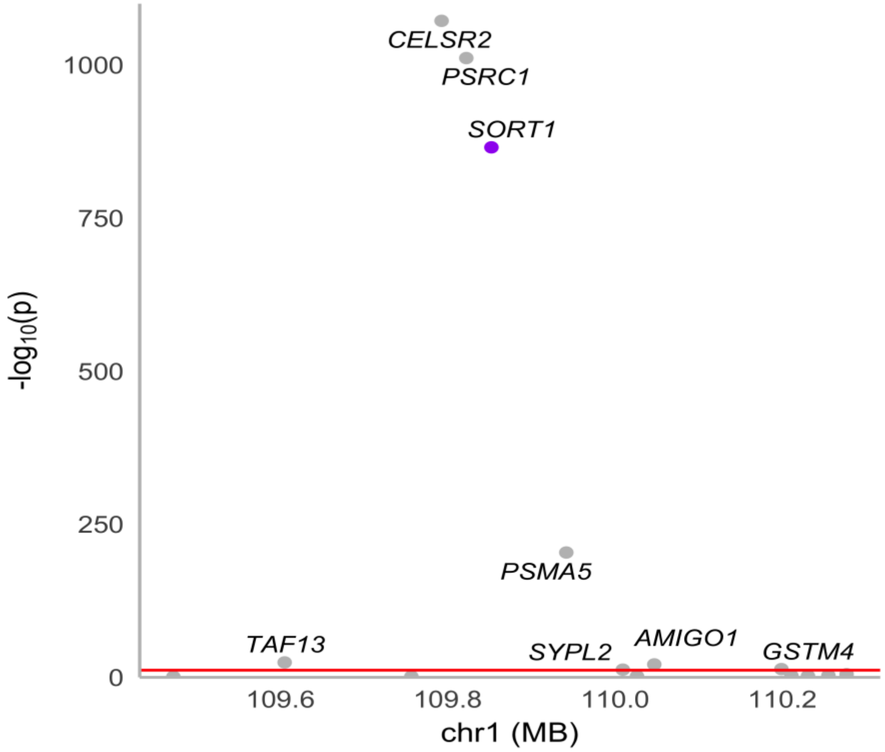

#### Acknowledgements to ADGC

The National Institutes of Health, National Institute on Aging (NIH-NIA) supported this work through the following grants: ADGC, U01 AG032984, RC2 AG036528; Samples from the National Cell Repository for Alzheimer's Disease (NCRAD), which receives government support under a cooperative agreement grant (U24 AG21886) awarded by the National Institute on Aging (NIA), were used in this study. We thank contributors who collected samples used in this study, as well as patients and their families, whose help and participation made this work possible; Data for this study were prepared, archived, and distributed by the National Institute on Aging Alzheimer's Disease Data Storage Site (NIAGADS) at the University of Pennsylvania (U24-AG041689-01); NACC, U01 AG016976; NIA LOAD (Columbia University), U24 AG026395, U24 AG026390, R01AG041797; Banner Sun Health Research Institute P30 AG019610; Boston University, P30 AG013846, U01 AG10483, R01 CA129769, R01 MH080295, R01 AG017173, R01 AG025259, R01 AG048927, R01AG33193, R01 AG009029; Columbia University, P50 AG008702, R37 AG015473, R01 AG037212, R01 AG028786; Duke University, P30 AG028377, AG05128; Emory University, AG025688; Group Health Research Institute, UO1 AG006781, UO1 HG004610, UO1 HG006375, U01 HG008657; Indiana University, P30 AG10133, R01 AG009956, RC2 AG036650; Johns Hopkins University, P50 AG005146, R01 AG020688; Massachusetts General Hospital, P50 AG005134; Mayo Clinic, P50 AG016574, R01 AG032990, KL2 RR024151; Mount Sinai School of Medicine, P50 AG005138, P01 AG002219; New York University, P30 AG08051, UL1 RR029893, 5R01AG012101, 5R01AG022374, 5R01AG013616, 1RC2AG036502, 1R01AG035137; North Carolina A&T University, P20 MD000546, R01 AG28786-01A1; Northwestern University, P30 AG013854; Oregon Health & Science University, P30 AG008017, R01 AG026916; Rush University, P30 AG010161, R01 AG019085, R01 AG15819, R01 AG17917, R01 AG030146, R01 AG01101, RC2 AG036650, R01 AG22018; TGen, R01 NS059873; University of Alabama at Birmingham, P50 AG016582; University of Arizona, R01 AG031581; University of California, Davis, P30 AG010129; University of California, Irvine, P50 AG016573; University of California, Los Angeles, P50 AG016570; University of California, San Diego, P50 AG005131; University of California, San Francisco, P50 AG023501, P01 AG019724; University of Kentucky, P30 AG028383, AG05144; University of Michigan, P50 AG008671; University of Pennsylvania, P30 AG010124; University of Pittsburgh, P50 AG005133, AG030653, AG041718, AG07562, AG02365; University of Southern California, P50 AG005142; University of Texas Southwestern, P30 AG012300; University of Miami, R01 AG027944, AG010491, AG027944, AG021547, AG019757; University of Washington, P50 AG005136, R01 AG042437; University of Wisconsin, P50 AG033514; Vanderbilt University, R01 AG019085; and Washington University, P50 AG005681, P01 AG03991, P01 AG026276. The Kathleen Price Bryan Brain Bank at Duke University Medical Center is funded by NINDS grant # NS39764, NIMH MH60451 and by Glaxo Smith Kline. Support was also from the Alzheimer's Association (LAF, IIRG-08-89720; MP-V, IIRG-05-14147), the US Department of Veterans Affairs Administration, Office of Research and Development, Biomedical Laboratory Research Program, and BrightFocus Foundation (MP-V, A2111048). P.S.G.-H. is supported by Wellcome Trust, Howard Hughes Medical Institute, and the Canadian Institute of Health Research. Genotyping of the TGEN2 cohort was supported by Kronos Science. The TGen series was also funded by NIA grant AG041232 to AJM and MJH, The Banner Alzheimer's Foundation, The Johnnie B. Byrd Sr. Alzheimer's Institute, the Medical Research Council, and the state of Arizona and also includes samples from the following sites: Newcastle Brain Tissue Resource (funding via the Medical Research Council, local NHS trusts and Newcastle University), MRC London Brain Bank for Neurodegenerative Diseases (funding via the Medical Research Council),South West Dementia Brain Bank (funding via numerous sources including the Higher Education Funding Council for England (HEFCE), Alzheimer's Research Trust (ART), BRACE as well as North Bristol NHS Trust Research and Innovation Department and DeNDRoN), The Netherlands Brain Bank (funding via numerous sources including Stichting MS Research, Brain Net Europe, Hersenstichting Nederland Breinbrekend Werk, International Parkinson Fonds, Internationale Stiching Alzheimer Onderzoek), Institut de Neuropatologia, Servei Anatomia Patologica, Universitat de Barcelona. ADNI data collection and sharing was funded by the National Institutes of Health Grant U01 AG024904 and Department of Defense award number W81XWH-12-2-0012. ADNI is funded by the National Institute on Aging, the National Institute of Biomedical Imaging and Bioengineering, and through generous contributions from the following: AbbVie, Alzheimer's Association; Alzheimer's Drug Discovery Foundation; Araclon Biotech; BioClinica, Inc.; Biogen; Bristol-Myers Squibb Company; CereSpir, Inc.; Eisai Inc.; Elan Pharmaceuticals, Inc.; Eli Lilly and Company; EuroImmun; F. Hoffmann-La Roche Ltd and its affiliated company Genentech, Inc.; Fujirebio; GE Healthcare; IXICO Ltd.; Janssen Alzheimer Immunotherapy Research & Development, LLC.; Johnson & Johnson Pharmaceutical Research & Development LLC.; Lumosity; Lundbeck; Merck & Co., Inc.; Meso Scale Diagnostics, LLC.; NeuroRx Research; Neurotrack Technologies; Novartis Pharmaceuticals Corporation; Pfizer Inc.; Piramal Imaging; Servier; Takeda Pharmaceutical Company; and Transition Therapeutics. The Canadian Institutes of Health Research is providing funds to support ADNI clinical sites in Canada. Private sector contributions are facilitated by the Foundation for the National Institutes of Health (www.fnih.org). The grantee organization is the Northern California Institute for Research and Education, and the study is coordinated by the Alzheimer's Disease Cooperative Study at the University of California, San Diego. ADNI data are disseminated by the Laboratory for Neuro Imaging at the University of Southern California. We thank Drs. D. Stephen Snyder and Marilyn Miller from NIA who are *ex-officio* ADGC members.

We would also like to thank all the members of ADGC: Erin Abner [1], Perrie M. Adams [2], Marilyn S. Albert [3], Roger L. Albin [4-,6], Liana G. Apostolova [7-10], Steven E. Arnold [11], Sanjay Asthana [12-14], Craig S. Atwood [12-14], Clinton T. Baldwin [15], Robert C. Barber [16], Lisa L. Barnes [17-19], Sandra Barral [20-22], Thomas G. Beach [23], James T. Becker [24], Gary W. Beecham [25,26], Duane Beekly [27], David A. Bennett [17,19], Eileen H. Bigio [28,29], Thomas D. Bird [30,31], Deborah Blacker [32,33], Bradley F. Boeve [34], James D. Bowen [35], Adam Boxer [36], James R. Burke [37], Jeffrey M. Burns [38], Joseph D. Buxbaum [39-41], Nigel J. Cairns [42], Laura B. Cantwell [43], Chuanhai Cao [44], Chris S. Carlson [45], Cynthia M. Carlsson [12-14], Regina M. Carney [46], Minerva M. Carrasquillo [47], Helena C. Chui [48], Paul K. Crane [49], David H. Cribbs [50], Elizabeth A. Crocco [46], Carlos Cruchaga [51], Philip L. De Jager [52,53], Charles DeCarli [54], Malcolm Dick [55], Dennis W. Dickson [47], Rachelle S. Doody [56], Ranjan Duara [57], Nilufer Ertekin-Taner [47,58], Denis A. Evans [59], Kelley M. Faber [8], Thomas J. Fairchild [60], Kenneth B. Fallon [61], David W. Fardo [62], Martin R. Farlow [63], Lindsay A. Farrer [64-68], Steven Ferris [69], Tatiana M. Foroud [8], Matthew P. Frosch [70], Douglas R. Galasko [71], Marla

Gearing [72,73], Daniel H. Geschwind [74], Bernardino Ghetti [75], John R. Gilbert [25,26], Alison M. Goate [39], Neill R. Graff-Radford [47,58], Robert C. Green [76], John H. Growdon [77], Jonathan L. Haines [78], Hakon Hakonarson [79], Ronald L. Hamilton [80], Kara L. Hamilton-Nelson [25], John Hardy [81], Lindy E. Harrell [82], Lawrence S. Honig [20], Ryan M. Huebinger [83], Matthew J. Huentelman [84], Christine M. Hulette [85], Bradley T. Hyman [77], Gail P. Jarvik [86,87], Lee-Way Jin [88], Gyungah Jun [15,64,68], M. Ilyas Kamboh [89,90], Anna Karydas [36], Mindy J. Katz [91], John S.K. Kauwe [92], Jeffrey A. Kaye [93,94], C. Dirk Keene [95], Ronald Kim [96], Neil W. Kowall[67,97], Joel H. Kramer [98], Walter A. Kukull [99], Brian W. Kunkle [25], Amanda P. Kuzma [43], Frank M. LaFerla [100], James J. Lah [101], Eric B. Larson [49,102], James B. Leverenz [103], Allan I. Levey [101], Ge Li [31,104], Andrew P. Lieberman [105], Richard B. Lipton [91], Oscar L. Lopez [90], Kathryn L. Lunetta [64], Constantine G. Lyketsos [106], John Malamon [43], Daniel C. Marson [82], Eden R. Martin [25,26], Frank Martiniuk [107], Deborah C. Mash [108], Eliezer Masliah [71,109], Richard Mayeux [20,21], Wayne C. McCormick [49], Susan M. McCurry [110], Andrew N. McDavid [45], Stefan McDonough [111], Ann C. McKee [67,97], Marsel Mesulam [29,112], Bruce L. Miller[36], Carol A. Miller [113], Joshua W. Miller [88], Thomas J. Montine [95], John C. Morris [42,114], Shubhabrata Mukherjee [49], Amanda J. Myers [46], Adam C. Naj [43], Sid O'Bryant [115], John M. Olichney [54], Joseph E. Parisi[116], Henry L. Paulson [117], Margaret A. Pericak-Vance [25,26], Elaine Peskind [104], Ronald C. Petersen [34], Aimee Pierce [50], Wayne W. Poon [55], Huntington Potter [118], Liming Qu [43], Joseph F. Quinn [93,94], Ashok Raj[44], Murray Raskind [104], Eric M. Reiman [84,119-121], Barry Reisberg [69,122], Joan S. Reisch [123], Christiane Reitz[20-22,124], John M. Ringman [48], Erik D. Roberson [82], Ekaterina Rogaeva [125], Howard J. Rosen [36], Roger N. Rosenberg [127], Donald R. Royall [128], Mark A. Sager [13], Mary Sano [40], Andrew J. Saykin [7,8], Gerard D. Schellenberg [43], Julie A. Schneider [17,19,129], Lon S. Schneider [48,130], William W. Seeley [36], Amanda G. Smith[44], Joshua A. Sonnen [95], Salvatore Spina [75], Peter St George-Hyslop [131,132], Robert A. Stern [67], Russell H. Swerdlow [38], Rudolph E. Tanzi [77], John Q. Trojanowski [133], Juan C. Troncoso [134], Debby W. Tsuang [31,104], Otto Valladares [43], Vivianna M. Van Deerlin [133], Linda J. Van Eldik [135], Badri N. Vardarajan [20-22], Harry V. Vinters [136,137], Jean Paul Vonsattel [138], Li-San Wang [43], Sandra Weintraub [28,29], Kathleen A. Welsh-Bohmer[37,139], Kirk C. Wilhelmsen [140], Jennifer Williamson [20], Thomas S. Wingo [101], Randall L. Woltjer [141], Clinton B. Wright [142], Chuang-Kuo Wu [143], Steven G. Younkin [47], Chang-En Yu [49], Lei Yu [17,19], Yi Zhao [43]

[1]Sanders-Brown Center on Aging, College of Public Health, Department of Epidemiology, University of Kentucky, Lexington, Kentucky, [2]Department of Psychiatry, University of Texas Southwestern Medical Center, Dallas, Texas, [3]Department of Neurology, Johns Hopkins University, Baltimore, Maryland,

[4]Department of Neurology, University of Michigan, Ann Arbor, Michigan, [5]Geriatric Research, Education and Clinical Center (GRECC), VA Ann Arbor Healthcare System (VAAAHS), Ann Arbor, Michigan, [6]Michigan Alzheimer Disease Center, Ann Arbor, Michigan, [7]Department of Radiology, Indiana University, Indianapolis, Indiana, [8]Department of Medical and Molecular Genetics, Indiana University, Indianapolis, Indiana, [9]Indian Alzheimer's Disease Center, Indiana University, Indianapolis, Indiana, [10]Department of Neurology, Indiana University, Indianapolis, Indiana, [11]Department of Psychiatry, University of Pennsylvania Perelman School of Medicine, Philadelphia, Pennsylvania, [12]Geriatric Research, Education and Clinical Center (GRECC), University of Wisconsin, Madison, Wisconsin, [13]Department of Medicine, University of Wisconsin, Madison, Wisconsin, [14]Wisconsin Alzheimer's Disease Research Center, Madison, Wisconsin, [1] Department of Medicine (Genetics Program), Boston University, Boston, Massachusetts, [16]Department of Pharmacology and Neuroscience, University of North Texas Health Science Center, Fort Worth, Texas, [17]Department of Neurological Sciences, Rush University Medical Center, Chicago, Illinois, [18]Department of Behavioral Sciences, Rush University Medical Center, Chicago, Illinois, [19]Rush Alzheimer's Disease Center, Rush

University Medical Center, Chicago, Illinois, [20]Taub Institute on Alzheimer's Disease and the Aging Brain, Department of Neurology, Columbia University, New York, New York, [21]Gertrude H. Sergievsky Center, Columbia University, New York, New York, [22]Department of Neurology, Columbia University, New York, New York, [23]Civin Laboratory for Neuropathology, Banner Sun Health Research Institute, Phoenix, Arizona, [24]Departments of Psychiatry, Neurology, and Psychology, University of Pittsburgh School of Medicine, Pittsburgh, Pennsylvania, [25]The John P. Hussman Institute for Human Genomics, University of Miami, Miami, Florida, [26]Dr. John T. Macdonald Foundation Department of Human Genetics, University of Miami, Miami, Florida, [27]National Alzheimer's Coordinating Center, University of Washington, Seattle, Washington, [28]Department of Pathology, Northwestern University Feinberg School of Medicine, Chicago, Illinois, [29]Cognitive Neurology and Alzheimer's Disease Center, Northwestern University Feinberg School of Medicine, Chicago, Illinois, [30]Department of Neurology, University of Washington, Seattle, Washington, [31]VA Puget Sound Health Care System/GRECC, Seattle, Washington, [32]Department of Epidemiology, Harvard School of Public Health, Boston, Massachusetts, [33]Department of Psychiatry, Massachusetts General Hospital/Harvard Medical School, Boston, Massachusetts, [34]Department of Neurology, Mayo Clinic, Rochester, Minnesota, [35]Swedish Medical Center, Seattle, Washington, [36]Department of Neurology, University of California San Francisco, San Francisco, California, [37]Department of Medicine, Duke University, Durham, North Carolina, [38]University of Kansas Alzheimer's Disease Center, University of Kansas Medical Center, Kansas City, Kansas, [39]Department of Neuroscience, Mount Sinai School of Medicine, New York, New York, [40]Department of Psychiatry, Mount Sinai School of Medicine, New York, New York, [41]Departments of Genetics and Genomic Sciences, Mount Sinai School of Medicine, New York, New York, [42]Department of Pathology and Immunology, Washington University, St. Louis, Missouri, [43]Penn Neurodegeneration Genomics Center, Department of Pathology and Laboratory Medicine, University of Pennsylvania Perelman School of Medicine, Philadelphia, Pennsylvania, [44]USF Health Byrd Alzheimer's Institute, University of South Florida, Tampa, Florida, [45]Fred Hutchinson Cancer Research Center, Seattle, Washington, [46]Department of Psychiatry and Behavioral Sciences, Miller School of Medicine, University of Miami, Miami, Florida, [47]Department of Neuroscience, Mayo Clinic, Jacksonville, Florida, [48]Department of Neurology, University of Southern California, Los Angeles, California, [49]Department of Medicine, University of Washington, Seattle, Washington, [50]Department of Neurology, University of California Irvine, Irvine, California, [51]Department of Psychiatry and Hope Center Program on Protein Aggregation and Neurodegeneration, Washington University School of Medicine, St. Louis, Missouri, [52]Program in Translational NeuroPsychiatric Genomics, Institute for the Neurosciences, Department of Neurology & Psychiatry, Brigham and Women's Hospital and Harvard Medical School, Boston, Massachusetts, [53]Program in Medical and Population Genetics, Broad Institute, Cambridge, Massachusetts, [54]Department of Neurology, University of California Davis, Sacramento, California, [55]Institute for Memory Impairments and Neurological Disorders, University of California Irvine, Irvine, California, [56]Alzheimer's Disease and Memory Disorders Center, Baylor College of Medicine, Houston, Texas, [57]Wien Center for Alzheimer's Disease and Memory Disorders, Mount Sinai Medical Center, Miami Beach, Florida, [58]Department of Neurology, Mayo Clinic, Jacksonville, Florida, [59]Rush Institute for Healthy Aging, Department of Internal Medicine,

Rush University Medical Center, Chicago, Illinois, [60]Office of Strategy and Measurement, University of North Texas Health Science Center, Fort Worth, Texas, [61]Department of Pathology, University of Alabama at Birmingham, Birmingham, Alabama, [62]Sanders-Brown Center on Aging, Department of Biostatistics, University of Kentucky, Lexington, Kentucky, [63]Department of Neurology, Indiana University, Indianapolis, Indiana, [64]Department of Biostatistics, Boston University, Boston, Massachusetts, [65]Department of Epidemiology, Boston University, Boston, Massachusetts, [66]Department of Medicine (Biomedical Genetics), Boston University, Boston, Massachusetts, [67]Department of Neurology, Boston University, Boston, Massachusetts, [68]Department of Ophthalmology, Boston University, Boston, Massachusetts, [69]Department of Psychiatry, New York University, New York, New York, [70]C.S. Kubik Laboratory for Neuropathology, Massachusetts General Hospital, Charlestown, Massachusetts, [71]Department of Neurosciences, University of California San Diego, La Jolla, California, [72]Department of Pathology and Laboratory Medicine, Emory University, Atlanta, Georgia, [73]Emory Alzheimer's Disease Center, Emory University, Atlanta, Georgia, [74]Neurogenetics Program, University of California Los Angeles, Los Angeles, California, [75]Department of Pathology and Laboratory Medicine, Indiana University, Indianapolis, Indiana, [76]Division of Genetics, Department of Medicine and Partners Center for Personalized Genetic Medicine, Brigham and Women's Hospital and Harvard Medical School, Boston, Massachusetts, [77]Department of Neurology, Massachusetts General Hospital/Harvard Medical School, Boston, Massachusetts, [78]Department of Epidemiology and Biostatistics, Case Western Reserve University, Cleveland, Ohio, [79]Center for Applied Genomics, Children's Hospital of Philadelphia, Philadelphia, Pennsylvania, [80]Department of Pathology (Neuropathology), University of Pittsburgh, Pittsburgh, Pennsylvania, [81]Institute of Neurology, University College London, Queen Square, London, United Kingdom, [82]Department of Neurology, University of Alabama at Birmingham, Birmingham, Alabama, [83]Department of Surgery, University of Texas Southwestern Medical Center, Dallas, Texas, [84]Neurogenomics Division, Translational Genomics Research Institute, Phoenix, Arizona, [85]Department of Pathology, Duke University, Durham, North Carolina, [86]Department of Genome Sciences, University of Washington, Seattle, Washington, [87]Department of Medicine (Medical Genetics), University of Washington, Seattle, Washington, [88]Department of Pathology and Laboratory Medicine, University of California Davis, Sacramento, California, [89]Department of Human Genetics, University of Pittsburgh, Pittsburgh, Pennsylvania, [90]University of Pittsburgh Alzheimer's Disease Research Center, Pittsburgh, Pennsylvania, [91]Department of Neurology, Albert Einstein College of Medicine, New York, New York, [92]Department of Biology, Brigham Young University, Provo, Utah, [93]Department of Neurology, Oregon Health & Science University, Portland, Oregon, [94]Department of Neurology, Portland Veterans Affairs Medical Center, Portland, Oregon, [95]Department of Pathology, University of Washington, Seattle, Washington, [96]Department of Pathology and Laboratory Medicine, University of California Irvine, Irvine, California, [97]Department of Pathology, Boston University, Boston, Massachusetts, [98]Department of Neuropsychology, University of California San Francisco, San Francisco, California, [99]Department of Epidemiology, University of Washington, Seattle, Washington, [100]Department of Neurobiology and Behavior, University of California Irvine, Irvine, California, [101]Department of Neurology, Emory University, Atlanta, Georgia, [102]Group Health Research Institute, Group Health, Seattle, Washington, [103]Cleveland Clinic Lou Ruvo Center for Brain Health, Cleveland

Clinic, Cleveland, Ohio, [104]Department of Psychiatry and Behavioral Sciences, University of Washington School of Medicine, Seattle, Washington, [105]Department of Pathology, University of Michigan, Ann Arbor, Michigan, [106]Department of Psychiatry, Johns Hopkins University, Baltimore, Maryland, [107]Department of Medicine - Pulmonary, New York University, New York, New York, [108]Department of Neurology, University of Miami, Miami, Florida, [109]Department of Pathology, University of California San Diego, La Jolla, California, [110]School of Nursing Northwest Research Group on Aging, University of Washington, Seattle, Washington, [111]PharmaTherapeutics Clinical Research, Pfizer Worldwide Research and Development, Cambridge, Massachusetts, [112]Department of Neurology, Northwestern University Feinberg School of Medicine, Chicago, Illinois, [113]Department of Pathology, University of Southern California, Los Angeles, California, [114]Department of Neurology, Washington University, St. Louis, Missouri, [115]Internal Medicine, Division of Geriatrics, University of North Texas Health Science Center, Fort Worth, Texas, [116]Department of Laboratory Medicine and Pathology, Mayo Clinic, Rochester, Minnesota, [117]Michigan Alzheimer's Disease Center, Department of Neurology, University of Michigan, Ann Arbor, Michigan, [118]Department of Neurology, University of Colorado School of Medicine, Aurora, Colorado, [119]Arizona Alzheimer’s Consortium, Phoenix, Arizona, [120]Banner Alzheimer's Institute, Phoenix, Arizona, [121]Department of Psychiatry, University of Arizona, Phoenix, Arizona, [122]Alzheimer's Disease Center, New York University, New York, New York, [123]Department of Clinical Sciences, University of Texas Southwestern Medical Center, Dallas, Texas, [124]Department of Epidemiology, Columbia University, New York, New York, [125] Tanz Centre for Research in Neurodegenerative Disease, University of Toronto, Toronto, Ontario, [127]Department of Neurology, University of Texas Southwestern, Dallas, Texas, [128]Departments of Psychiatry, Medicine, Family & Community Medicine, South Texas Veterans Health Administration Geriatric Research Education & Clinical Center (GRECC), UT Health Science Center at San Antonio, San Antonio, Texas, [129]Department of Pathology (Neuropathology), Rush University Medical Center, Chicago, Illinois, [130]Department of Psychiatry, University of Southern California, Los Angeles, California, [131]Tanz Centre for Research in Neurodegenerative Disease, University of Toronto, Toronto, Ontario, [132]Cambridge Institute for Medical Research and Department of Clinical Neurosciences, University of Cambridge, Cambridge, United Kingdom, [133]Department of Pathology and Laboratory Medicine, University of Pennsylvania Perelman School of Medicine, Philadelphia, Pennsylvania, [134]Department of Pathology, Johns Hopkins University, Baltimore, Maryland, [135]Sanders-Brown Center on Aging, Department of Anatomy and Neurobiology, University of Kentucky, Lexington, Kentucky, [136]Department of Neurology, University of California Los Angeles, Los Angeles, California, [137]Department of Pathology & Laboratory Medicine, University of California Los Angeles, Los Angeles, California, [138]Taub Institute on Alzheimer's Disease and the Aging Brain, Department of Pathology, Columbia University, New York, New York, [139]Department of Psychiatry & Behavioral Sciences, Duke University, Durham, North Carolina, [140]Department of Genetics, University of North Carolina Chapel Hill, Chapel Hill, North Carolina, [141]Department of Pathology, Oregon Health & Science University, Portland, Oregon, [142]Evelyn F. McKnight Brain Institute, Department of Neurology, Miller School of Medicine, University of Miami, Miami, Florida, [143]Departments of Neurology, Pharmacology & Neuroscience, Texas Tech University Health Science Center, Lubbock, Texas.
